# The genome of the blind soil-dwelling and ancestrally wingless dipluran *Campodea augens*, a key reference hexapod for studying the emergence of insect innovations

**DOI:** 10.1101/585695

**Authors:** Mosè Manni, Felipe A. Simao, Hugh M. Robertson, Marco A. Gabaglio, Robert M. Waterhouse, Bernhard Misof, Oliver Niehuis, Nikolaus U. Szucsich, Evgeny M. Zdobnov

**Affiliations:** Department of Genetic Medicine and Development, University of Geneva Medical School, and Swiss Institute of Bioinformatics, Geneva, Switzerland; Department of Entomology, University of Illinois at Urbana-Champaign, Urbana, IL, USA; Department of Ecology and Evolution, University of Lausanne, and Swiss Institute of Bioinformatics, Lausanne, Switzerland; Center for Molecular Biodiversity Research, Zoological Research Museum Alexander Koenig, Bonn, Germany; Department of Evolutionary Biology and Ecology, Albert Ludwig University, Institute of Biology I (Zoology), Freiburg, Germany; Natural History Museum Vienna, 3^rd^ Zoological Dept., Vienna, Austria

**Keywords:** Two-pronged bristletails, Entognatha, chemosensory genes, ionotropic receptors, gustatory receptors, photoreceptors

## Abstract

The dipluran two-pronged bristletail *Campodea augens* is a blind ancestrally wingless hexapod with the remarkable capacity to regenerate lost body appendages such as its long antennae. As sister group to Insecta (*sensu stricto*), Diplura are key to understanding the early evolution of hexapods and the origin and evolution of insects. Here we report the 1.2-Gbp draft genome of *C. augens* and results from comparative genomic analyses with other arthropods. In *C. augens* we uncovered the largest chemosensory gene repertoire of ionotropic receptors in the animal kingdom, a massive expansion which might compensate for the loss of vision. We found a paucity of photoreceptor genes mirroring at the genomic level the secondary loss of an ancestral external photoreceptor organ. Expansions of detoxification and carbohydrate metabolism gene families might reflect adaptations for foraging behaviour, and duplicated apoptotic genes might underlie its high regenerative potential.

The *C. augens* genome represents one of the key references for studying the emergence of genomic innovations in insects, the most diverse animal group, and opens up novel opportunities to study the under-explored biology of diplurans.

## Introduction

Diplura, also referred to as two-pronged bristletails, are ancestrally wingless hexapods. With a worldwide distribution, they constitute a ubiquitous component of soil ecosystems, occurring primarily in the litter layer and humid soils of forests and grasslands, but even colonising deeper soil layers (Orgiazzi et al. 2016). Their soft, mostly unpigmented and weakly sclerotised elongated bodies range up to ten millimetres (Sendra et al. 2018). All diplurans are blind and possess long myocerate (bead-like) antennae, a pair of filamentous terminal appendages (cerci), as well as remnants of abdominal legs (styli) (Grimaldi 2010). Larval development is epimorphic (*i.e.*, body segmentation is completed before hatching), and moulting continues during the entire life span (Koch 2009), likely playing an important role in their remarkable capacity to regenerate lost body appendages (Orgiazzi et al. 2016; B.K.Tyagi & Veer 2016; Maruzzo et al. 2005; Whalen & Sampedro 2010). Fertilisation of the diplurans’ eggs is indirect: males produce and deposit spermatophores (sperm packets) that are subsequently collected by conspecific females (Koch 2009). Approximately 1,000 species of Diplura have been described and divided into three main groups: Campodeoidea with long segmented and filiform cerci, Japygoidea with unsegmented pincer-like cerci, and Projapygoidea with short cerci bearing spinning glands (Koch 2009). While the latter two are predators and eat small arthropods, Campodeoidea, of which *Campodea augens* is a typical representative, are omnivores and are part of the decomposer soil community (Carpenter 1988; Lock et al. 2010). One distinctive feature of diplurans respect to insects (Insecta *sensu stricto*) is the position of the mouthparts which are hidden within head pouches (entognathous) (Böhm et al. 2012) like in Collembola (springtails) and Protura (coneheads), while in insects the mouthparts are exposed (ectognathous). However, many features present in diplurans have been retained in insects (*sensu stricto)*, and phylogenetic and morphological studies suggested that Diplura likely represent the sister group of insects (Misof et al. 2014; Machida 2006; Sasaki et al. 2013), making Diplura a crucial reference taxon when studying the early evolution of insect genomes. Despite their evolutionary importance, diplurans have remained under-explored in particular at the genomic level, hampering a deeper understanding of the early evolution of hexapod genomes. Therefore we sequenced and annotated the genome of *C. augens*, and present here the first analysis of a dipluran genome. For some of our analyses we got permission to include another dipluran genome (*Catajapyx aquilonaris*) representing the lineage Japygoidea, which have been sequenced in the context of the i5k initiative (https://i5k.nal.usda.gov/). By analysing and comparing *C. augens* genome with those of twelve other arthropods we found evidence for rapid gene family evolution in *C. augens*. For example, we uncovered a massive expansion of the ionotropic receptor gene family, which might compensate for the loss of vision. We also described expansions of gene families related to detoxification and carbohydrate metabolism, which might reflect adaptations of *C. augens*’ foraging behaviour, and duplications of apoptotic genes, which could underlie its regenerative potential. Intriguingly, in the genome we discovered putative remnants of endogenous viral elements resembling negative single-stranded RNA viruses of the Orthomoxyviridae, a viral family that includes the Influenza viruses. With Diplura representing the sister clade to Insecta, the genome of *C. augens* serves as a key out-group reference for studying the emergence of insect innovations, such as the insect chemosensory system, and opens up novel opportunities to study the under-explored biology of diplurans.

## Results and discussion

### The *Campodea augens* genome

We assembled the genome of the dipluran *Campodea augens* (Figure 1 and Supplementary Figure 1) using 120 Gb of Illumina paired-end reads data, representing ca. 100x coverage of the genome, and additional 90 Gb of mate-pair reads (Supplementary Figure 2 and Supplementary Table 1). The inferred draft assembly spans 1.13 Gbp, which is close to the genome size of 1.2 Gbp estimated via flow cytometry, indicating that most of the genome is likely present in the assembly (Supplementary Note and Supplementary Table 2). The high level of completeness was also confirmed with BUSCO (Waterhouse et al. 2018) as detailed below. Although genomes of many hexapod groups have recently been sequenced, only few are from ancestrally wingless hexapods: four collembolans (*Folsomia candida*, *Orchesella cincta, Holacanthella duospinosa* and *Sinella curviseta*) (Faddeeva-Vakhrusheva et al. 2017, 2016; Wu et al. 2017; Zhang et al.), whose genomes have been published; and a dipluran (*Catajapyx aquilonaris*) which have been sequenced in the context of the i5k initiative (https://i5k.nal.usda.gov/). Currently, there are no sequenced genomes of proturans.

**Fig. 1.**
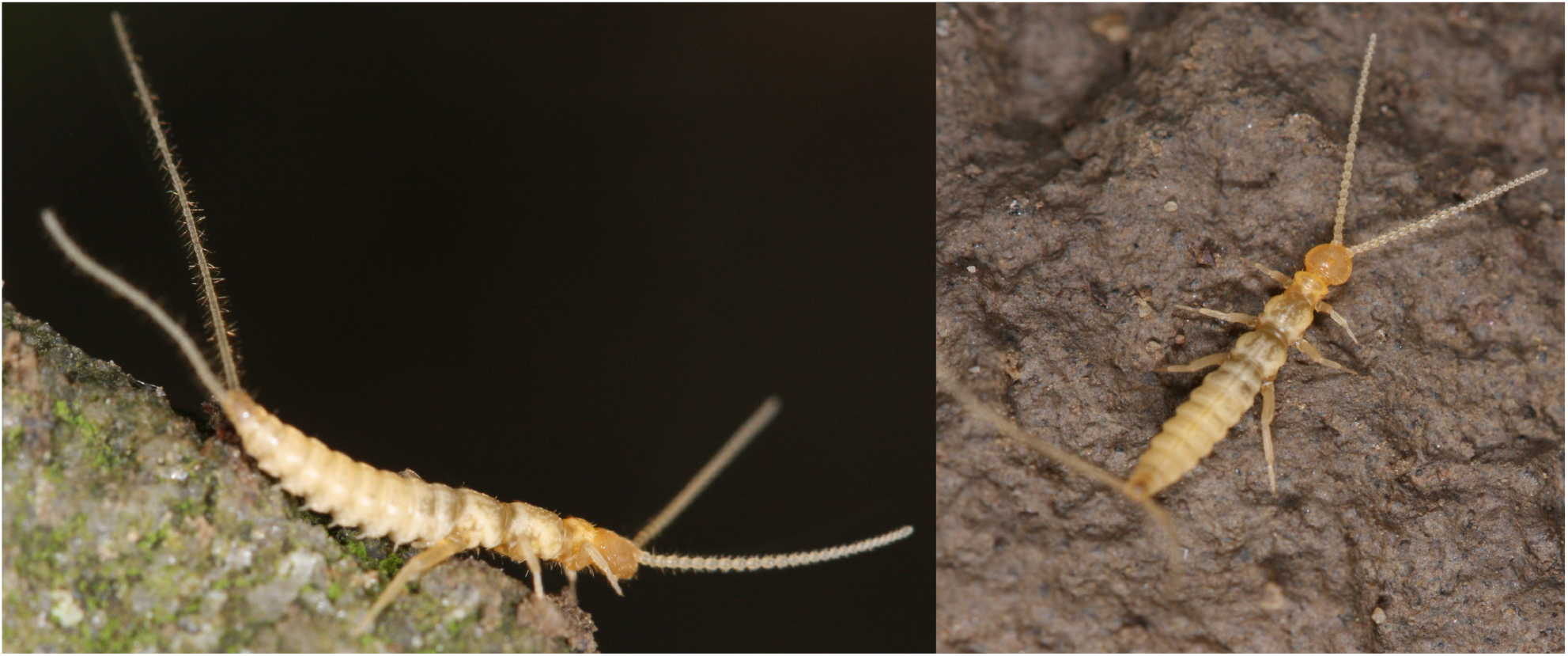
Lateral and dorsal view of *Campodea augens*.

Genomes of ancestrally wingless hexapods have proven difficult to sequence, as they are often large (as in the case of *C. augens*), and because many species are typically difficult to rear. Thus, it is almost impossible to reduce natural heterozygosity (e.g., via inbreeding experiments), which potentially compromises genome assembly efforts (Richards & Murali 2015). Our *C. augens* assembly is much larger than the assemblies of *C. aquilonaris* (312 Mbp) and of the four collembolans (221–381Mbp) (Supplementary Fig. 3 and Supplementary Table 2). The large genome size of *C. augens* is likely explained by the proliferation of repeats, which account for ca. 46% of the assembly (Supplementary Table 3). Most of these repeats are unclassified and represent 33% (376 Mb) of the assembly. Among the identified repetitive elements, the *hAT* superfamily is the most abundant DNA transposon and accounts for 5.9% of the genome sequence (Supplementary Table 3). The high level of repeats posed a challenge for genome assembly, resulting in a total of 18,765 scaffolds with N_50_ values for contigs and scaffolds at 33 Kbp and 235 Kbp, respectively (Supplementary Fig. 3 and Supplementary Table 2). The annotated gene set of *C. augens* comprises 23,978 predicted protein-coding genes and exceeds in number that of *C. aquilonaris* (10,901), however it is comparable to that of *F. candida* (22,100) and that of *O. cincta* (20,247) (Supplementary Table 4). More than 90% of the gene models have support from *C. augens* transcripts. Approximately 74% of the predicted proteins show a significant amino acid similarity (BLASTP e-value < 1e-10) to existing entries in the Uniref50 database, while ca. 95% of the proteins contain a domain assigned with InterProScan, providing high confidence in the accuracy of annotations. Assessments with Benchmarking Universal Single-Copy Orthologs (BUSCOs) (Waterhouse et al. 2018) identified 97.0–97.6% (90.4–91.8% complete) of the 1,066 BUSCOs expected to be conserved in arthropods, indicating high completeness of the assembly and gene models (Supplementary Fig. 4 and Supplementary Table 5). The completeness of the gene set is additionally indicated by the successful identification of all *Hox* genes known from *F. candida* (*lab, hox3, pb, ftz, src, dfd, antp, ubx, abd-A, abd-B*). In *C. augens*, however, these genes are distributed across five scaffolds (Supplementary Figure 5) spanning more than 1 Mbp.

The number of introns/exons per gene and exon lengths in *C. augens* are similar to those of *C. aquilonaris* and those of collembolans (Supplementary Table 4). However, the median size of the gene models in *C. augens* is roughly four times longer than in *C. aquilonaris* and three to five times longer than in collembolans (Supplementary Table 4 and Supplementary Figure 6). This is mostly due to longer introns in the genome of *C. augens* (Figure 2 and Supplementary Table 4 and Supplementary Figure 6). Indeed, introns remarkably span 27% of the *C. augens* genome, accounting for a total of 306 Mbp, which is equivalent in size to the entire *C. aquilonaris* genome. It has been proposed that longer introns are associated with an increased metabolic cost (Vinogradov 1999), so strong selection against long introns is expected. Various factors could have promoted the evolution of large intron sizes in *C. augens*, ranging from drift to low genome-wide recombination rates, and remain to be studied. The differences in gene content, genome size, and intron sizes between C*. augens* and *C. aquilonaris* suggest that these two dipluran genomes have differentiated greatly from each other, likely through the spread of repetitive elements in *C. augens*.

**Fig. 2.**
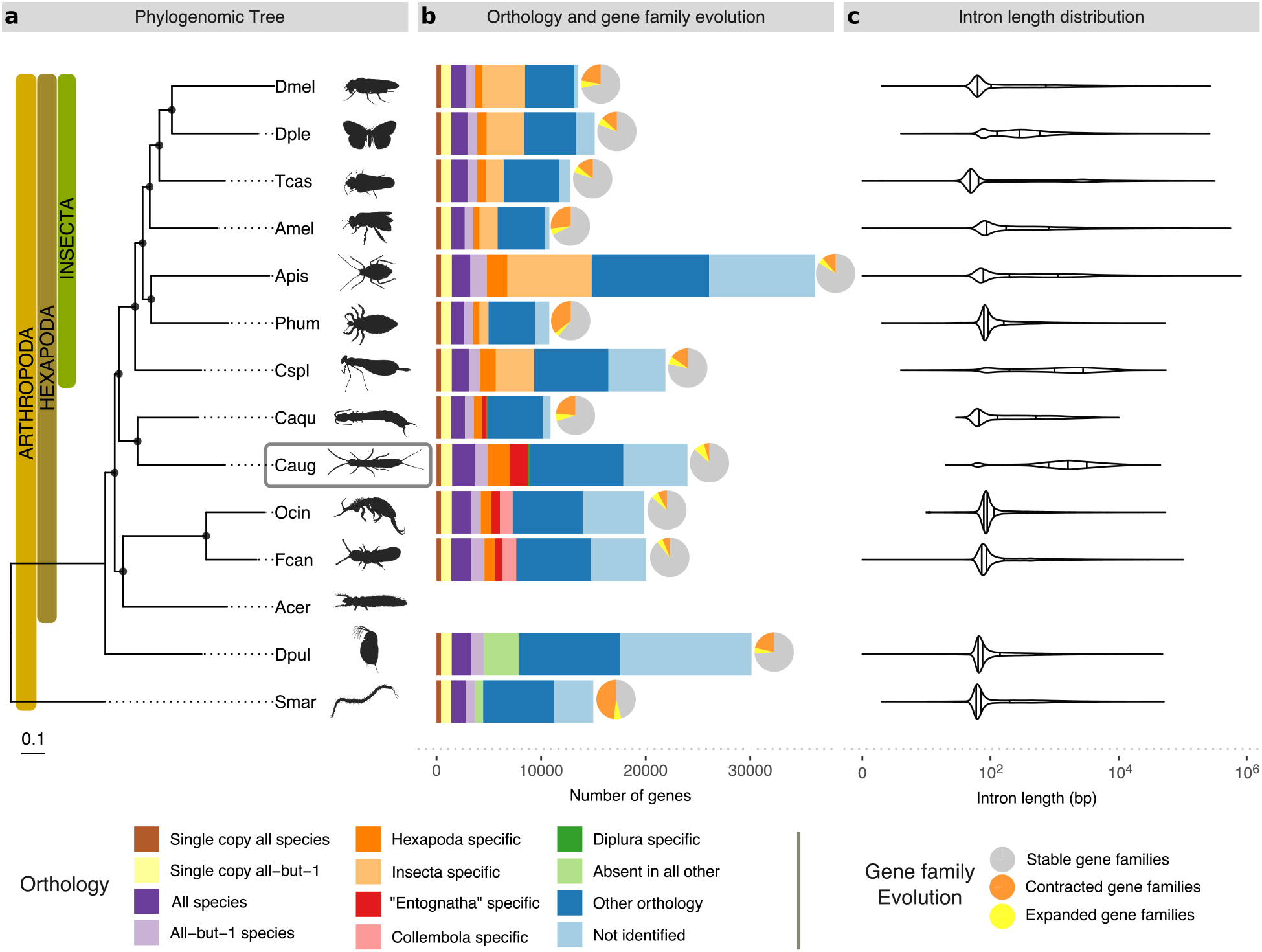
Species phylogeny, orthology, gene family expansions/contractions and intron length distributions of *Campodea augens* and 13 other arthropod species. **a**, Phylogenomic tree of 14 arthropod species, including ancestrally wingless hexapods: two diplurans, two collembolans and one proturan (data for proturan were obtained from transcriptome). The maximum likelihood phylogeny was estimated from the aligned protein sequences of 371 single-copy orthologs, using the centipede, *Strigamia maritima*, as the outgroup. Branch lengths represent substitutions per site. Black circles on nodes indicate bootstrap support > 0.95. Dmel, *Drosophila melanogaster* (fruit fly); Dple, *Danaus plexippus* (monarch butterfly); Tcas, *Tribolium castaneum* (red flour beetle); Amel, *Apis mellifera* (honey bee); Phum, *Pediculus humanus* (body louse); Apis, *Acyrthosiphon pisum* (pea aphid); Cspl, *Calopteryx splendens* (banded demoiselle); Caqu, *Catajapyx aquilonaris* (northern forcepstail); Acer, *Acerentomon* sp. (coneheads); Fcan, *Folsomia candida* (springtail); Ocin, *Orchesella cincta* (springtail); Dpul, *Daphnia pulex* (water flea); Smar, *Strigamia marítima* (centipede). **b**, Bars: total gene counts per species grouped into the following categories: single-copy orthologs in all 13 species or in all except one species, present in all or all except one species allowing for gene duplications, lineage-specific orthologs (Hexapoda, “Entognatha”, Collembola, Diplura, and Insecta), genes present in only one species (absent in other species), genes with all other ortholog relationships, and with no identifiable orthologs; pie charts: proportions of species-specific gene family expansions (yellow), contractions (orange) and stable gene families (grey) as estimated using CAFE (these proportions include the number of non-significant expansions/contractions, see text and Supplementary Figure 8 for more details). **c**, Violin plots of intron length distributions of the corresponding species in the phylogeny. Vertical bars correspond to median, lower and upper quartiles. Silhouette images were obtained from PhyloPic (http://phylopic.org). All are under public domain except: *Calopteryx splendens* by Maxime Dahirel; Protura, *Orchesella* springtail and *Strigamia* centipede by Birgit Lang; all licensed under the Creative Commons Attribution 3.0 Unported license (http://creativecommons.org/licenses/by/3.0/); and *Acyrthosiphon pisum* by Joseph Hughes licensed under Creative Commons Attribution NonCommercial-ShareAlike 3.0 Unported licence (https://creativecommons.org/licenses/by-nc-sa/3.0/).

### Phylogenomics and orthology

Although the monophyly of Hexapoda is now well supported (Misof et al. 2014; Grimaldi 2010; Sasaki et al. 2013), the relationships among Protura, Collembola, and Diplura (traditionally grouped as “Entognatha”) and their placements with respect to ectognathous hexapods (Insecta) have been much debated. The following hypotheses exist regarding the phylogenetic position of Diplura: *i*) Diplura as sister group of Protura and Collembola (Entognatha hypothesis) (von Reumont et al. 2009); *ii*) Diplura as sister group of Protura (Nonoculata hypothesis) (Andrew 2011; Luan et al. 2005; Meusemann et al. 2010); and *iii*) Diplura as sister group of Insecta *s. str.* (Kukalová-Peck 1998; Misof et al. 2014), with “Entognatha” being paraphyletic. While the fossil record of diplurans is poor, with the oldest known fossil from Lower Cretaceous deposits of Brazil (Wilson & Martill 2001), they probably originated in the Early Devonian (Misof et al. 2014). Phylogenomic analysis employing a supermatrix of 358 concatenated single-copy protein-coding genes of 14 arthropods support the idea of “Enthognatha” being paraphyletic and Diplura representing the sister group of Insecta (Figure 2, Panel a). This result is in agreement with previous phylogenetic studies that analysed phylogenomic data (Misof et al. 2014), Sanger-sequenced nuclear-encoded protein-coding genes (Regier et al. 2004), or studied fossil (Kukalová-Peck 1987), morphological (Giribet et al. 2004) and comparative embryological (Ikeda & Machida 2001; Tomizuka & Machida 2015) evidence. However, more extensive sampling of genomes of Collembola, Diplura and Protura is desirable to improve confidence in the inferred phylogenetic relationships.

Results from analysing gene orthology via clustering the *C. augens* gene set with genes of 169 other arthropods are available from OrthoDB v10 (Kriventseva et al. 2019) (https://www.orthodb.org/). Analysis of the full set of orthologous groups identified 19,063 *C. augens* genes that have orthologs in at least one other arthropod genome included in OrthoDB. Of these, 17,871 genes have orthologs in at least one of the 12 genomes under consideration in Figure 2. In contrast, 4,929 *C. augens* genes have no identifiable orthology relationships with genes of other arthropods (and 6,121 genes when only considering species in Figure 2), and similarly for the two collembolans. The median protein length of these predicted genes is 209 amino acids, and some are likely to be gene fragments. 3008 and 3896 of them have at least one hit to a PFAM domain and at least one hit to another protein from the rest of the classified *C. augens* proteome (e-value < 1e-05), respectively. Sequencing the genomes of additional ancestrally wingless hexapods and closely related outgroups (e.g. Remipedia) is required to reveal what fraction of these genes is indeed unique to *C. augens* or Diplura, or whether they evolved earlier in the history of Arthropoda.

### Gene family evolution

To examine in detail the most dramatic changes in the gene repertoires among ancestrally wingless hexapods, we modelled gene family evolutionary dynamics using the Computational Analysis of gene Family Evolution (CAFE) approach (Han et al. 2013). We detected significant (*P* < 0.01) gene copy-number changes in orthologous groups on both the branches leading to Diplura (19 expansions and zero contractions) and to Collembola (45 expansions and seven contractions) (Supplementary Figure 8 and Supplementary Table 8). The number of species-specific significant expansions and contractions varied within ancestrally wingless hexapods (*F. candida*: +49/-12; *O. cincta*: +28/-9; *C. aquilonaris*: +6/-30), with *C. augens* displaying the largest number of expansions and the smallest number of contractions (+90/-1) (Figure 2, Panel b). Among the expanded gene families in *C. augens* there are families of chemosensory receptors, specifically gustatory (GRs) and ionotropic receptors (IRs). A detailed analysis of these is presented below, where we report a massive expansion of IR gene family that accounts for ca. 8% of the entire gene set content. Other expanded gene families detected with CAFE are involved in detoxification and sugar metabolism/transport, which may be related to the foraging ecology of *C. augens*, and in apoptosis, which may play a role in the high regeneration capacity of *C. augens* (Supplementary Tables 8 and 9 and Supplementary Figure 9).

### The gene repertoire of *C. augens* mirrors the secondary loss of an ancestral external photoreceptor organ

The lack of external eyes and ocelli (simple eyes) in diplurans is presumably a consequence of degenerative processes of tissues that are functionless in dark soil environments. The eyeless collembolan *F. candida* detects both UV and white light (Fox et al. 2007; Gallardo Ruiz et al. 2017), and it has been hypothesised that the light avoidance behaviour of *F. candida* is mediated by non-ocular photoreceptors (Fox et al. 2007). Campodeids also avoid light and may sense it over their body surfaces (George 1963), having a thin and weakly sclerotised cuticle. A similar mechanism of internal photoreceptor cells or organelles might apply to *C. augens*. This kind of light-sensitivity might be sufficient for organisms that require light perception, but not spatial light resolution. However, it cannot be excluded that other environmental triggers, such as air temperature or relative humidity, play a role in the observed light-avoidance behaviour of *C. augens*. (George 1963). We searched the *C. augens* gene set for known photoreceptor genes, such as opsins, which in complex with molecular chromophores are sensitive to specific wavelengths of light. Although we found 149 G-protein-coupled receptor rhodopsin-like domains (Pfam PF00001), none of the corresponding proteins belong to the opsin subfamily. Using both profile hidden Markov models (HMMs) of known opsins and TBLASTN searches of the genome, we found two fragments of putative opsins with best-hits to ultraviolet-sensitive opsins. One fragment contains the opsin retinal binding site (prosite identifier: PS00238). Whether the opsin fragments are part of functional genes and play a role in the reported light avoidance behaviour of diplurans awaits further investigation. In flies, the Gr28b set of receptors includes some involved in perception of light and heat (Xiang et al. 2010; Ni et al. 2013), and it is possible that in the absence of functional opsins, relatives of these GRs might confer light sensitivity to *Campodea* and other diplurans. We did not identify clear relatives of the DmGr28a/b lineage in the genomes of *C. augens* or that of *C. aquilonaris* (Figure 3), however, it is possible that other GRs have evolved this function in diplurans. Furthermore, we did not find DNA photolyases, which would repair DNA damage induced by UV light (Supplementary Table S10), or cryptochrome proteins, which are components of the arthropod central circadian clock. DNA photolyases or cryptochromes were not found in the centipede *Strigamia maritima* (Chipman et al. 2014) and appear to be absent also in the dipluran *C. aquilonaris* and the collembolan *F. candida*, all three of which are eyeless. Intriguingly, the collembolan *O. cincta*, which possesses simple eyes, has DNA photolyases (Supplementary Table S10). The loss of cryptochromes and DNA photolyases thus appears to be a common feature among blind arthropods with a subterranean lifestyle. With the paucity of photoreceptor genes, the gene repertoire of *C. augens* thus mirrors at the genomic level the secondary loss of the ancestral external photoreceptor organ. The absence of cryptochromes also calls into question the presence of the canonical circadian clock system, which in most organisms regulates various behaviours, biochemical and physiological functions, and which is set by light. Searches for components of the major regulatory feedback loop of the arthropod circadian clock (e.g., *clock*, *cycle*, *jetlag, period*, *timeless*), revealed only partial homologs of *timeless* and *clock*. The absence of cryptochromes and of other known circadian clock genes such as *cycle*, *jetlag* and *period* may indicate that *C. augens* has not maintained canonical circadian rhythms and thus possibly retained light-sensitivity primarily for orientation and habitat choice.

**Fig. 3.**
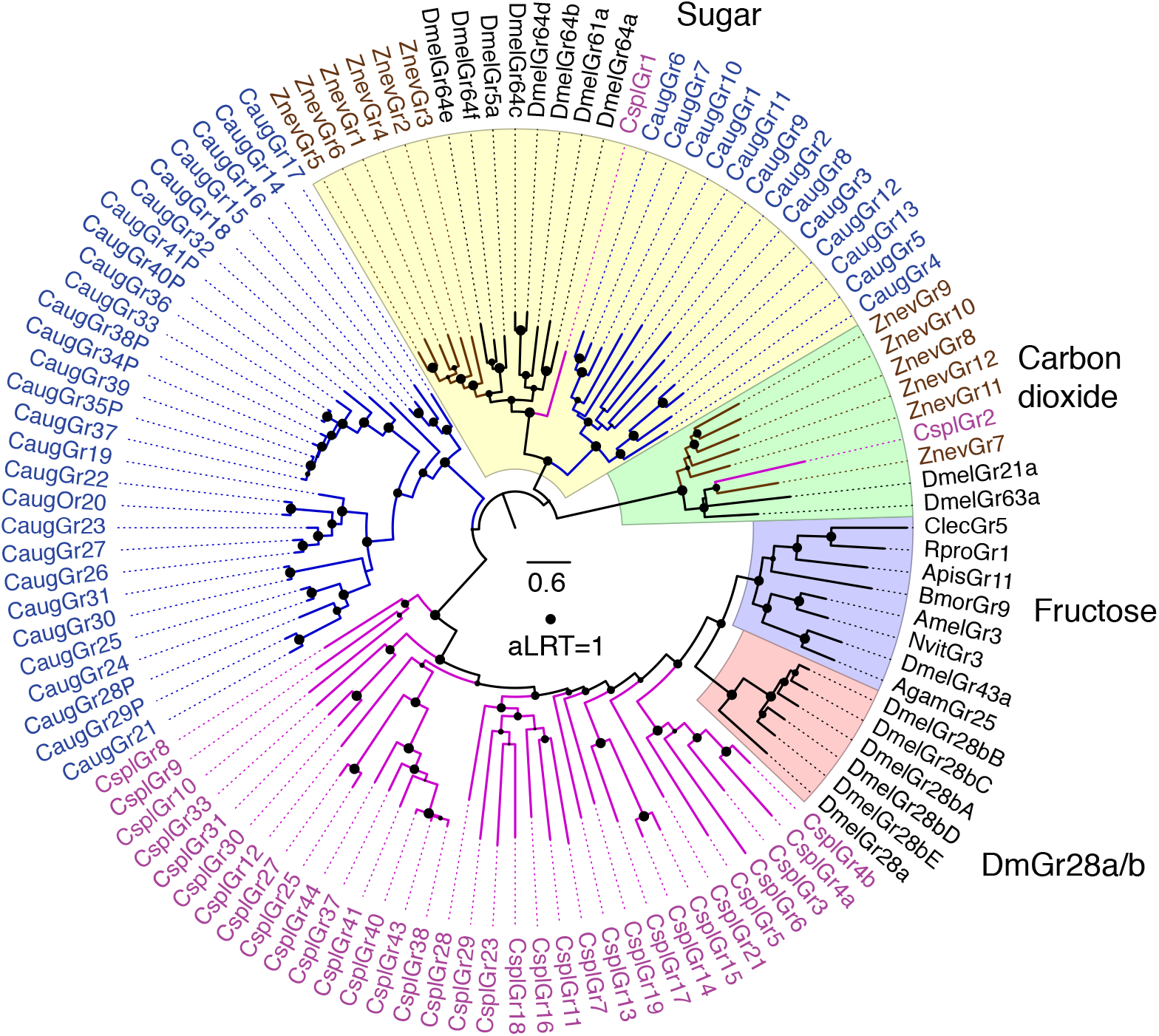
Phylogenetic relationships of the *C. augens* GRs with representatives from other insects. The sugar and carbon dioxide receptor clades were defined as the outgroup based on their location in trees of the entire animal GR/GRL family (Robertson 2015). Major subfamilies or lineages are highlighted by colours and indicated outside the circle. *C. augens* GRs are coloured blue. Representative GRs from other insects include the damselfly *C. splendens* (Cspl – purple) (Ioannidis et al. 2017), dampwood termite *Z. nevadensis* (Znev – brown) (Terrapon et al. 2014) and selected endopterygotes in black. The scale bar indicates substitutions per site and the size of the filled circles on nodes indicates approximate Likelihood Ratio Tests (aLRT) from PhyML v3.0 from 0 to 1.

### GRs and the massive expansion of the IR family might compensate for the loss of vision

Antennae are one of the most important olfactory organs of insects, and *C. augens* possesses one pair of long myocerate (bead-like) antennae that contain muscles protruding from the head and innervated on each side, allowing controlled movements of the antennae (Böhm et al. 2012). In the central nervous system, *C. augens* possesses two mushroom bodies and two different types of putative olfactory glomeruli: four to five large and elongated ventral glomeruli and a set of small spheroidal dorsal glomeruli (Böhm et al. 2012) innervated from the antennae. These anatomical features suggest an advanced olfactory system in Diplura (Böhm et al. 2012). In fact, being blind, diplurans like *C. augens* must depend largely on their senses of smell, taste and touch to negotiate their environments, as indicated by their continual use of their long antennae while moving around. Given the likely sister group relationship to Insecta, Diplura represent a crucial taxon to understand the evolutionary origin of the chemosensory system in insects. However, studies of the genomic chemosensory amenities of Diplura are scarce. Brand et al. (2018) noted that the sequenced genome of the dipluran *C. aquilonaris* contains no odorant receptor (OR) gene family members. We likewise were unable to identify ORs or the Odorant receptor co-receptor (Orco) in the *C. augens* genome. Along with the absence of ORs in three collembolan genomes (Wu et al. 2017), this observation corroborates the conclusion by Brand et al. (2018) that the OR gene family evolved in the stem lineage of Insecta. Instead, manual curation of the GR and IR gene families highlighted by the CAFE and domain analyses identified a modestly sized family of 41 GRs and a hugely expanded family of 2431 IRs. This large repertoire of chemosensory genes likely provides *C. augens* with a sophisticated chemosensory system. Seven of the GRs are clear pseudogenes, and several gene fragments indicate additional pseudogene remnants. The most prominent subfamilies of arthropod GRs are the sugar receptors, which are also present in crustaceans like *Daphnia pulex* (Peñalva-Arana et al. 2009) and *Hyalella azteca* (Poynton et al. 2018), and the carbon dioxide receptors (and their relatives), which have been found in early divergent insect lineages, such as Odonata (Ioannidis et al. 2017; Robertson 2019). *C. augens* has 13 candidate sugar receptors clustering with insect representatives of this subfamily (Figure 3) and sharing a glutamic acid immediately after the TY in the conserved TYhhhhhQF motif in the seventh transmembrane domain (where h is usually a hydrophobic amino acid). In contrast, there is no evidence for the presence of the carbon dioxide receptor subfamily or of a distinctive lineage of fructose receptors, both found in insect genomes (Figure 3). The remaining 28 GRs cluster together, with no close relatives in insects and by analogy with those of insects might include receptors for various “bitter” plant compounds and for cuticular hydrocarbons (Robertson 2019). The remarkably large set of 2,431 *C. augens* IRs contains representatives of each of the three conserved co-receptors, named after their *Drosophila melanogaster* orthologs (Ir8a, 25a, and 76b), with duplications resulting in four copies (paralogs) of Ir25a (we also detected fragments of a possibly fifth copy). Most insects have an additional set of single-copy IRs (Ir21a, 40a, 68a, and 93a) involved in the perception of temperature and humidity (Knecht et al. 2017), but only Ir93a was identified in *C. augens*. Intriguingly, Ir93a is also the only one of these four IRs to have been identified in several other arthropods groups, like ticks, mites, centipedes, and crustaceans (Robertson 2019). However, the copepod *Eurytemora affinis* seems to encode a distant relative of Ir21 (Eyun et al. 2017; Poynton et al. 2018), implying that Ir21a was already present in the stem group of Hexapoda and has likely been lost from *C. augens* and other arthropods. No relatives of other IR lineages that are widely present in insects, for example the Ir75 clade involved in perception of diverse acids (Prieto-Godino et al. 2017), seem to be present in the *C. augens* genome (Figure 4 and Supplementary Figure 10). On the other hand, we found an extraordinary expansion of 2,424 mostly intronless IR genes, named from Ir101 to Ir2525, 901 of which are pseudogenes (defined as encoding at least 50% length of related IRs, there being innumerable additional shorter gene fragments). The vast majority of these genes are intronless in their coding regions, however there are several lineages with one or two introns interrupting the coding region, all apparently gained independently from intronless ancestors, something previously observed in several insects, e.g. the cockroach *Blatella germanica* (Robertson et al. 2018) in which the IR family expansion clearly represents an independent event from the expansion observed in *C. augens*. Thus Ir101-118 have a phase-2 intron about 1/3 into the coding region, although the divergent first exon was not identified for some of them. Divergent subfamilies of these IRs were discovered by weak matches in TBLASTN searches, so additional intron-containing genes were only detected towards the end of this iterative search process (because matches are weaker when the coding region is split by an intron). Thus Ir2291-2334 also have a phase-2 intron roughly 1/3 along. Ir2335-2371 have a phase-0 intron near the middle of the coding region, as do Ir2372-2413, but in a different location roughly 1/3 along. Finally, Ir2497-2523 have a phase-0 intron followed by a phase-2 intron. All of these six introns are not only in two different phases and are in different locations in the coding sequences, but as shown in Figure S10 each is in a distinct gene lineage, so these six introns have been gained independently, and somewhat surprisingly have never been lost from any genes within each of these lineages. These IR genes exhibit many of the traits common to highly-expanded gene families like these chemoreceptors in other animals, including some large arrays, albeit commonly with genes on both strands, implying considerable short-range inversions from an original tandem-orientation resulting from duplication by unequal crossing-over. Occasionally a complicated pattern was noticed in which several scaffolds had comparable suites of otherwise distantly-related genes in them, suggesting that segmental duplications have also played a part in the expansion of this gene family and hence the genome. Nevertheless, vast numbers are present as singletons in a scaffold, or far removed from others in a large scaffold, so there has been enormous genomic flux in this genome moving even individual genes around. These intronless and divergent IRs consist of numerous gene lineage expansions, many of which are very recent, as indicated by short branches, and are replete with pseudogenes and fragments, indicating rapid gene family turnover of the IRs in *C. augens* (Figure 4 and Supplementary Figure 10). Large gene families typically have numerous pseudogenes and indeed at least 901 (37%) are pseudogenes, and as noted above there are innumerable apparently pseudogenic fragments. This is near the top end for percentage of chemoreceptor pseudogenes in arthropods (Robertson 2019), but less than some mammals, with over 50% pseudogenes being common (Niimura 2014). These pseudogenes are distributed throughout the family (Figure S10), although there is a tendency for the most highly and recently expanded lineages to have the highest frequencies of pseudogenes, for example in the bottom half of this circular tree. This is visually conspicuous in the tree because these pseudogene names have a P at the end, making them longer than the related intact gene names. The numbers of obvious pseudogenizing mutations (stop codons, frameshifts, and large deletions) were noted for each pseudogene and a histogram (Figure S11) reveals that the majority have single mutations, a pattern seen previously for the cockroach IRs (Robertson et al. 2018), consistent with the young nature of most of these pseudogenes in recent gene lineage expansions. Presumably, most older pseudogenes also eventually suffer major deletions that reduce them to the innumerable fragments observed. The total of 2472 chemoreceptor genes is comparable to the largest numbers seen in some mammals, such as 2294 ORs in the cow, 2658 in the horse, and 4267 in the elephant (Niimura 2014), although these counts do not include their vomeronasal or taste receptors, and is far larger than any chemoreceptor repertoire known in other arthropods, the two highest so far being 825 in the spider mite *Tetranychus urticae* (Ngoc et al. 2016) and 1576 in *B. germanica* (Robertson et al. 2018). This remarkable expansion of the IR family exceeds even those found in the genomes of cockroaches with 640 and 897 (Li et al. 2018; Robertson et al. 2018), making this the largest IR family known in animals. This massive expansion of the IR family of chemoreceptors in this genome presumably reflects the dependence of this dipluran on its chemical senses of both smell and taste.

**Fig. 4.**
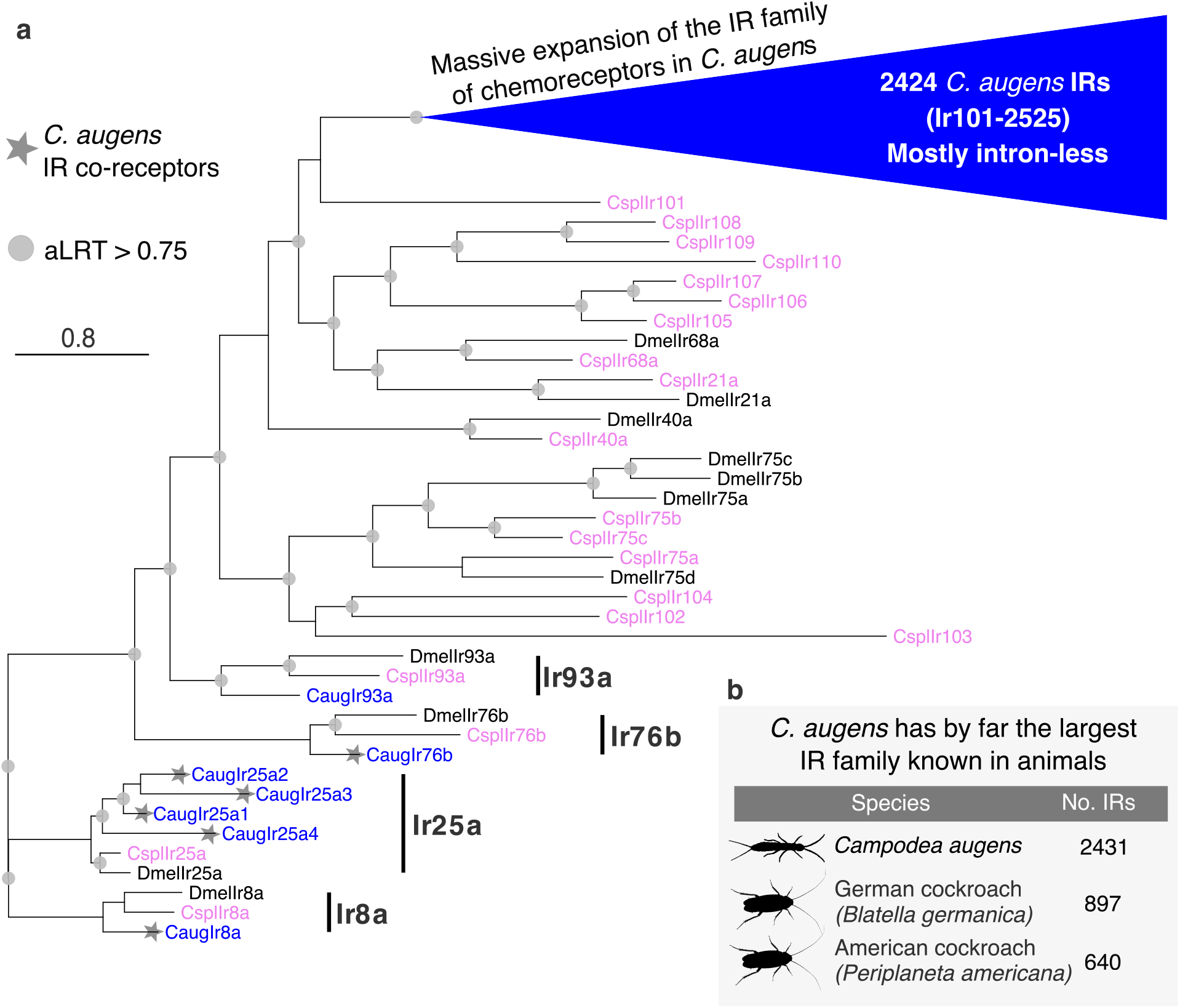
Phylogenetic tree of the *C. augens* IR family. **a**, The tree was rooted by declaring the Ir8a and 25a lineages as the outgroup, based on their basal positions within larger trees including the ionotropic glutamate receptors from which the IRs evolved. The blue triangle represents the massive expansion of *C. augens* IR family. Most of these 2424 IRs are intron-less, except for five lineages that have idiosyncratically gained introns (See Supplementary Fig. 10 for details). The *Campodea augens* (Caug) proteins are in blue, the *Drosophila melanogaster* (Dmel) proteins for the seven conserved IRs with orthologs in *C. augens*, as well as the Ir75 clade, are coloured black, while the *Calopteryx splendens* (Cspl) proteins are coloured purple. The four conserved lineages are marked with a black bar. *C. augens* IR co-receptors are marked with a star. The scale bar indicates substitutions per site. Grey circles indicate nodes with an approximate Likelihood-Ratio Test (aLRT) > 0.75. **b**, Table displaying the animal species with the largest repertoire of IRs. *C. augens* possesses by far the largest IR family known in animals to date. Silhouette images were obtained from PhyloPic (http://phylopic.org). All are under public domain.

### Expansion of gene families related to xenobiotic detoxification and apoptosis

Analysis of the gene sets revealed that *C. augens* possesses the largest set of enzymes related to detoxification (808 genes) among the 13 species examined including representatives of main hexapod orders (Table 1). These families were found to be expanded and include ATP-binding cassette transporters (ABC transporters), carboxylesterases (CEs), cytochrome P450s (CYPs), UDP-glucoronosyl/glucosyl transferases (UGTs), and Glutathione S-transferases (GSTs) (Supplementary Tables 8 and 9). *C. augens* has the largest set of ABC transporters, 290 in total, thus 1.5 times as many as the second largest set of ABC transporters identified in the collembolan *F. candida* (193 in total). Likewise, carboxylesterases are more abundant in *C. augens* (147) with repertoires in collembolans being nearly as large (*F. candida*, 120 genes; *O. cincta*, 110 genes). Cytochrome P450 domains in *C. augens* (202 in total) are similar in number to those in collembolans (*F. candida*, 214; *O. cincta*, 260), but they are much higher than in all other arthropods examined. GST counts are within the ranges observed in the other species, while UGTs are again most abundant in *C. augens* (104 in total), double to those in collembolans (51 and 57, respectively) and ten times as many than in the closely related dipluran *C. aquilonaris* (eight in total). Several of these genes are clustered in the *C. augens* genome, for example sets of five P450s and four GSTs on scaffold_1992 and a set of six UGTs on scaffold_1829 (Supplementary Figure 11). As hypothesised for the two collembolans (Faddeeva-Vakhrusheva et al. 2016, 2017), the large repertoire of detoxification enzymes in *C. augens* might reflect adaptations to living in soil environments. The strikingly lower counts of detoxification genes in *C. aquilonaris* (212 in total), which also lives in the soil, may reflect its feeding behaviour, which is different from that of *C. augens*. While *C. augens* and the two collembolans are herbivorous or detritivorous, *C. aquilonaris* is a predator, feeding on small arthropods. Thus, *C. aquilonaris* might not have the same detoxification needs as *C. augens*. The latter likely has to cope with ingesting xenobiotic compounds which persist in decaying organic matter, such as plant anti-herbivory toxins, lignocellulose by-products and feeding deterrents.

**Table 1.**
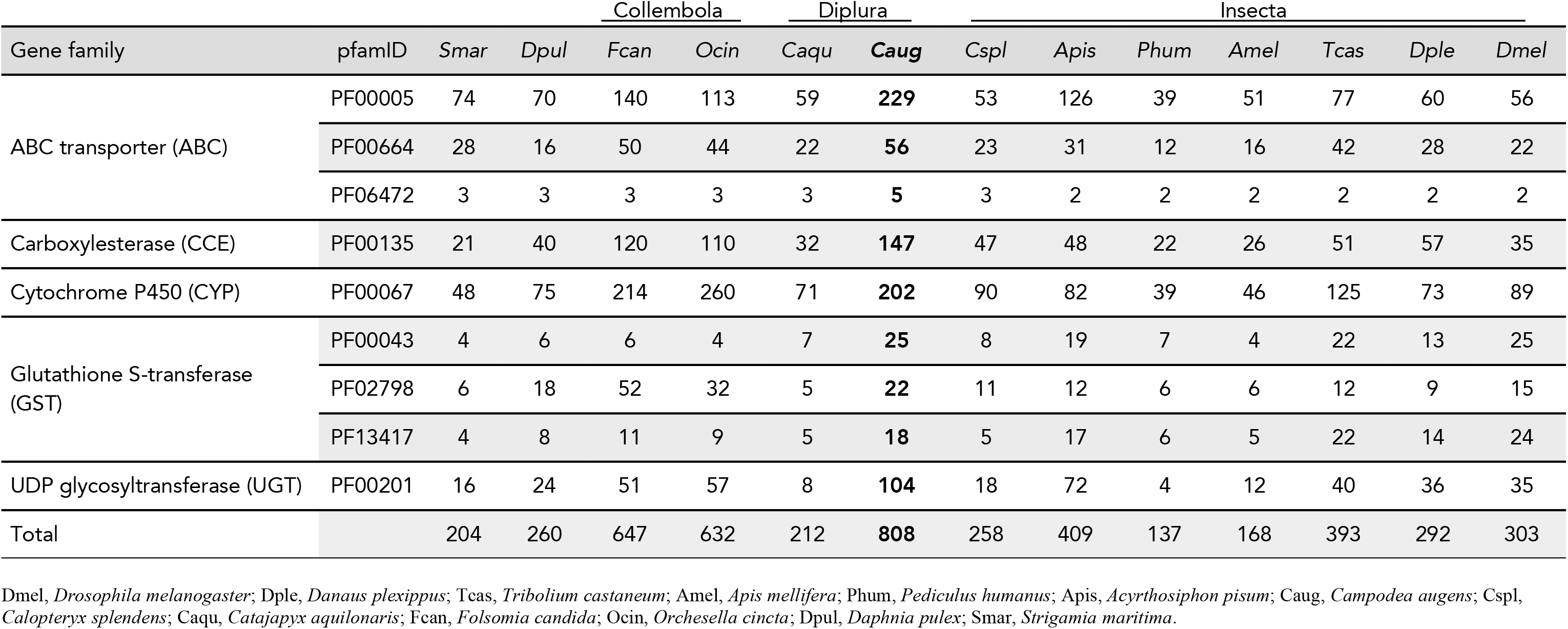
Counts of protein Pfam domains associated with detoxification enzymes in *C. augens* and 12 other arthropods.

In contrast to other hexapods, diplurans have the remarkable capacity to regenerate lost body appendages (antennae, cerci and legs) over a series of moults, even as adults (Conde 1955; Lawrence 1953; Maruzzo et al. 2005; B.K.Tyagi & Veer 2016; Whalen & Sampedro 2010). The genetic amenities to facilitate these adaptations have not been investigated to date. However, candidate genes related to regenerative capacities of tissues have been studied in model organisms (Bergmann & Steller 2010). For example, caspases are protease enzymes involved in programmed cell death that also play a role in tissue regeneration and differentiation in some animal models (Bergmann & Steller 2010; Sun & Irvine 2014), including *D. melanogaster*. Apoptosis and caspase activity have been detected in regenerating tissue (Boland et al. 2013; Shalini et al. 2015). In *C. augens*, we identified expansions of the caspase gene family, C-type lectin-like proteins, and inhibitor of apoptosis proteins (IAPs) that can bind to and inhibit caspases and other proteins involved in apoptosis. In particular, we found a high abundance of caspases in *C. augens* (35 proteins), while in the 12 other arthropods under consideration, the counts range from four in *D. plexippus* to 18 in *D. pulex* (Supplementary Table 11). Whether or not the expansion of gene families linked to apoptosis and regeneration is indeed related to the high regenerative potential of *C. augens* needs further experimental evaluations. Transcriptomic and proteomic studies in tissues under regeneration might help to explain these observed genomic patterns.

### Endogenous viral elements in the genome of *C. augens*

We found seven fragments of non-retroviral integrated RNA viruses (NIRVs) in the genome of *C. augens* that resemble viral sequences of the genus Quaranjavirus, with the highest similarity to Wuhan Louse Fly Virus 3, Wuhan Mosquito Virus 3 and 5, Shuangao Insect Virus 4, and Wellfleet Bay Virus (Supplementary Table 12). Quaranjaviruses belong to Orthomyxoviridae, a family of segmented negative single-stranded RNA viruses (-ssRNA) that also includes the influenza viruses (Presti et al. 2009). All quaranjavirus-like insertions found in *C. augens* encode the PB1 protein (polymerase basic protein 1) (Supplementary Table 12), a subunit of the RNA-dependent RNA polymerase (RdRp), which in Orthomyxoviridae is typically a heterotrimeric complex (formed by the PA, PB1, and PB2 proteins) (Stubbs & te Velthuis 2014), contrary to RdRps from other RNA viruses (e.g. double-stranded and positive-stranded RNA viruses) which are typically monomeric (Wolf et al. 2018). The corresponding open reading frames of PB1 proteins are incomplete, though, and contain stop codons and/or frameshift mutations (Supplementary Fasta), suggesting that these elements are not functional. As indicated by the phylogenetic analyses in Figure 5 and Supplementary Figure 13 and 14, the putative viral insertions are more closely related to a cluster of quaranjaviruses that have been identified in arthropod samples through metagenomic RNA sequencing (Li et al. 2015) than to other quaranjaviruses (e.g., Johnston Atoll quaranjavirus and Quaranfil quaranjavirus) (Supplementary Table 12). Only one case of a quaranjavirus-like insertion has been previously described in the genome of the black-legged tick (Katzourakis & Gifford 2010). However, this sequence is more closely related to that of Johnston Atoll quaranjavirus and encodes a different viral protein (Supplementary Figure 14). The presence of orthomyxovirus-like NIRVs in *C. augens* is of particular interest since Orthomyxoviridae include the Influenza viruses. Recent studies have suggested a central role of arthropods in the origin and evolution of viral species (Li et al. 2015; Shi et al. 2016; Dudas & Obbard 2015), and NIRVs are instrumental for studying virus-host interactions (Olson & Bonizzoni 2017; Aswad & Katzourakis 2016) and to gain insights on viral long-term evolution.

**Fig. 5.**
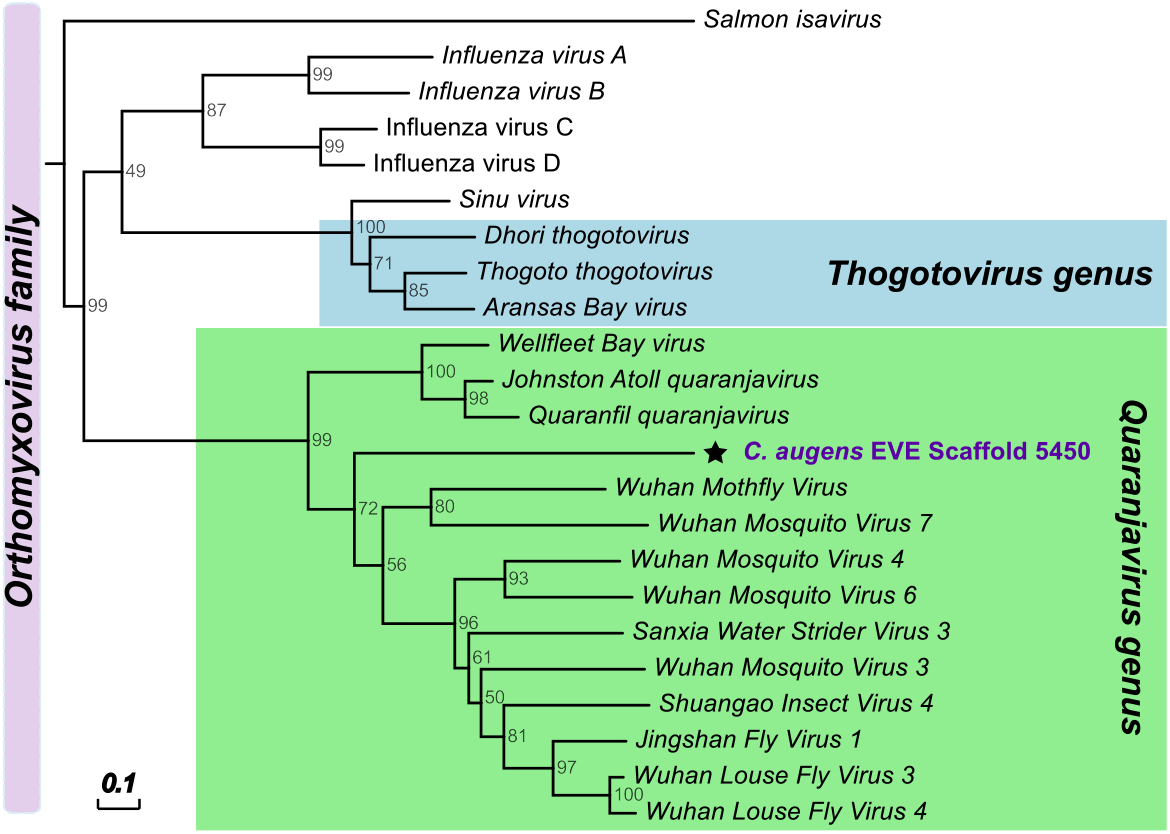
Phylogenetic relationships of the putative endogenous viral elements in *C. augens* genome related to - ssRNA viruses of the Orthomyxoviridae family. All endogenous viral elements found in *C. augens* correspond to orthomyxoviral Polymerase Basic protein 1 (PB1) (Pfam Id PF00602; “Flu_PB1”) (See also Supplementary Fig. 12 and 13). Neighbour-joining trees was constructed using amino acid sequences of EVEs and BP1 proteins of representatives of the Orthomyxoviridae family. Support for trees was evaluated using 1,000 pseudo replicates. Node values correspond to the bootstrap support. Scale bar indicate amino acid substitutions per site. Green box highlights viruses of the Quaranjavirus genus, while light blue box indicates the Thogotovirus genus.

## Conclusions

We sequenced and annotated the 1.2-Gbp genome of *C. augens*, a blind and ancestrally wingless hexapod belonging to the systematic order Diplura. Gene structure analysis highlighted a genome-wide trend of remarkably long introns that span 27% of the assembly. We identified a paucity of photoreceptor genes mirroring at the genomic level the secondary loss of an ancestral external photoreceptor organ, and by contrast the presence of the largest ionotropic receptor gene repertoire in the animal kingdom, which account for ca. 8% of the total *C. augens* gene set. We further detected expansions of gene families related to detoxification and carbohydrate metabolism, which might reflect adaptations in *C. augens*’ foraging behaviour. To date, most of the existing and recent experimental data on diplurans derive from *C. augens* and *C. aquilonaris*, for example, studies on brain anatomy and circulatory organs (Gereben-Krenn & Pass 1999; Böhm et al. 2012). We thus think *C. augens* and *C. aquilonaris* represent good candidates as model species for diplurans. The availability of their genomes can enable detailed molecular studies and unlock the potential of linking diplurans’ peculiar morphological/behavioural features to genetic evidence. The *C. augens* genome opens up novel opportunities to study both the under-explored biology of diplurans as well as the origin and evolution of gene families in insects.

## Methods

### Sample collection and sequencing

*C. augens* samples were collected at Rekawinkel, Austria (48° 11’ 06,68’’ N, 16° 01’ 28,98’’ E) and determined based on the key of Palissa (1964) complemented by more recent taxonomic information. *C. augens* genome size was estimated to be ca. 1.2 Gbp by flow-through cytometry following the protocol given by DeSalle et al. (2005) using *Acheta domestica* as size standard (ca. 3.9 Gbp). Two female adults were used for genome sequencing. Before DNA extraction, the individuals were carefully washed to remove any non-target organisms that might adhere on the body surface. Genomic DNA was extracted using a Qiagen DNeasy Blood & Tissue kit (Qiagen, Hilden, Germany) and following the “insect” nucleic acid isolation protocol described by the manufacturer. Four Illumina paired-end (PE) sequencing libraries, 2 × 350-bp and 2 × 550-bp insert sizes, were constructed using Illumina’s TruSeq DNA Nano kit (Illumina, San Diego, CA, USA) following the standard protocol. Four additional mate pair (MP) libraries (3-, 6-, 9- and 12-Kbp insert sizes) were prepared using Illumina’s Nextera Mate Pair kit with size selection performed on precast E-gel (Life Technologies, Europe BV) 0.8% agarose gels. Libraries were sequenced on a HiSeq 2500 platform (Illumina, San Diego, CA, USA) using a read-length configuration of 2 × 100 bp. All raw reads (around 2.1 billion in total) are deposited in the NCBI Sequence Read Archive (SRA) under the accession numbers SRX3424039–SRX3424046 (BioProject: PRJNA416902).

### Genome assembly

Low-quality reads and reads with adaptor and spacer sequences were trimmed with trimmomatic v0.36 (Bolger et al. 2014). The kmer content of reads from short-insert libraries was calculated using KmerGenie v1.7023 (Chikhi & Medvedev 2014). GenomeScope v1.0 (Vurture et al. 2017) was used to assess the heterozygosity level. All libraries were initially screened to detect reads derived from 16S genes of bacterial and archaeal species with the program parallel-meta v3 (Jing et al. 2017) using the “shotgun” option. Further screening of the reads were performed with Kraken (Wood & Salzberg 2014), using a set of custom databases representing full genomes of archaea, bacteria, fungi, nematodes, plants, protozoa, viruses, and worms. An initial draft assembly constructed from short-insert libraries using sparseassembler (Ye et al. 2012) was used for assessing the presence of contamination by Taxon-Annotated GC-Coverage (TAGC) plots as in Blobtools (Laetsch & Blaxter 2017). For taxonomic annotation of the draft contigs, results from MegaBLAST v2.3.0+ (Camacho et al. 2009) using the NCBI nucleotide database (nt) as well as Diamond v0.8.34.96 (Buchfink et al. 2015) with UniRef90 protein database (Suzek et al. 2015) were used as input for Blobtools (run using the “bestsumorder” rule). Read coverage for each contig was calculated by mapping the four libraries to the draft assembly using BWA v0.7.13 (Li & Durbin 2009). We further analysed the assembly with PhylOligo (Mallet et al. 2017) using hierarchical DBSCAN clustering (phyloselect.py), and with Anvi’o (Eren et al. 2015) following the procedure shown in Delmont & Eren (2016) in order to visualise the assembly (only contigs > 2.5 Kbp) along with GC content and mapping profiles to detect potential contaminants. In order to identify *C. augens* mitochondrial sequences, the draft contigs were compared with blastn (Camacho et al. 2009) to available mitochondrial genomes of two dipluran species (*Campodea fragilis* and *Campodea lubbocki*) (Podsiadlowski et al. 2006). The results of the above-mentioned analyses were used to filter the raw reads: contigs that showed a substantially different coverage/ GC composition relative to that of the main cluster of contigs or hits to putative contaminants were used to filter the reads by mapping with BWA. Filtered short-insert libraries were then reassembled using the heterozygous-aware assembler Platanus v1.2.4 (Kajitani et al. 2014) with its default parameters. Redundans v0.13a (Pryszcz & Gabaldón 2016) was used to decrease the assembly redundancy, which occurs when heterozygous regions are assembled separately (Kelley & Salzberg 2010). Scaffolding of the contigs was performed with SSPACE v3.0 (Boetzer et al. 2011) using both short-insert and mate pair libraries. A final step of scaffolding was performed with AGOUTI v0.3.3 (Zhang et al. 2016) using *C. augens* transcripts obtained from reassembling raw reads (SRR921576) from whole-body RNA-seq data produced in the context of the 1KITE project (www.1kite.org). Scaffolds < 1 Kbp were filtered out unless they showed a significant (e-value <1e-05) similarity to entries in the NCBI nr database. To assess the quality of the assemblies and to monitor the improvement in assembly quality, each step was monitored with BUSCO v3.0.2 (Waterhouse et al. 2018), using the arthropoda_odb9 dataset. The completeness and accuracy of the genome assembly were further assessed by mapping the paired-end reads to the assembly using BWA and the assembled transcripts to the genome assembly using TopHat (Trapnell et al. 2009). This Whole Genome Shotgun project has been deposited at DDBJ/ENA/GenBank under the accession VUNT00000000. The version described in this paper is version VUNT01000000. *C. augens* mitochondrial genome was assembled separately using NOVOPlasty v2.6.7 (Dierckxsens et al. 2017), and annotated using MITOS (Bernt et al. 2013). The mitogenome sequence was deposited in GenBank under the accession number MN481418. The assembly and additional information such as the short (< 1 Kbp) scaffolds can also be obtained from http://cegg.unige.ch/campodea_augens.

### Genome annotation

A custom *C. augens* repeat library was constructed using RepeatModeler and RepeatClassifier (http://www.repeatmasker.org/). The species-specific repeats library and the RepBase (Bao et al. 2015) library (update 2 March 2017) were used to mask the genome with RepeatMasker v4.0.7 (http://www.repeatmasker.org). Automated annotation was performed with MAKER pipeline v2.31.8 (Holt & Yandell 2011) on the masked genomic scaffolds. Evidence-based gene structural annotations were inferred using eight arthropod proteomes obtained from OrthoDB v9 (Zdobnov et al. 2017), all entries from the SwissProt database (Boutet et al. 2016), and the *C. augens* transcripts obtained by re-assembling raw reads (SRR921576) from whole-body RNA-seq data. MAKER was set to infer gene models from all evidence combined (not only transcripts), and gene predictions without transcript evidence were allowed. AUGUSTUS v3.2 (Stanke & Morgenstern 2005), trained with parameters obtained from *C. augens* single-copy genes identified with BUSCO (Seppey et al. 2019), was used for *ab initio* gene prediction. The *C. augens* genome along with gene predictions, raw reads, and additional tracks were loaded into Apollo (Lee et al. 2013), which was used to visualise and manually curate gene models of families of interest. Chimeric gene models were split into individual genes, erroneously split gene models were joined, and exon–intron boundaries were corrected according to transcript evidence. This resulted in 23,992 gene models, which were assigned identifiers from CAUGE_000001 to CAUGE_023992 (where ‘CAUGE’ stands for *Campodea AUGEns)*. Functional annotation of predicted protein-coding genes was performed using InterProScan for finding conserved domains. Additionally, proteins were searched against Uniref50 database (Suzek et al. 2015) and clustered in OrthoDB v10 (Kriventseva et al. 2019) for finding conserved functions. To assign *C. augens* genes to Orthologous Groups (OGs), the predicted gene set was clustered with the gene sets of 169 arthropods in OrthoDB v10 database (Kriventseva et al. 2019). Subsequently, a phylogenomic analysis was conducted based on 371 single-copy genes shared among *C. augens* and 13 other arthropods including a) insect species: *Acyrthosiphon pisum* (pea aphid), *Apis mellifera* (honey bee), *Calopteryx splendens* (banded demoiselle), *Danaus plexippus* (monarch butterfly), *Drosophila melanogaster* (fruit fly), *Pediculus humanus* (body louse) and *Tribolium castaneum* (red flour beetle); b) representatives of ancestrally wingless hexapods: *Acerentomon* sp. (coneheads), *Catajapyx aquilonaris* (northern forcepstail), *Folsomia candida* (springtail), *Orchesella cincta* (springtail); and c) two non-hexapod species: *Daphnia pulex* (water flea) and *Strigamia maritima* (centipede). No genome is available for the Protura order, but since this taxon is crucial in the context of our phylogenetic analysis, we re-assembled the transcriptome of *Acerentomon* sp. from raw reads deposited at SRA (SRR921562) and used the resulting assembly as the source of phylogenomic markers. The *Acerentomon* sp. transcriptome was assembled using Trinity (Grabherr et al. 2011) with its default parameters. For extracting single-copy genes and performing phylogenomic analysis, we followed the pipeline described in Waterhouse et al. (2018). Briefly, MAFFT (Katoh & Standley 2013) was used for generating multiple sequence alignments of the amino acid sequences, alignments were then automatically trimmed with Trimal v1.2.59 (Capella-Gutiérrez et al. 2009), and RAxML v8 (Stamatakis 2014) was used for inferring a phylogenetic tree under the maximum likelihood method (Supplementary Note). The resulting phylogenetic tree was viewed and annotated using the R package ggtree (Yu et al. 2017).

### Gene family evolution (expansions and contractions)

To study gene family evolution, we first analysed expansions and contractions of gene families using the software CAFE (Han et al. 2013) and OrthoDB clusters (Kriventseva et al. 2019). Furthermore, to detect major size changes in gene families between the dipluran *C. augens* and more recent lineages of hexapods, we also compared the gene counts of gene families in *C. augens* to the average and median of gene counts from the six species of Insecta included in the analysis. The clustering data at the level Arthropoda from OrthoDB were converted to CAFE input files, and CAFE v4.1 was used to calculate the average gene expansion/contraction rate and to identify gene families that have undergone significant size changes (*P* < 0.01) across the phylogeny (number of gene gains and losses). The rooted phylogenetic tree required for the analysis was inferred with RAxML as described in the previous section, excluding data for the proturan *Acerentomon* sp., for which a genome is not yet available. The time-calibrated phylogeny was estimated using the r8s program (Sanderson 2003) using calibration points retrieved from timetree.org (Kumar et al. 2017) and from Misof et al. (2014) (Misof et al. 2014). CAFE was run using the default *P*-value thresholds, and estimated rates of birth (λ) and death (μ) were calculated under a common lambda across the tree and additionally under different lambda for Collembola, Diplura, Insecta, and non-hexapod species. Using OrthoDB clusters, we additionally identified gene families in *C. augens* significantly larger or smaller with respect to the average of counts in Insecta (s. str.) using a two-tailed Fisher’s exact test in R. The *P*-values were sequentially corrected using the Benjamini and Hochberg method (Benjamini & Hochberg 1995) in R, and only gene families with corrected *P*-values less than 0.01 were considered as significantly different.

We complemented the analysis of gene families by performing an overall protein domain analysis. We analysed the proteomes of all 13 arthropod species included in the phylogenomic analysis using InterProScan (Jones et al. 2014) and searched for known Pfam protein domains, filtering out entries covering less than 75% of the corresponding Pfam hidden Markov model (HMM). Applying the same Fisher’s test as above, we identified protein domains with significantly larger counts in *C. augens* respect to the average found in the insect species.

### Identification and phylogenetic analysis of endogenous viral elements

*C. augens* sequences with high identity with those of viral peptides were extracted and used to screen the GenBank non-redundant (nr) database in a reciprocal tBLASTn search. Analyses of the putative insertion sites (i.e., raw read coverage of the sites and corresponding flanking regions, presence of host genes in the surrounding regions, and phylogenetic analyses) were performed to exclude the possibility of dealing with assembly artefacts. For the phylogenies, regions of putative endogenous viral elements were translated into amino acid sequences and individually aligned to amino acid sequences of the polymerase basic protein 1 (PB1) from orthomyxoviruses retrieved from GenBank database (Supplementary Table 13). Alignments were generated with MAFFT (Katoh & Standley 2013), and the phylogenetic trees were constructed using the neighbour-joining method with a non-parametric bootstrap analysis (1,000 replicates). The resulting trees were visualised and annotated in Evolview (He et al. 2016).

### In-depth analysis of genes of interest

The chemoreceptor genes in the GR and IR families were manually annotated. An attempt was made to build all models that include at least half of the coding region of related full-length IRs, including genes truncated due to problems with the genome assembly (sequence gaps or misassemblies indicated by single Ns) and pseudogenes. *C. augens* IRs were named in a series starting from Ir101 (Ir101-117 share a single intron) to avoid confusion with *D. melanogaster* genes, which were named after their cytological location and hence stopping with Ir100a. TBLASTN searches with representatives from insects, such as the damselfly *Calopteryx splendens* (Ioannidis et al. 2017), were used to identify genes using no filter for low complexity regions, word size of two, and an E-value of 1,000. Models for the GRs and the intron-containing IRs were built in the Apollo genome browser (Lee et al. 2013) employing the automated models from MAKER and RNA-seq for support. The large set of intronless IR models and their many pseudogenes were built in a text editor, because pseudogene models are difficult to build in the Apollo browser. Encoded proteins were aligned with representatives from other species using ClustalX v2.0 (Larkin et al. 2007), and models refined in light of these alignments. The final alignments were trimmed using TrimAl v4.1 (Capella-Gutiérrez et al. 2009) with the “gappyout” option for the GRs and the “strictplus” option for the IRs, which have more length variation than the GRs. This analysis excludes the few available partial GR sequences from two orders of early diverging insects (Archaeognatha and Zygentoma) (Missbach et al. 2014). Unfortunately, the available draft genomes of these two orders are too fragmented to reliably annotate their GRs (Brand et al. 2018), and the GRs have been only partially annotated for one of the three draft genomes of Collembola (Wu et al. 2017). Given the phylogenetic distance of Diplura to Collembola and ancestrally wingless insects (*i.e.*, Archaeognatha and Zygentoma), inclusion of their GRs is unlikely to change our assessment of the dipluran GR complement. Maximum likelihood phylogenetic analysis was performed using PhyML v3.0 (Guindon et al. 2010) at the PhyML webserver using the software’s default settings (http://www.atgc-montpellier.fr/phyml/). Trees were prepared using FigTree v1.4.2 (http://tree.bio.ed.ac.uk/software/figtree/). The Hox genes, photoreceptors and clock genes were searched with TBLASTN on the genome assembly using a set of known proteins and by InterProScan results. Gene models were checked and curated in the Apollo browser. For candidate opsins, the *C. augens* gene set and whole genome were scanned using 16 reference opsins (Hering & Mayer 2014) with blastp and tblastn respectively, matches with an e-value < 1e-4 were retained as candidate opsins. Additionally HMM profiles created for the nine major opsin clades (cnidopsins, vertebrate c-opsins, pteropsins, Group 4 opsins, arthropsins, melanopsins, non-arthropod r-opsins, arthropod visual opsins and onychopsins) (Hering et al. 2012) were used to scan the *C. augens* gene set.

## Supporting information

curated_chemosensory_genes

## Data availability

This Whole Genome Shotgun project has been deposited at DDBJ/ENA/GenBank under the accession VUNT00000000. The version described in this paper is version VUNT01000000. Raw sequencing data has been deposited in the NCBI Sequence Read Archive (SRA) under the accessions SRX3424039-SRX3424046. The annotated mitochondrial genome sequence was deposited in GenBank under the accession number MN481418. The genome and mitochondrial assemblies, gene set and additional data such as the short scaffolds (< 1 Kbp), the curated chemoreceptor genes, and the supermatrix used for the species phylogeny in Figure 2 can also be obtained from http://cegg.unige.ch/campodea_augens. Credentials for accessing the Apollo browser are provided upon request.

## Acknowledgments

This work was supported by Swiss National Science Foundation grant [31003A_143936] to EMZ. RMW acknowledges support from Swiss National Science Foundation grant [PP00P3_1706642].

We thank Stephen Richards (Baylor College of Medicine) for granting us permission to use the unpublished genome of *Catajapyx aquilonaris*.

## Author contributions

EZM and MM managed and coordinated the study; MM performed genome assembly, annotation, orthology, and subsequent analyses; FAS carried out sequencing of samples; BM and ON provided genome size estimates based on flow-through cytometry and organized the sample collection; SN collected wild *C. augens* samples; HMR performed the identification and characterization of chemoreceptor genes; MAG assembled the mitogenome and analysed gene models for *Hox* genes; MM wrote the manuscript and the supplementary information with contributions from RMW, ON, BM, HMR and EZM; HMR wrote the results for chemoreceptor genes; MM prepared figures and tables for the manuscript and supplementary information; All authors read, corrected and commented on the manuscript.

## Competing interests

The authors declare no competing financial interests.

