## supplementary_file for "The genome of the blind soil-dwelling and ancestrally wingless dipluran *Campodea augens*, a key reference hexapod for studying the emergence of insect innovations"

### SUPPLEMENTARY NOTES

##### *Contamination*

By scanning raw reads for bacterial contamination, we detected a small number of reads with high nucleotide similarity to fragments of the *Wolbachia* genome. *Wolbachia* endosymbionts commonly infect arthropods and can occasionally transfer parts of their genome to the nuclear genome of their arthropod host (Robinson et al. 2013). The low abundance of reads possibly originating from a *Wolbachia* genome suggests that the samples used for sequencing the genome of *C. augens* were likely not infected by the endosymbiont. We consider it more likely that the *Wolbachia*-like reads originated by historical Lateral Gene Transfer (LGT) events. In total, we found on 24 scaffolds in the *C. augens* assembly 29 short nucleotide sequence sections of putative bacterial origin and likely incorporated into the genomes by LGT (Supplementary Table 6).

##### *Mitochondrial genome*

The mitochondrial genome (mitogenome) of *C. augens* was assembled separately from the nuclear genome using the de novo assembler NOVOPlasty v2.6.7, resulting in an assembly of 15,735 bp in length, a size comparable to that of the mitogenome of *Campodea fragilis* (14,965 bp) (Podsiadlowski et al. 2006) (Supplementary Fig. 7).

The annotation of the *C. augens* mitogenome was performed using the web version of the metazoan mt genome annotator MITOS (Bernt et al. 2013), using the invertebrate genetic code and the RefSeq 63 Metazoa as reference database. The *C. augens* mitogenome encodes for 13 protein-coding genes, 22 tRNAs, and two rRNAs. The order of genes is identical to *Campodea fragilis* (Supplementary Table 7, Supplementary Fig. 7). The AT-rich control region (around 1 Kbp in size) is located between the *trnL* and *rrnS* genes.

***Phylogenomic analysis***

A phylogenomic analysis was conducted using *C. augens* and 13 other arthropod species of interest including a) insect species: *Drosophila melanogaster* (fruit fly), *Danaus plexippus* (monarch butterfly), *Tribolium castaneum* (red flour beetle), *Apis mellifera* (honey bee), *Pediculus humanus* (body louse), *Acyrthosiphon pisum* (pea aphid), *Calopteryx splendens* (banded demoiselle); b) representatives of Collembola, Diplura and Protura: *Folsomia candida* (springtail), *Orchesella cincta* (springtail), *Catajapyx aquilonaris* (northern forcepstail), *Acerentomon sp.* (coneheads); and c) two non-hexapod species: *Daphnia pulex* (water flea) and *Strigamia marítima* (centipede).

The order Prutura (coneheads) still lack a sequenced genome, but a representative of this taxon is crucial in the context of our phylogenetic analysis to resolve the relationships among ancestrally wingless hexapod. Therefore we re-assembled the transcriptome of *Acerentomon sp.* from raw reads deposited at SRA (SRR921562), produced in the context of 1KITE project, and used it as the source of phylogenomic markers. Assembly of the transcriptome was performed using Trinity (Grabherr et al. 2011) with default parameters. For extracting single-copy genes shared among the 13 species of interest mentioned above we used BUSCO following the pipeline described in (Waterhouse et al. 2018), which allows to extract single-copy genes using as input gene sets, genomes or transcriptomes (scripts available at https://gitlab.com/ezlab/busco_usecases/tree/master/phylogenomics). For building the phylogenetic tree, we employed 358 single-copy genes, out of the ca. 1000 universal single-copy genes expected to be found in arthropoda according to BUSCO arthropoda dataset (Waterhouse et al. 2018; Kriventseva et al. 2019), that were present in all the 14 species under consideration, thus minimizing missing data in the matrix. The corresponding OrthoDB IDs of these 358 genes can be found in the text file of partitions "Species_phylogeny_partitions.txt" at http://cegg.unige.ch/campodea_augens. The corresponding description can be retrieve from ODB9 website (https://www.orthodb.org/v9/index.html) by using the ID as query. The amino acid sequences of the 358 single-copy genes shared among the 13 species were then aligned individually, with MAFFT (Katoh & Standley 2013) using the L- INS-i algorithm. Each protein multiple sequence alignment (MSA) was visually inspected to detect presence of misalignments. Each MSA were subsequently trimmed with Trimal v1.2.59 for automatic trimming using the "-auto" option. MSAs were then concatenated in a supermatrix 145,145 aa long. The phylogenetic analysis was performed using a partitioned model-based tree reconstruction, considering the selected single-copy genes as evolutionary units. We conducted a meta-partition analysis using the protTest software version 3.4.2 (Darriba et al. 2011). protTest was run on each individual trimmed MSA, and the best model according to AICc was assigned to each MSA. The MSA and partition file are available at http://cegg.unige.ch/campodea_augens. To infer the phylogenetic tree based on a Maximum Likelihood (ML) approach we used the PTHREADS version of RAxML v8 (Stamatakis 2014). The partition information derived with protest was provided to RAxML using the "-q" option. We estimated 100 bootstrap replicates and drawn bootstrap values on the best ML tree using the standard RAxML options. Tree was rooted by selecting S*trigamia maritima* as the outgroup species, and it was annotated using the R package ggtree (Yu et al. 2017). Alternatively, we also analyzed the whole matrix using various mixture models implemented in IQ-TREE v1.6.10 (Nguyen et al. 2015). Support values were estimated by ultrafast bootstrapping for 1000 replicates. The results were congruent with the topology obtained using partitions in RAxML, placing Diplura as a sister clade to Insecta. The trees obtained with IQ-TREE using different mixture models are reported below:

EX_EHO model:
(((ACERE:0.3722466053,(FCAND:0.1932572727,OCINC:0.2238088850)100:0.2884263193)98:0.0367511722,((((AMELL:0.2573580956,
((DMELA:0.3680389924,DPLEX:0.3146272518)100:0.0517523676,TCAST:0.2464391402)100:0.0390115020)100:0.0300424702,
(APISU:0.4196732606,PHUMA:0.2834345904)100:0.0378272404)100:0.0313712042,CSPLE:0.2512345655)100:0.0629332830,
(CAQUI:0.2327130107,CAUGE:0.3176562358)100:0.0728476348)86:0.0196511931)100:0.0409085274,DPULE:0.3468198516,SMARI:0.3432093
826);

EX2 model:
(((ACERE:0.3653026123,(FCAND:0.1938355157,OCINC:0.2221617439)100:0.2801976402)99:0.0397104878,((((AMELL:0.2547959289,
((DMELA:0.3606424543,DPLEX:0.3093933852)100:0.0541921806,TCAST:0.2435941194)100:0.0410729731)100:0.0320518128,
(APISU:0.4105711164,PHUMA:0.2791013121)100:0.0405677499)100:0.0335293535,CSPLE:0.2501829340)100:0.0644800555,
(CAQUI:0.2315965170,CAUGE:0.3130919435)100:0.0734698875)91:0.0221596281)100:0.0427428037,DPULE:0.3407857788,SMARI:0.3374211
934);

EX3 model:
(ACERE:0.3688243196,(((((AMELL:0.2545129145,
((DMELA:0.3648701223,DPLEX:0.3122653738)100:0.0528636633,TCAST:0.2452033362)100:0.0398988541)100:0.0308996506,
(APISU:0.4142856836,PHUMA:0.2812713489)100:0.0386355939)100:0.0321504263,CSPLE:0.2520255112)100:0.0638158811,
(CAQUI:0.2335132755,CAUGE:0.3146898327)100:0.0733888059)85:0.0205965418,
(DPULE:0.3445738398,SMARI:0.3396821478)100:0.0415661780)100:0.0377115529,
(FCAND:0.1925376333,OCINC:0.2228080143)100:0.2858256698);

EHO model:
(((ACERE:0.3425637704,(FCAND:0.1868623586,OCINC:0.2116748237)100:0.2562105105)100:0.0418614319,((((AMELL:0.2436079148,
((DMELA:0.3374428965,DPLEX:0.2904640996)100:0.0543869361,TCAST:0.2311603875)100:0.0404896155)100:0.0322090250,
(APISU:0.3831956100,PHUMA:0.2625109490)100:0.0425219333)100:0.0330212671,CSPLE:0.2371375195)100:0.0604887373,
(CAQUI:0.2194560961,CAUGE:0.2942413438)100:0.0709061967)87:0.0232039777)100:0.0440228678,DPULE:0.3187625410,SMARI:0.3170877
377);

##### *Chemoreceptors*

The GR and IR families were manually annotated (see supplementary file Curated_chemosensory_genes.txt). This effort was complicated, especially in the case of the IRs, by three factors. First, the genome assembly has many single-N gaps, which mark the locations of artifactual duplications and more rarely deletions. As these sometimes affect parts of genes, and occasionally entire genes, care was taken not to include such identical adjacent direct repeat fragments of genes. If such a duplication had occurred within a gene, the genomic sequence was repaired to fix the model, however if a gene terminus was missing due to a single N the assembly was not repaired and the model treated as incomplete. Second, pseudogenes were included if they encoded at least 50% of the length of a related intact IR (these IRs vary in length by two-fold from 445 to 897 amino acids, so no single length criterion could be employed). In most cases pseudogenes result from one or more stop codons or short frameshifting indels, and occasionally loss of either terminus or large internal deletions, however there are some with so many pseudogenizing mutations that they were too difficult to reconstruct and include in the analysis, despite representing more than half length of a gene (generally those with more than seven pseudogenizing mutations). Third, there are innumerable gene fragments encoding less than half of a related intact IR, and most of these are assumed to be pseudogenic fragments and ignored. Each set of closely related IRs was iteratively searched against the genome using TBLASTN searches to find all close relatives in an expanded lineage, and divergent relatives identified in the same scaffolds were sequestered for later searching as a way to discover the entire family (commonly around 20% amino acid identity for only the central relatively-conserved region of the protein). These divergent relatives were then used as queries to explore divergent lineages towards the end of the effort. Extensive searches with each of these divergent lineages eventually found only already annotated genes, suggesting that all IR genes in this genome have been recovered.

As described in the main text, the GR family is modestly-sized with 41 genes, seven of them pseudogenes. The family composition is relatively straightforward with 13 relatives of the sugar receptor subfamily in insects, no relatives of the carbon dioxide and fructose receptors commonly conserved in insects, and 28 otherwise divergent members.

In contrast, the IR family is enormously expanded with 2431 genes and pseudogenes named and included in the analysis. Conserved orthologs were found for the co-receptors Ir8a, 25a, and 76b (named for their *D. melanogaster* orthologs), however Ir25a has been duplicated at least four times, with additional gene fragments suggesting there is a fifth copy. These 4 or 5 Ir25a orthologs might be involved in dimerization with the many divergent IRs found.

The numbers of obvious pseudogenizing mutations (stop codons, frameshifts, and large deletions) were noted for each pseudogene. The histogram in Figure S11 reveals that the majority have single mutations. The caveat noted above applies to the right hand of this histogram, as some reasonably long pseudogenes with too many mutations were too hard to reconstruct and hence excluded from the set, but the preponderance of pseudogenes with single mutations versus those with 2-5 mutations is real as the latter were consistently reconstructed.

##### *Expansion of gene families related to sugar metabolism and transport*

In relation to the foraging ecology of *C. augens*, we also found a number of expanded gene families associated with carbohydrate metabolism and transport. This may reflect adaptations to the carbohydrate-rich diet of *C. augens*. In particular, we identified substantial changes in the gene copy numbers of trehalose receptors, glucose-methanol-choline (GMC) oxidoreductases, glycosyltransferases, histidine phosphatases, and glycosyl hydrolases (Supplementary Tables 8 and 9). Catalytic functions of histidine phosphatases include phytase activity to hydrolyse phytic acid, present in many plants tissues. Glycosyl hydrolases catalyze the hydrolysis of glycosidic bonds in complex sugars and they are involved in the degradation of cellulose, hemicellulose, and starch. Genes involved in sugar transport, such as the major facilitator superfamily (MFS), sodium sulfate symporters, and the EamA-like transporter family, were also found to be expanded (Supplementary Tables 8 and 9).

### SUPPLEMENTARY TABLES

**Table S1:** Summary of *C. augens* sequencing libraries.

| Library | Library type | Insert size | Read length (bp) | Number of reads (million) | Sequencing output (Gb) | Genome coverage (X) |
|---|---|---|---|---|---|---|
| 350_L1 | paired-end | 350 bp | 100 | 277,91 | 27,79 | 23,16 |
| 350_L6 | paired-end | 350 bp | 100 | 358,09 | 35,81 | 29,84 |
| 550_L2 | paired-end | 550 bp | 100 | 270,31 | 27,03 | 22,53 |
| 550_L7 | paired-end | 550 bp | 100 | 298,62 | 29,86 | 24,89 |
| m3 | mate pair | 3 kb | 100 | 277,62 | 27,76 | 23,13 |
| m6 | mate pair | 6 kb | 100 | 230,00 | 23,00 | 19,17 |
| m9 | mate pair | 9 kb | 100 | 218,07 | 21,81 | 18,17 |
| m12 | mate pair | 12 kb | 100 | 207,42 | 20,74 | 17,29 |
| | | | Total | 2138,03 | 213,80 | 178,17 |

**Table S2:** Statistics of *C. augens* genome compared to the four available ancestrally wingless hexapod genomes, and other insects with large genomes.

| | Non-insect hexapods | | | | | Insects | | | | |
|---|---|---|---|---|---|---|---|---|---|---|
| Metric | *Campodea augens* | *Catajapyx aquilonaris* | *Folsomia candida* | *Orchesella cincta* | *Holacanthella duospinosa* | *Calopteryx splendens* | *Blatella germanica* | *Locusta migratoria* | *Medauroidea extradendata* | *Clitarchus hookeri* |
| Assembly size (Mb) | 1130 | 312 | 221 | 286 | 328 | 1630 | 2040 | 5760 | 2600 | 3800 |
| No. of scaffolds | 18,765 | 132,454 | 162 | 9,402 | 62,430 | 8,896 | 24,818 | 1,397,492 | 135,691 | 785,781 |
| No. of contigs | 70,883 | 151,491 | 228 | 9,641 | 70,136 | 645,677 | 317,827 | 1,397,492 | 167,455 | 906,073 |
| Scaffold L/N50 (Kb) | 1,367/235 | 2,531/31 | 8/6520 | 925/66 | 242/310 | 1,013/422 | 576/1050 | 174,483/10 | 17,103/43 | 2,981/316 |
| Contig L/N50 (Kb) | 8,725/33 | 6,775/12 | 12/4890 | 936/64 | 985/80 | 83,275/4 | 39,325/12 | 174,483/10 | 26,796/28 | 21,033/46 |
| Max scaffold length (Mb) | 2.19 Mb | 0.42 | 28.53 | 0.81 | 2.81 | 2.78 | 7.47 | 0.11 | 0.43 | 4.95 |
| Max contig length (Kb) | 390 | 142 | 20230 | 807 | 1270 | 63 | 1930 | 106 | 250 | 626 |
| No. of scaffolds > 50 Kb | 5,264 | 1,290 | 83 | 1,288 | 775 | 5,525 | 2,997 | 877 | 13,531 | 9,687 |
| Assembly in scaffolds > 50 Kb (%) | 90 | 35 | 99 | 57 | 72 | 94 | 93 | 1 | 44 | 76 |
| GC | 0.33 | 0.43 | 0.37 | 0.37 | 0.33 | 0.39 | 0.34 | 0.41 | 0.37 | 0.39 |
| Gap (%) | 8.4 | 0.3 | 0.1 | 0.0 | 2.5 | 18.7 | 16.0 | 0.0 | 0.3 | 3.1 |

**Table S3:** Repeat content of *C. augens* genome.

| Type | No. elements | Length (bp) | Percentage |
|---|---|---|---|
| **DNA transposons** | 338,259 | 89,947,607 | 7.980 |
| hAT-Ac | 105,066 | 27,773,842 | 2.464 |
| hAT-Tip100 | 70,523 | 18,910,018 | 1.678 |
| hAT-Blackjack | 39,085 | 11,083,906 | 0.983 |
| Crypton-I | 12,082 | 4,144,513 | 0.368 |
| RC | 17,021 | 3,827,203 | 0.340 |
| TcMar-Tc4 | 9,109 | 2,352,621 | 0.209 |
| hAT | 7,003 | 2,295,567 | 0.204 |
| hAT-Charlie | 8,210 | 2,229,463 | 0.198 |
| Sola-2 | 6,920 | 2,054,882 | 0.182 |
| TcMar-Mariner | 5,163 | 2,025,793 | 0.180 |
| hAT-Tag1 | 4,661 | 1,406,650 | 0.125 |
| Sola-1 | 7,536 | 1,269,310 | 0.113 |
| PIF-Harbinger | 5,513 | 1,225,180 | 0.109 |
| hAT-hATx | 2,798 | 997,184 | 0.088 |
| hAT-hAT5 | 3,947 | 839,482 | 0.074 |
| CMC-Chapaev-3 | 1,655 | 685,463 | 0.061 |
| MULE-MuDR | 2,586 | 675,591 | 0.060 |
| TcMar-Tc1 | 2,785 | 663,887 | 0.059 |
| CMC-EnSpm | 4,390 | 607,351 | 0.054 |
| PiggyBac | 1,480 | 605,286 | 0.054 |
| Merlin | 1,977 | 549,482 | 0.049 |
| MuLE-MuDR | 1,809 | 509,286 | 0.045 |
| CMC-Transib | 5,909 | 458,536 | 0.041 |
| Crypton | 1,900 | 442,963 | 0.039 |
| hAT-hAT19 | 1,298 | 437,367 | 0.039 |
| Crypton-V | 1,427 | 399,974 | 0.035 |
| IS3EU | 1,379 | 326,398 | 0.029 |
| Academ-1 | 1,502 | 316,671 | 0.028 |
| Maverick | 1,150 | 316,523 | 0.028 |
| Sola | 571 | 205,488 | 0.018 |
| TcMar-Tc2 | 1,204 | 151,704 | 0.013 |
| Kolobok-T2 | 375 | 125,298 | 0.011 |

| | | | |
|---|---|---|---|
| hAT-hATm | 106 | 23,23 | 0.002 |
| hAT-Pegasus | 45 | 4,211 | 0.000 |
| hAT-hobo | 36 | 3,702 | 0.000 |
| MULE-NOF | 23 | 2,079 | 0.000 |
| Zator | 12 | 1,36 | 0.000 |
| Kolobok-Hydra | 2 | 104 | 0.000 |
| TcMar-Pogo | 1 | 39 | 0.000 |
| **Retroelements** | 1,588,521 | 380,226,633 | 33.733 |
| **LINEs** | 96,449 | 25,730,249 | 2.283 |
| Penelope | 53,537 | 12,204,040 | 1.083 |
| L3 | 14,604 | 4,263,651 | 0.378 |
| CR2 | 10,535 | 3,625,019 | 0.322 |
| RTE-X | 3,636 | 1,669,624 | 0.148 |
| Jockey | 3,906 | 1,172,241 | 0.104 |
| L1-Tx2 | 4,752 | 963,984 | 0.086 |
| Dong-R5 | 1,041 | 559,161 | 0.050 |
| L2 | 741 | 468,809 | 0.042 |
| RTE-BovB | 1,381 | 356,411 | 0.032 |
| CR1-Zenon | 387 | 143,446 | 0.013 |
| I-Jockey | 1,169 | 142,842 | 0.013 |
| I-Nimb | 204 | 85,242 | 0.008 |
| CRE-II | 210 | 55,67 | 0.005 |
| R2 | 255 | 14,491 | 0.001 |
| R1-LOA | 39 | 2,209 | 0.000 |
| R3 | 35 | 1,826 | 0.000 |
| R2-NeSL | 17 | 1,583 | 0.000 |
| **SINEs** | 25,772 | 4,257,031 | 0.378 |
| tRNA | 11,654 | 1,680,388 | 0.149 |
| tRNA-Deu-RTE | 10,098 | 1,678,969 | 0.149 |
| MIR | 1,738 | 505,956 | 0.045 |
| tRNA-L3 | 1,419 | 263,956 | 0.023 |
| tRNA-V | 863 | 127,762 | 0.011 |
| **LTR elements** | 1,466,300 | 350,239,353 | 31.072 |
| Gypsy | 26,969 | 7,222,488 | 0.641 |
| Copia | 20,279 | 3,592,811 | 0.319 |
| Pao | 2,101 | 932,997 | 0.083 |

| | | | |
|---|---|---|---|
| Ngaro | 1,378 | 569,249 | 0.051 |
| ERVL | 2,219 | 485,597 | 0.043 |
| ERV1 | 1,424 | 350,196 | 0.031 |
| **Unknown** | 2002392 | 376811916 | 33.43 |
| **Low complexity** | 86 | 27,471 | 0.002 |
| **Satellites** | 2,644 | 631,963 | 0.056 |
| **Simple repeats** | 28,426 | 4,938,721 | 0.438 |
| **Total repeats** | | 475,772,395 | 45.734 |

**Table S4:** Gene composition of *C. augens* compared to the other ancestrally wingless hexapods.

| | Diplura | | Collembola | | |
|---|---|---|---|---|---|
| Metric | *Campodea augens* | *Catajapyx aquilonaris* | *Folsomia candida* | *Orchesella cincta* | *Holacanthella duospinosa* |
| Assembly size (Mb) | 1130 | 312 | 221 | 286 | 327 |
| No. of protein-coding genes | 23,992 | 10,901 | 22,100 | 20,247 | 9,895 |
| Total gene length (Mb) | 338 | 39 | 94 | 61 | 57 |
| Genome covered by genes (%) | 30.0 | 12.6 | 42.3 | 21.1 | 17.3 |
| Mean gene length (bp) | 14,081 | 3,609 | 4,240 | 2,990 | 5,726 |
| Median gene length (bp) | 9,728 | 2,529 | 2,240 | 2,043 | 3,784 |
| Total size of exons (Mb) / (%) | 31.9 / (2.83) | 15.5 / (4.96) | 53.1 / (23.94) | 36.2 / (12.63) | 22.7 / (6.93) |
| Mean exon length (bp) | 227 | 250 | 256 | 306 | 285 |
| Total size of introns (Mb) / (%) | 306.2 / (27,17) | 24.0 / (7,68) | 81.5 / (36,77) | 24.5 / (8,55) | 34.1 / (10.41) |
| Mean intron length (bp) | 2631 | 468 | 449 | 250 | 489 |
| Median intron length (bp) | 1636 | 128 | 88 | 90 | 107 |
| Mean exons per gene | 6 | 6 | 8 | 6 | 8 |
| Mean introns per gene | 5 | 5 | 7 | 5 | 7 |

*Intron and exon sizes were calculated based on the GFF files of the corresponding genomes (see Table SX for data sources).

**Table S5:** BUSCO analysis of *C. augens* assembly and gene set along with busco scores for the other available ancestrally wingless hexapods. BUSCO v3 was run using arthropoda_odb9 dataset.

| Species | Complete + Fragmented | Complete | Complete and single-copy | Complete and duplicated | Fragmented | Missing |
|---|---|---|---|---|---|---|
| *Campodea augens* | 97.6 | 91.8 | 89.8 | 2.0 | 5.8 | 2.4 |
| | 97.0 | 90.4 | 86.6 | 3.8 | 6.6 | 3.0 |
| *Catajapyx aquilonaris* | 99.3 | 97.0 | 94.9 | 2.1 | 2.3 | 0.7 |
| | 96.9 | 92.3 | 90.2 | 2.1 | 4.6 | 3.1 |
| *Folsomia candida* | 97.2 | 96.1 | 94.7 | 1.4 | 1.1 | 2.8 |
| | 98.6 | 97.8 | 72.0 | 2.2 | 0.8 | 1.4 |
| *Orchesella cincta* | 96.6 | 95.4 | 87.4 | 8.0 | 1.2 | 3.4 |
| | 92.2 | 88.5 | 81.6 | 6.9 | 3.7 | 7.8 |
| *Holacanthella duospinosa* | 83.9 | 80.4 | 73.6 | 6.8 | 3.5 | 16.1 |
| | 83.3 | 76.7 | 69.4 | 7.3 | 6.6 | 16.7 |

*Grey lines refers to busco scores for gene sets, blank lines to scores for genomes.

**Table S6:** Bacterial hits in *C. augens* genome.

| | Scaffold | HGT candidate | Start | Stop | Strand | Best hit gene ID | Clade | Species | Description |
|---|---|---|---|---|---|---|---|---|---|
| 1 | Scaffold_283 | CAUGE_02727 | 69101 | 71989 | - | gi\|1127377281\|gb\|APR98680.1\| | a-proteobacteria | Wolbachia endosymbiont of Folsomia candida | hypothetical protein ASM33_05545 [Wolbachia endosymbiont of Folsomia candida] |
| 2 | Scaffold_283 | CAUGE_02728 | 81704 | 85936 | - | gi\|1127377281\|gb\|APR98680.1\| | a-proteobacteria | Wolbachia endosymbiont of Folsomia candida | hypothetical protein ASM33_05545 [Wolbachia endosymbiont of Folsomia candida] |
| 3 | Scaffold_364 | CAUGE_03437 | 144611 | 182926 | - | gi\|736445120\|ref\|WP_034466997.1\| | a-proteobacteria | Afipia sp. P52-10 | CoA transferase subunit B [Afipia sp. P52-10] |
| 4 | Scaffold_433 | CAUGE_03961 | 57354 | 79310 | - | gi\|505401443\|ref\|WP_015588545.1\| | a-proteobacteria | Wolbachia endosymbiont of Drosophila simulans | Ankyrin repeat domain protein [Wolbachia endosymbiont of Drosophila simulans] |
| 5 | Scaffold_501 | CAUGE_04532 | 228425 | 236587 | - | gi\|1267475813\|ref\|WP_098291706.1\| | firmicutes | Bacillus thuringiensis | hypothetical protein [Bacillus thuringiensis] |
| 6 | Scaffold_516 | CAUGE_04651 | 609623 | 630080 | + | gi\|662720153\|gb\|AIE61363.1\| | firmicutes | Bacillus methanolicus MGA3 | hypothetical protein BMMGA3_15030 [Bacillus methanolicus MGA3] |
| 7 | Scaffold_550 | CAUGE_04872 | 44396 | 59277 | + | gi\|1169884537\|ref\|WP_080518662.1\| | b-proteobacteria | Burkholderia mallei | hypothetical protein [Burkholderia mallei] |
| 8 | Scaffold_1016 | CAUGE_07765 | 285612 | 315167 | + | gi\|1131919362\|emb\|SIT08514.1\| | a-proteobacteria | Insolitispirillum peregrinum | Ca2+-binding protein, RTX toxin-related [Insolitispirillum peregrinum] |
| 9 | Scaffold_1250 | CAUGE_08970 | 230 | 5447 | + | gi\|503535025\|ref\|WP_013769101.1\| | CFB group bacteria | Haliscomenobacter hydrossis | hypothetical protein [Haliscomenobacter hydrossis] |
| 10 | Scaffold_1281 | CAUGE_09116 | 101679 | 102350 | + | gi\|1154081569\|ref\|WP_078460720.1\| | g-proteobacteria | Solemya velum gill symbiont | hypothetical protein [Solemya velum gill symbiont] |
| 11 | Scaffold_1309 | CAUGE_09231 | 342507 | 346766 | + | gi\|1127377281\|gb\|APR98680.1\| | a-proteobacteria | Wolbachia endosymbiont of Folsomia candida | hypothetical protein ASM33_05545 [Wolbachia endosymbiont of Folsomia candida] |
| 12 | Scaffold_1309 | CAUGE_09232 | 371494 | 373902 | + | gi\|1244309946\|gb\|PBQ26533.1\| | a-proteobacteria | Wolbachia pipientis wAus | hypothetical protein BTO27_04595, partial [Wolbachia pipientis wAus] |
| 13 | Scaffold_1309 | CAUGE_09233 | 401769 | 405515 | + | gi\|1031777381\|ref\|WP_064125268.1\| | a-proteobacteria | Wolbachia endosymbiont of Dactylopius coccus | hypothetical protein [Wolbachia endosymbiont of Dactylopius coccus] |
| 14 | Scaffold_1587 | CAUGE_10534 | 27565 | 28146 | + | gi\|1200292621\|gb\|OUU14650.1\| | a-proteobacteria | Alphaproteobacteria bacterium TMED37 | hypothetical protein CBB97_24755 [Candidatus Endolissoclinum sp. TMED37] |

| | | | | | | | | | |
|---|---|---|---|---|---|---|---|---|---|
| 15 | Scaffold_1587 | CAUGE_10535 | 31594 | 32148 | + | gi\|1200292621\|gb\|OUU14650.1\| | a-proteobacteria | Alphaproteobacteria bacterium TMED37 | hypothetical protein CBB97_24755 [Candidatus Endolissoclinum sp. TMED37] |
| 16 | Scaffold_1587 | CAUGE_10536 | 34910 | 35467 | + | gi\|1200292621\|gb\|OUU14650.1\| | a-proteobacteria | Alphaproteobacteria bacterium TMED37 | hypothetical protein CBB97_24755 [Candidatus Endolissoclinum sp. TMED37] |
| 17 | Scaffold_1633 | CAUGE_10756 | 299421 | 307144 | + | gi\|981458280\|ref\|WP_059667283.1\| | b-proteobacteria | Burkholderia ubonensis | hypothetical protein [Burkholderia ubonensis] |
| 18 | Scaffold_3169 | CAUGE_15978 | 107069 | 144500 | - | gi\|1127377281\|gb\|APR98680.1\| | a-proteobacteria | Wolbachia endosymbiont of Folsomia candida | hypothetical protein ASM33_05545 [Wolbachia endosymbiont of Folsomia candida] |
| 19 | Scaffold_3228 | CAUGE_16137 | 15620 | 19066 | + | gi\|1180269025\|ref\|WP_083787830.1\| | a-proteobacteria | Rickettsia endosymbiont of Ixodes scapularis | hypothetical protein [Rickettsia endosymbiont of Ixodes scapularis] |
| 20 | Scaffold_3582 | CAUGE_16967 | 327 | 5679 | + | gi\|494597110\|ref\|WP_007355367.1\| | cyanobacteria | Oscillatoria sp. PCC 6506;Kamptonema | MULTISPECIES: hypothetical protein [Kamptonema] |
| 21 | Scaffold_3959 | CAUGE_17788 | 57789 | 60155 | + | gi\|1127377281\|gb\|APR98680.1\| | a-proteobacteria | Wolbachia endosymbiont of Folsomia candida | hypothetical protein ASM33_05545 [Wolbachia endosymbiont of Folsomia candida] |
| 22 | Scaffold_4330 | CAUGE_18555 | 1289 | 11179 | + | gi\|501450016\|ref\|WP_012473465.1\| | CFB group bacteria | Candidatus Amoebophilus asiaticus | hypothetical protein [Candidatus Amoebophilus asiaticus] |
| 23 | Scaffold_4450 | CAUGE_18804 | 41770 | 42873 | + | gi\|1234008076\|ref\|WP_094649383.1\| | a-proteobacteria | Rickettsia endosymbiont of Culicoides newsteadi | translational GTPase TypA [Rickettsia endosymbiont of Culicoides newsteadi] |
| 24 | Scaffold_6069 | CAUGE_20726 | 5785 | 6549 | - | gi\|1127377281\|gb\|APR98680.1\| | a-proteobacteria | Wolbachia endosymbiont of Folsomia candida | hypothetical protein ASM33_05545 [Wolbachia endosymbiont of Folsomia candida] |
| 25 | Scaffold_18317 | CAUGE_23038 | 28848 | 35599 | - | gi\|942689125\|ref\|WP_055393802.1\| | CFB group bacteria | Flagellimonas eckloniae | collagen-like protein [Flagellimonas eckloniae] |
| 26 | Scaffold_18509 | CAUGE_23530 | 97286 | 106692 | - | gi\|1084511163\|gb\|OGN73343.1\| | bacteria | Chlamydiae bacterium RIFCSPLOWO2_12_FULL_49_12 | hypothetical protein A3G30_02530 [Chlamydiae bacterium RIFCSPLOWO2_12_FULL_49_12] |
| 27 | Scaffold_18672 | CAUGE_23831 | 107631 | 108875 | - | gi\|488851699\|ref\|WP_002764105.1\| | spirochetes | Leptospira sp. 200901116 | hypothetical protein [Leptospira mayottensis] |
| 28 | Scaffold_172 | CAUGE_01697 | 183982 | 226051 | - | gi\|857977947\|emb\|CEO95922.1\| | cercozoans | Plasmodiophora brassicae | hypothetical protein PBRA_004612 [Plasmodiophora brassicae] |
| 29 | Scaffold_1009 | CAUGE_07722 | 198811 | 205880 | - | gi\|914546906\|gb\|KOB60119.1\| | apicomplexans | Plasmodium falciparum HB3 | hypothetical protein PFHG_01882 [Plasmodium falciparum HB3] |

**Table S7:** Characteristics of *C. augens* mitogenome.

| Name | Start | Stop | Strand | Length (bp) | ovl/nc | Start/Stop Codons | Length in *C. fragilis* | *Length in C. lubbocki* | *Length in D. pulex* |
|---|---|---|---|---|---|---|---|---|---|
| trnI(gat) | 1 | 62 | + | 62 | -3 | | 62 | 61 | 64 |
| trnQ(ttg) | 60 | 126 | - | 67 | 15 | | 65 | 64 | 68 |
| trnM(cat) | 142 | 205 | + | 64 | -12 | | 64 | 64 | 64 |
| nad2 | 194 | 1213 | + | 1020 | -2 | ATT/TAA | 1005 | 1005 | 988 |
| trnW(tca) | 1212 | 1277 | + | 66 | -1 | | 64 | 66 | 66 |
| trnC(gca) | 1277 | 1337 | - | 61 | 0 | | 62 | 53 | 64 |
| trnY(gta) | 1338 | 1398 | - | 61 | -2 | | 63 | 62 | 64 |
| cox1 | 1397 | 2935 | + | 1539 | -5 | TTG/TAA | 1540 | 1542 | 1538 |
| trnL2(taa) | 2931 | 2992 | + | 62 | -21 | | 64 | 63 | 68 |
| cox2 | 2972 | 3656 | + | 685 | 15 | ATA/T(AA) | 679 | 684 | 679 |
| trnK(ctt) | 3672 | 3737 | + | 66 | -2 | | 57 | 68 | 70 |
| trnD(gtc) | 3736 | 3797 | + | 62 | 0 | | 63 | 61 | 65 |
| atp8 | 3798 | 3953 | + | 156 | -7 | ATA/TAA | 156 | 156 | 162 |
| atp6 | 3947 | 4621 | + | 675 | 3 | ATG/TAA | 675 | 675 | 674 |
| cox3 | 4625 | 5411 | + | 787 | 0 | ATG/T(AA) | 787 | 787 | 789 |
| trnG(tcc) | 5412 | 5470 | + | 59 | 6 | | 59 | 60 | 61 |
| nad3 | 5477 | 5824 | + | 348 | -2 | ATA/TAG | 357 | 347 | 353 |
| trnA(tgc) | 5823 | 5890 | + | 68 | 0 | | 62 | 60 | 66 |
| trnR(tcg) | 5891 | 5941 | + | 51 | -3 | | 52 | 53 | 65 |
| trnN(gtt) | 5939 | 5999 | + | 61 | -2 | | 62 | 61 | 67 |
| trnS1(gct) | 5998 | 6052 | + | 55 | -1 | | 54 | 55 | 65 |

| | | | | | | | | | |
|---|---|---|---|---|---|---|---|---|---|
| trnE(ttc) | 6052 | 6114 | + | 63 | -2 | | 64 | 61 | 68 |
| trnF(gaa) | 6113 | 6174 | - | 62 | -1 | | 61 | 60 | 66 |
| nad5 | 6174 | 7880 | - | 1707 | 0 | ATT/TAG | 1707 | 1710 | 1708 |
| trnH(gtg) | 7881 | 7942 | - | 62 | -17 | | 64 | 60 | 64 |
| nad4 | 7926 | 9266 | - | 1341 | -7 | ATG/TAA | 1338 | 1326 | 1321 |
| nad4l | 9260 | 9544 | - | 285 | 5 | ATG/TAA | 285 | 285 | 276 |
| trnT(tgt) | 9550 | 9610 | + | 61 | 0 | | 60 | 61 | 65 |
| trnP(tgg) | 9611 | 9677 | - | 67 | 5 | | 67 | 63 | 65 |
| nad6 | 9683 | 10189 | + | 507 | -19 | ATT/TAA | 510 | 524 | 513 |
| cob | 10288 | 11373 | + | 1086 | -47 | ATG/TAG | 1143 | 1140 | 1134 |
| trnS2(tga) | 11327 | 11379 | + | 53 | 14 | | 56 | 55 | 69 |
| nad1 | 11427 | 12338 | - | 912 | -32 | ATA/TAG | 924 | 921 | 936 |
| trnL1(tag) | 12354 | 12414 | - | 61 | 33 | | 63 | 59 | 67 |
| rrnL | 12419 | 13460 | - | 1041 | | | 1093 | 1096 | 1314 |
| trnV(tac) | 13508 | 13567 | - | 60 | 602 | | 62 | 61 | 72 |
| rrnS | 13575 | 14258 | - | 683 | | | 722 | 738 | 753 |
| Putative Control Region | 14648 | 15673 | + | 1025 | | | 557 | 620 | 689 |

**Table S8:** Significant gene family contractions and expansions in *C. augens*.
"Family size change" corresponds to number of significantly expanded (+) and contracted (-) gene families (Viterbi p-value < 0.01), numbers in parentheses correspond to family size at the most recent common ancestor and family size in *C. augens*. Domain and domain description correspond to the most abundant domain which occurs in proteins of the cluster. Number of genes is the number of proteins in the cluster, with values in parentheses indicating the number of proteins which contains the corresponding most abundant domain.

| Cluster | Family size change | Most abundant domain | Domain description of the most abundant domain | Number of genes |
|---|---|---|---|---|
| 2960 | +5 (1 -> 6) | PF00027 | Cyclic nucleotide-binding domain | 19 (15) |
| 5412 | +6 (4 -> 10) | PF02801 | Beta-ketoacyl synthase, C-terminal domain | 66 (55) |
| 8640 | +15 (14 -> 29) | PF00005 | ABC transporter | 13 (134) |
| 13331 | +36 (14 -> 50) | PF00083 | Sugar (and other) transporter | 13 (122) |
| 17578 | +12 (8 -> 20) | PF00732 | GMC oxidoreductase | 69 (69) |
| 27531 | +10 (4 -> 14) | PF06974 | Protein of unknown function (DUF1298) | 92 (69) |
| 27628 | +11 (7 -> 18) | PF00149 | Calcineurin-like phosphoesterase | 676 (64) |
| 33010 | +60 (5 -> 65) | PF00651 | BTB/POZ domain | 757 (74) |
| 33393 | +4 (2 -> 6) | PF04937 | Protein of unknown function (DUF 659) | 50 (34) |
| 104662 | +15 (7 -> 22) | PF00083 | Sugar (and other) transporter | 76 (75) |
| 39781 | +33 (4 -> 37) | PF00060 | Ligand-gated ion channel (Ionotropic glutamate receptor) | 50 (48) |
| 40422 | +11 (14 -> 25) | PF04083 | Partial alpha/beta-hydrolase lipase region | 227 (173) |
| 42033 | +17 (4 -> 21) | PF00135 | Carboxylesterase family | 52 (52) |
| 44392 | +16 (5 -> 21) | PF00501 | AMP-binding enzyme | 52 (52) |
| 47035 | +40 (26 -> 66) | PF00067 | Cytochrome P450 | 357 (355) |
| 47488 | +8 (8 -> 16) | PF07690 | Major Facilitator Superfamily | 109 (94) |
| 49279 | +4 (1 -> 5) | PF00083 | Sugar (and other) transporter | 33 (32) |
| 49470 | +16 (7 -> 23) | PF00201 | UDP-glucoronosyl and UDP-glucosyl transferase | 97 (97) |
| 49474 | +5 (2 -> 7) | PF13516 | Leucine Rich repeat | 40 (4) |
| 49785 | +22 (20 -> 42) | PF00067 | Cytochrome P450 | 255 (255) |
| 51218 | +8 (6 -> 14) | PF00232 | Glycosyl hydrolase family 1 | 101 (101) |
| 53356 | +30 (10 -> 40) | PF00939 | Sodium:sulfate symporter transmembrane region | 91 (88) |
| 53444 | +39 (14 -> 53) | PF00201 | UDP-glucoronosyl and UDP-glucosyl transferase | 193 (190) |
| 53491 | +10 (2 -> 12) | PF14291 | Domain of unknown function (DUF4371) | 83 (19) |
| 53706 | +5 (3 -> 8) | PF00651 | BTB/POZ domain | 51 (50) |
| 53800 | +16 (4 -> 20) | PF00653 | Inhibitor of Apoptosis domain | 54 (51) |

| | | | | |
|---|---|---|---|---|
| 56848 | +19 (5 -> 24) | PF00656 | Caspase domain | 51 (48) |
| 58929 | +27 (5 -> 32) | PF08395 | 7tm Chemosensory receptor | 79 (1) |
| 59208 | +19 (4 -> 23) | PF02932 | Neurotransmitter-gated ion-channel transmembrane region | 35 (32) |
| 59400 | +7 (4 -> 11) | PF06151 | Trehalose receptor | 55 (43) |
| 60485 | +8 (3 -> 11) | PF00060 | Ligand-gated ion channel (Ionotropic glutamate receptor) | 45 (18) |
| 63073 | +11 (6 -> 17) | PF00089 | Trypsin | 51 (50) |
| 64020 | +14 (6 -> 20) | PF05577 | Serine carboxypeptidase S28 | 66 (66) |
| 64853 | +5 (3 -> 8) | PF00665 | Integrase core domain | 65 (1) |
| 66494 | +15 (5 -> 20) | PF00078 | Reverse transcriptase (RNA-dependent DNA polymerase) | 89 (22) |
| 68788 | +23 (8 -> 31) | PF00487 | Fatty acid desaturase | 112 (106) |
| 75433 | +7 (6 -> 13) | PF00026 | Eukaryotic aspartyl protease | 50 (50) |
| 82070 | +25 (2 -> 27) | PF00651 | BTB/POZ domain | 33 (28) |
| 83501 | +13 (7 -> 20) | PF00450 | Serine carboxypeptidase | 82 (82) |
| 84393 | -5 (5 -> 0) | PF00067 | Cytochrome P450 | 92 (92) |
| 86472 | +5 (3 -> 8) | PF00328 | Histidine phosphatase superfamily (branch 2) | 40 (34) |
| 89385 | +32 (3 -> 35) | PF00075 | RNase H | 48 (45) |
| 91160 | +8 (2 -> 10) | PF00650 | CRAL/TRIO domain | 38 (31) |
| 94341 | +36 (15 -> 51) | PF13358 | DDE superfamily endonuclease | 195 (86) |
| 95946 | +16 (6 -> 22) | PF07690 | Major Facilitator Superfamily | 53 (48) |
| 97696 | +14 (8 -> 22) | PF05699 | hAT family C-terminal dimerisation region | 127 (2) |
| 97817 | +16 (6 -> 22) | PF00078 | Reverse transcriptase (RNA-dependent DNA polymerase) | 129 (68) |
| 97889 | +34 (8 -> 42) | PF09588 | YqaJ-like viral recombinase domain | 107 (57) |
| 98186 | +29 (11 -> 40) | PF02958 | Ecdysteroid kinase | 120 (117) |
| 98261 | +15 (7 -> 22) | PF00650 | CRAL/TRIO domain | 87 (85) |
| 99868 | +19 (11 -> 30) | PF00650 | CRAL/TRIO domain | 151 (137) |
| 100866 | +9 (3 -> 12) | PF03055 | Retinal pigment epithelial membrane protein | 52 (52) |
| 101599 | +9 (2 -> 11) | PF06585 | Haemolymph juvenile hormone binding protein (JHBP) | 58 (57) |
| 39034 | +4 (1 -> 5) | PF05444 | Protein of unknown function (DUF753) | 83 (1) |
| 108430 | +9 (4 -> 13) | PF13639 | Ring finger domain | 52 (37) |
| 109223 | +7 (6 -> 13) | PF09588 | YqaJ-like viral recombinase domain | 83 (41) |
| 109614 | +4 (2 -> 6) | PF00027 | Cyclic nucleotide-binding domain | 30 (20) |
| 111466 | +11 (4 -> 15) | PF00106 | short chain dehydrogenase | 57 (52) |
| 111645 | +3 (1 -> 4) | PF11901 | Protein of unknown function (DUF3421) | 24 (23) |

| | | | | |
|---|---|---|---|---|
| 112386 | +26 (12 -> 38) | PF00858 | Amiloride-sensitive sodium channel | 187 (177) |
| 112624 | +15 (4 -> 19) | PF03184 | DDE superfamily endonuclease | 76 (49) |
| 115371 | +9 (5 -> 14) | PF00474 | Sodium:solute symporter family | 47 (45) |
| 117797 | +10 (10 -> 20) | PF00135 | Carboxylesterase family | 138 (138) |
| 118905 | +31 (9 -> 40) | PF00059 | Lectin C-type domain | 88 (77) |
| 122860 | +22 (4 -> 26) | PF02995 | Protein of unknown function (DUF229) | 67 (67) |
| 122921 | +17 (5 -> 22) | PF01130 | CD36 family | 61 (61) |
| 123580 | +4 (2 -> 6) | PF00589 | Phage integrase family | 39 (0) |
| 123691 | +6 (1 -> 7) | PF00043 | Glutathione S-transferase, C-terminal domain | 29 (17) |
| 124181 | +14 (7 -> 21) | PF00188 | Cysteine-rich secretory protein family | 75 (57) |
| 125861 | +17 (9 -> 26) | PF00892 | EamA-like transporter family | 109 (101) |
| 127068 | +10 (4 -> 14) | PF00106 | short chain dehydrogenase | 78 (75) |
| 127634 | +28 (10 -> 38) | PF05050 | Methyltransferase FkbM domain | 62 (57) |
| 127702 | +6 (2 -> 8) | PF13359 | DDE superfamily endonuclease | 32 (19) |
| 131842 | +3 (1 -> 4) | PF09588 | YqaJ-like viral recombinase domain | 20 (17) |
| 131847 | +19 (3 -> 22) | PF00061 | Lipocalin / cytosolic fatty-acid binding protein family | 38 (5) |
| 135298 | +9 (1 -> 10) | PF00060 | Ligand-gated ion channel (Ionotropic glutamate receptor) | 36 (6) |
| 141959 | +8 (1 -> 9) | PF00043 | Glutathione S-transferase, C-terminal domain | 26 (10) |
| 143375 | +8 (6 -> 14) | PF00628 | PHD-finger | 143 (97) |
| 145937 | +3 (1 -> 4) | PF00125 | Core histone H2A/H2B/H3/H4 | 29 (2) |
| 147915 | +31 (11 -> 42) | PF14223 | gag-polypeptide of LTR copia-type | 225 (77) |
| 150999 | +16 (9 -> 25) | PF16211 | C-terminus of histone H2A | 184 (112) |
| 153470 | +29 (9 -> 38) | PF00125 | Core histone H2A/H2B/H3/H4 | 131 (118) |
| 155603 | +24 (10 -> 34) | PF15511 | Centromere kinetochore component CENP-T histone fold | 138 (138) |
| 158103 | +8 (4 -> 12) | PF14497 | Glutathione S-transferase, C-terminal domain | 75 (28) |
| 158468 | +10 (9 -> 19) | PF00644 | Poly(ADP-ribose) polymerase catalytic domain | 159 (151) |
| 158935 | +21 (3 -> 24) | PF13975 | gag-polyprotein putative aspartyl protease | 49 (41) |
| 159658 | +4 (2 -> 6) | PF14291 | Domain of unknown function (DUF4371) | 29 (4) |
| 160485 | +3 (1 -> 4) | PF05485 | THAP domain | 19 (11) |
| 166707 | +4 (2 -> 6) | PF00538 | linker histone H1 and H5 family | 32 (29) |
| 169474 | +7 (5 -> 12) | PF09005 | Domain of unknown function (DUF1897) | 90 (90) |
| 171004 | +16 (2 -> 18) | - | - | - |

**Table S9:** Gene families in *C. augens* that are significantly larger than the mean counts in insects.

| clid | PFAM | description | *Dpul* | *Apis* | *Tcas* | *Dmel* | *Amel* | *Dple* | *Ocin* | *Cspl* | *Phum* | *Smar* | *Fcan* | *Caug* | *Caqu* | Mean (Insecta) | P |
|---|---|---|---|---|---|---|---|---|---|---|---|---|---|---|---|---|---|
| 56848 | PF08395 | 7tm Chemosensory receptor | 1 | 3 | 4 | 1 | 1 | 1 | 6 | 2 | 1 | 3 | 2 | **24** | 2 | 1.9 | 6.72E-05 |
| 166091 | PF13304 | AAA domain, putative AbiEii toxin, Type IV TA system | 0 | 0 | 0 | 0 | 0 | 0 | 0 | 0 | 0 | 0 | 0 | **7** | 0 | 0.0 | 8.49E-03 |
| 8640 | PF00005 | ABC transporter | 4 | 5 | 11 | 10 | 3 | 5 | 22 | 6 | 2 | 9 | 16 | **29** | 14 | 6.0 | 3.19E-04 |
| 44392 | PF00501 | AMP-binding enzyme | 2 | 1 | 1 | 2 | 1 | 1 | 2 | 0 | 1 | 15 | 1 | **21** | 4 | 1.0 | 8.16E-05 |
| 109398 | PF16076 | Acyltransferase C-terminus | 1 | 0 | 1 | 6 | 1 | 1 | 2 | 1 | 1 | 1 | 1 | **12** | 1 | 1.6 | 5.67E-03 |
| 127634 | PF01425 | Amidase | 1 | 1 | 1 | 2 | 1 | 1 | 1 | 1 | 1 | 2 | 4 | **38** | 10 | 1.1 | 1.00E-07 |
| 112235 | PF00858 | Amiloride-sensitive sodium channel | 2 | 1 | 1 | 2 | 1 | 1 | 2 | 2 | 1 | 0 | 3 | **17** | 2 | 1.3 | 6.12E-04 |
| 53899 | PF00155 | Aminotransferase class I and II | 1 | 2 | 2 | 2 | 2 | 2 | 3 | 2 | 2 | 2 | 2 | **14** | 3 | 2.0 | 3.70E-03 |
| 133240 | PF12796 | Ankyrin repeats (3 copies) | 1 | 0 | 1 | 1 | 0 | 1 | 1 | 1 | 1 | 0 | 1 | **11** | 1 | 0.7 | 3.70E-03 |
| 68928 | PF07530 | Associated with zinc fingers | 0 | 2 | 0 | 0 | 0 | 0 | 0 | 0 | 0 | 0 | 0 | **8** | 0 | 0.3 | 8.28E-03 |
| 33010 | PF00651 | BTB/POZ domain | 1 | 0 | 1 | 1 | 0 | 1 | 1 | 1 | 1 | 1 | 1 | **65** | 1 | 0.7 | 0.00E+00 |
| 78766 | PF00651 | BTB/POZ domain | 2 | 4 | 1 | 1 | 2 | 2 | 2 | 2 | 1 | 3 | 2 | **14** | 4 | 1.9 | 3.32E-03 |
| 118905 | PF00651 | BTB/POZ domain | 2 | 0 | 0 | 6 | 0 | 3 | 14 | 10 | 0 | 1 | 6 | **40** | 6 | 2.7 | 1.00E-07 |
| 150999 | PF16211 | C-terminus of histone H2A | 83 | 43 | 0 | 0 | 0 | 1 | 2 | 3 | 0 | 27 | 0 | **25** | 0 | 6.7 | 2.13E-03 |
| 122860 | PF01130 | CD36 family | 0 | 1 | 8 | 9 | 1 | 2 | 5 | 2 | 1 | 2 | 9 | **26** | 1 | 3.4 | 1.14E-04 |
| 98186 | PF00650 | CRAL/TRIO domain | 9 | 1 | 19 | 5 | 2 | 7 | 8 | 6 | 2 | 2 | 13 | **40** | 7 | 6.0 | 4.50E-06 |
| 98261 | PF00650 | CRAL/TRIO domain | 5 | 5 | 7 | 4 | 5 | 8 | 5 | 4 | 6 | 5 | 10 | **22** | 1 | 5.6 | 2.78E-03 |
| 27628 | PF00149 | Calcineurin-like phosphoesterase | 4 | 1 | 9 | 3 | 2 | 2 | 4 | 3 | 5 | 3 | 7 | **18** | 6 | 3.6 | 2.94E-03 |

| | | | | | | | | | | | | | | | | | |
|---|---|---|---|---|---|---|---|---|---|---|---|---|---|---|---|---|---|
| 63073 | PF00149 | Calcineurin-like phosphoesterase | 0 | 3 | 3 | 5 | 2 | 5 | 0 | 5 | 2 | 0 | 3 | **17** | 6 | 3.6 | 4.16E-03 |
| 64020 | PF08434 | Calcium-activated chloride channel N terminal | 2 | 7 | 2 | 6 | 3 | 5 | 2 | 5 | 2 | 2 | 6 | **20** | 4 | 4.3 | 2.47E-03 |
| 42033 | PF00135 | Carboxylesterase family | 6 | 0 | 0 | 4 | 0 | 0 | 4 | 2 | 1 | 0 | 14 | **21** | 0 | 1.0 | 8.16E-05 |
| 116570 | PF00135 | Carboxylesterase family | 1 | 1 | 1 | 1 | 1 | 1 | 1 | 1 | 1 | 0 | 1 | **15** | 3 | 1.0 | 1.03E-03 |
| 56529 | PF00656 | Caspase domain | 1 | 3 | 1 | 1 | 1 | 1 | 2 | 1 | 1 | 4 | 5 | **10** | 2 | 1.3 | 9.49E-03 |
| 153470 | PF15511 | Centromere kinetochore component CENP-T histone fold | 6 | 9 | 5 | 20 | 5 | 5 | 24 | 2 | 5 | 7 | 4 | **38** | 1 | 7.3 | 3.01E-05 |
| 40709 | PF00755 | Choline/Carnitine o-acyltransferase | 4 | 3 | 4 | 7 | 3 | 4 | 3 | 10 | 5 | 4 | 5 | **21** | 8 | 5.1 | 2.96E-03 |
| 143584 | PF00125 | Core histone H2A/H2B/H3/H4 | 1 | 2 | 0 | 0 | 0 | 0 | 0 | 0 | 0 | 0 | 0 | **10** | 0 | 0.3 | 3.47E-03 |
| 151496 | PF00125 | Core histone H2A/H2B/H3/H4 | 0 | 0 | 0 | 0 | 0 | 0 | 0 | 0 | 0 | 0 | 0 | **22** | 0 | 0.0 | 1.81E-05 |
| 124181 | PF00188 | Cysteine-rich secretory protein family | 5 | 12 | 3 | 11 | 1 | 3 | 3 | 3 | 2 | 2 | 4 | **21** | 5 | 5.0 | 2.75E-03 |
| 47035 | PF00067 | Cytochrome P450 | 11 | 23 | 46 | 25 | 24 | 24 | 44 | 19 | 8 | 10 | 46 | **66** | 11 | 24.1 | 5.46E-05 |
| 49785 | PF00067 | Cytochrome P450 | 17 | 5 | 4 | 3 | 5 | 5 | 73 | 17 | 4 | 8 | 58 | **42** | 14 | 6.1 | 2.30E-06 |
| 92010 | PF13358 | DDE superfamily endonuclease | 6 | 0 | 2 | 1 | 1 | 1 | 0 | 2 | 1 | 2 | 1 | **10** | 2 | 1.1 | 8.49E-03 |
| 100866 | PF13359 | DDE superfamily endonuclease | 2 | 1 | 1 | 1 | 1 | 3 | 16 | 2 | 0 | 3 | 11 | **12** | 1 | 1.3 | 4.32E-03 |
| 121414 | PF13359 | DDE superfamily endonuclease | 43 | 289 | 1 | 0 | 0 | 0 | 24 | 44 | 1 | 4 | 1 | **11** | 2 | 47.9 | 1.09E-05 |
| 155603 | PF00226 | DnaJ domain | 11 | 8 | 3 | 23 | 5 | 2 | 28 | 1 | 6 | 15 | 2 | **34** | 0 | 6.9 | 8.55E-05 |
| 164437 | PF09005 | Domain of unknown function (DUF1897) | 1 | 0 | 1 | 1 | 1 | 1 | 1 | 1 | 1 | 1 | 1 | **11** | 1 | 0.9 | 4.32E-03 |
| 99868 | PF14291 | Domain of unknown function (DUF4371) | 29 | 1 | 1 | 1 | 2 | 1 | 21 | 1 | 0 | 1 | 58 | **30** | 5 | 1.0 | 2.00E-06 |
| 120794 | PF16064 | Domain of unknown function (DUF4806) | 2 | 0 | 0 | 0 | 0 | 1 | 2 | 0 | 0 | 0 | 5 | **10** | 1 | 0.1 | 2.99E-03 |

| | | | | | | | | | | | | | | | | |
|---|---|---|---|---|---|---|---|---|---|---|---|---|---|---|---|---|
| 125861 | PF00892 | EamA-like transporter family | 12 | 0 | 22 | 17 | 1 | 7 | 5 | 3 | 2 | 8 | 1 | **26** | 5 | 7.4 | 2.34E-03 |
| 97889 | PF02958 | Ecdysteroid kinase | 3 | 35 | 1 | 1 | 0 | 0 | 1 | 19 | 0 | 0 | 0 | **42** | 5 | 8.0 | 1.09E-05 |
| 66242 | PF00487 | Fatty acid desaturase | 0 | 2 | 1 | 6 | 1 | 1 | 8 | 2 | 1 | 5 | 6 | **12** | 3 | 2.0 | 8.39E-03 |
| 191185 | PF00041 | Fibronectin type III domain | 0 | 0 | 0 | 0 | 0 | 0 | 0 | 0 | 0 | 0 | 0 | **7** | 0 | 0.0 | 8.49E-03 |
| 17578 | PF00732 | GMC oxidoreductase | 0 | 0 | 1 | 0 | 1 | 1 | 27 | 1 | 1 | 8 | 0 | **20** | 9 | 0.7 | 8.63E-05 |
| 134686 | PF17172 | Glutathione S-transferase N-terminal domain | 87 | 7 | 3 | 2 | 4 | 11 | 19 | 2 | 1 | 3 | 15 | **23** | 15 | 4.3 | 7.79E-04 |
| 122921 | PF00043 | Glutathione S-transferase, C-terminal domain | 4 | 1 | 4 | 6 | 2 | 4 | 7 | 3 | 4 | 1 | 1 | **22** | 2 | 3.4 | 6.05E-04 |
| 155892 | PF14497 | Glutathione S-transferase, C-terminal domain | 3 | 7 | 2 | 23 | 4 | 3 | 11 | 0 | 6 | 4 | 1 | **30** | 0 | 6.4 | 3.12E-04 |
| 158935 | PF01531 | Glycosyl transferase family 11 | 0 | 18 | 4 | 0 | 0 | 0 | 0 | 0 | 0 | 0 | 1 | **24** | 2 | 3.1 | 2.15E-04 |
| 83501 | PF13896 | Glycosyl-transferase for dystroglycan | 0 | 6 | 5 | 5 | 2 | 7 | 9 | 5 | 4 | 2 | 13 | **20** | 4 | 4.9 | 3.42E-03 |
| 117797 | PF06585 | Haemolymph juvenile hormone binding protein (JHBP) | 11 | 12 | 6 | 3 | 3 | 3 | 23 | 5 | 1 | 3 | 40 | **20** | 8 | 4.7 | 3.16E-03 |
| 53444 | PF00653 | Inhibitor of Apoptosis domain | 11 | 12 | 25 | 26 | 6 | 12 | 8 | 6 | 3 | 12 | 16 | **53** | 3 | 12.9 | 5.90E-06 |
| 115371 | PF00059 | Lectin C-type domain | 0 | 4 | 2 | 1 | 1 | 1 | 6 | 1 | 1 | 0 | 9 | **14** | 7 | 1.6 | 2.68E-03 |
| 118113 | PF00059 | Lectin C-type domain | 0 | 0 | 0 | 0 | 0 | 0 | 0 | 0 | 0 | 0 | 0 | **7** | 0 | 0.0 | 8.49E-03 |
| 39781 | PF00060 | Ligand-gated ion channel | 3 | 1 | 1 | 1 | 1 | 0 | 0 | 0 | 1 | 3 | 1 | **37** | 1 | 0.7 | 1.00E-07 |
| 53800 | PF00060 | Ligand-gated ion channel | 2 | 5 | 2 | 2 | 3 | 2 | 3 | 2 | 2 | 3 | 6 | **20** | 2 | 2.6 | 6.15E-04 |
| 137329 | PF00060 | Ligand-gated ion channel | 5 | 0 | 0 | 0 | 0 | 0 | 11 | 0 | 0 | 0 | 11 | **12** | 2 | 0.0 | 1.16E-03 |
| 58929 | PF00917 | MATH domain | 12 | 32 | 0 | 0 | 0 | 1 | 0 | 0 | 0 | 0 | 0 | **32** | 2 | 4.7 | 3.67E-05 |
| 94341 | PF07690 | Major Facilitator Superfamily | 0 | 76 | 5 | 0 | 0 | 2 | 1 | 32 | 1 | 2 | 15 | **51** | 10 | 16.6 | 1.04E-04 |
| 127655 | PF05050 | Methyltransferase FkbM domain | 2 | 0 | 0 | 0 | 0 | 0 | 9 | 0 | 0 | 2 | 5 | **13** | 2 | 0.0 | 7.27E-04 |
| 59077 | PF02932 | Neurotransmitter-gated ion- | 1 | 0 | 0 | 0 | 0 | 0 | 1 | 0 | 0 | 0 | 1 | **13** | 0 | 0.0 | 7.27E-04 |

| | | | | | | | | | | | | | | | | |
|---|---|---|---|---|---|---|---|---|---|---|---|---|---|---|---|---|
| | | channel transmembrane region | | | | | | | | | | | | | | |
| 68788 | PF01057 | Parvovirus non-structural protein NS1 | 5 | 9 | 13 | 6 | 7 | 11 | 8 | 3 | 5 | 3 | 11 | **31** | 1 | 7.7 | 4.99E-04 |
| 15709 | PF02460 | Patched family | 2 | 0 | 1 | 0 | 2 | 1 | 2 | 1 | 1 | 0 | 4 | **15** | 3 | 0.9 | 8.62E-04 |
| 157983 | PF00644 | Poly(ADP-ribose) polymerase catalytic domain | 1 | 0 | 0 | 0 | 0 | 0 | 0 | 0 | 0 | 0 | 1 | **8** | 0 | 0.0 | 5.67E-03 |
| 171004 | PF00644 | Poly(ADP-ribose) polymerase catalytic domain | 0 | 0 | 0 | 0 | 0 | 0 | 0 | 13 | 0 | 0 | 0 | **18** | 0 | 1.9 | 7.07E-04 |
| 160053 | PF00069 | Protein kinase domain | 1 | 0 | 1 | 0 | 1 | 0 | 2 | 6 | 1 | 3 | 6 | **10** | 8 | 1.3 | 9.49E-03 |
| 127736 | PF04937 | Protein of unknown function (DUF 659) | 2 | 0 | 0 | 0 | 0 | 1 | 0 | 2 | 0 | 0 | 0 | **11** | 0 | 0.4 | 2.78E-03 |
| 27531 | PF06974 | Protein of unknown function (DUF1298) | 0 | 1 | 1 | 0 | 1 | 1 | 22 | 2 | 1 | 13 | 35 | **14** | 1 | 1.0 | 1.60E-03 |
| 50974 | PF06974 | Protein of unknown function (DUF1298) | 1 | 1 | 1 | 1 | 1 | 1 | 6 | 1 | 1 | 2 | 8 | **12** | 1 | 1.0 | 3.32E-03 |
| 46593 | PF02995 | Protein of unknown function (DUF229) | 0 | 2 | 1 | 1 | 1 | 2 | 0 | 2 | 1 | 3 | 0 | **19** | 5 | 1.4 | 3.19E-04 |
| 65583 | PF02995 | Protein of unknown function (DUF229) | 13 | 1 | 1 | 0 | 0 | 0 | 5 | 1 | 1 | 7 | 8 | **12** | 4 | 0.6 | 2.28E-03 |
| 84519 | PF08373 | RAP domain | 1 | 1 | 1 | 1 | 1 | 1 | 2 | 2 | 2 | 2 | 3 | **14** | 1 | 1.3 | 2.12E-03 |
| 86631 | PF00075 | RNase H | 3 | 1 | 0 | 1 | 2 | 1 | 5 | 2 | 1 | 2 | 8 | **10** | 1 | 1.1 | 8.49E-03 |
| 147915 | PF00075 | RNase H | 2 | 66 | 2 | 0 | 0 | 0 | 1 | 28 | 0 | 83 | 0 | **42** | 1 | 13.7 | 4.32E-04 |
| 86362 | PF00355 | Rieske [2Fe-2S] domain | 1 | 1 | 1 | 1 | 0 | 0 | 1 | 1 | 0 | 0 | 1 | **16** | 1 | 0.6 | 4.32E-04 |
| 107419 | PF13639 | Ring finger domain | 55 | 212 | 5 | 0 | 0 | 3 | 3 | 60 | 0 | 6 | 2 | **15** | 0 | 40.0 | 1.55E-03 |
| 108430 | PF13639 | Ring finger domain | 1 | 5 | 1 | 3 | 2 | 1 | 14 | 1 | 1 | 2 | 6 | **13** | 2 | 2.0 | 5.67E-03 |
| 141366 | PF00856 | SET domain | 1 | 0 | 0 | 0 | 0 | 0 | 0 | 1 | 0 | 1 | 0 | **12** | 2 | 0.1 | 1.43E-03 |
| 139416 | PF12146 | Serine aminopeptidase, S33 | 5 | 0 | 1 | 2 | 1 | 1 | 1 | 4 | 1 | 3 | 6 | **12** | 6 | 1.4 | 5.09E-03 |

| | | | | | | | | | | | | | | | | |
|---|---|---|---|---|---|---|---|---|---|---|---|---|---|---|---|---|
| 82070 | PF00450 | Serine carboxypeptidase | 0 | 0 | 0 | 1 | 1 | 1 | 1 | 0 | 1 | 0 | 1 | **27** | 0 | 0.6 | 4.30E-06 |
| 63522 | PF05577 | Serine carboxypeptidase S28 | 1 | 1 | 1 | 1 | 1 | 2 | 1 | 1 | 2 | 2 | 1 | **12** | 1 | 1.3 | 4.32E-03 |
| 113596 | PF00474 | Sodium:solute symporter family | 16 | 0 | 0 | 0 | 0 | 0 | 8 | 0 | 0 | 0 | 0 | **8** | 0 | 0.0 | 5.67E-03 |
| 52512 | PF00939 | Sodium:sulfate symporter transmembrane region | 0 | 0 | 0 | 3 | 0 | 0 | 0 | 0 | 0 | 0 | 0 | **8** | 0 | 0.4 | 9.33E-03 |
| 7578 | PF00083 | Sugar (and other) transporter | 0 | 0 | 0 | 0 | 0 | 0 | 1 | 0 | 0 | 0 | 0 | **7** | 3 | 0.0 | 8.49E-03 |
| 13331 | PF00083 | Sugar (and other) transporter | 2 | 6 | 8 | 7 | 4 | 12 | 8 | 6 | 5 | 8 | 10 | **50** | 11 | 6.9 | 1.00E-07 |
| 52239 | PF00083 | Sugar (and other) transporter | 9 | 0 | 1 | 1 | 0 | 1 | 0 | 0 | 0 | 0 | 0 | **9** | 0 | 0.4 | 6.23E-03 |
| 135552 | PF00685 | Sulfotransferase domain | 0 | 1 | 1 | 1 | 0 | 1 | 0 | 0 | 1 | 0 | 0 | **13** | 1 | 0.7 | 1.79E-03 |
| 139968 | PF00335 | Tetraspanin family | 11 | 0 | 1 | 3 | 0 | 1 | 0 | 0 | 1 | 0 | 0 | **13** | 0 | 0.9 | 2.11E-03 |
| 215323 | PF00089 | Trypsin | 0 | 0 | 0 | 0 | 0 | 0 | 0 | 0 | 0 | 0 | 0 | **8** | 0 | 0.0 | 5.67E-03 |
| 220751 | PF00089 | Trypsin | 0 | 0 | 0 | 0 | 0 | 0 | 0 | 0 | 0 | 0 | 0 | **9** | 0 | 0.0 | 3.70E-03 |
| 89385 | PF01663 | Type I phosphodiesterase / nucleotide pyrophosphatase | 0 | 11 | 0 | 0 | 0 | 0 | 0 | 0 | 0 | 1 | 1 | **35** | 0 | 1.6 | 4.00E-07 |
| 49470 | PF00201 | UDP-glucoronosyl and UDP-glucosyl transferase | 7 | 16 | 5 | 4 | 2 | 9 | 9 | 1 | 0 | 1 | 19 | **23** | 1 | 5.3 | 1.73E-03 |
| 53356 | PF00201 | UDP-glucoronosyl and UDP-glucosyl transferase | 6 | 0 | 3 | 2 | 2 | 4 | 6 | 6 | 1 | 5 | 11 | **40** | 6 | 2.6 | 1.00E-07 |
| 215795 | PF02825 | WWE domain | 0 | 0 | 0 | 0 | 0 | 0 | 0 | 0 | 0 | 1 | 0 | **7** | 0 | 0.0 | 8.49E-03 |
| 127702 | PF09588 | YqaJ-like viral recombinase domain | 2 | 0 | 1 | 0 | 1 | 1 | 18 | 0 | 0 | 0 | 0 | **8** | 1 | 0.4 | 9.33E-03 |
| 104662 | PF11977 | Zc3h12a-like Ribonuclease NYN domain | 2 | 7 | 12 | 3 | 6 | 9 | 0 | 7 | 6 | 0 | 0 | **22** | 4 | 7.1 | 6.78E-03 |
| 42822 | PF00096 | Zinc finger, C2H2 type | 16 | 136 | 35 | 15 | 14 | 17 | 54 | 23 | 8 | 20 | 193 | **85** | 12 | 35.4 | 3.59E-05 |
| 117864 | PF00569 | Zinc finger, ZZ type | 0 | 2 | 2 | 3 | 1 | 1 | 0 | 1 | 2 | 0 | 2 | **11** | 0 | 1.7 | 9.36E-03 |
| 59244 | PF05699 | hAT family C-terminal dimerisation region | 0 | 0 | 0 | 0 | 0 | 0 | 0 | 2 | 6 | 1 | 0 | **11** | 0 | 1.1 | 5.67E-03 |

| | | | | | | | | | | | | | | | | | |
|---|---|---|---|---|---|---|---|---|---|---|---|---|---|---|---|---|---|
| 95946 | PF05699 | hAT family C-terminal dimerisation region | 4 | 2 | 8 | 1 | 2 | 1 | 1 | 1 | 1 | 2 | 2 | **22** | 6 | 2.3 | 2.15E-04 |
| 90160 | PF00106 | short chain dehydrogenase | 4 | 2 | 1 | 0 | 1 | 0 | 10 | 1 | 1 | 3 | 5 | **10** | 5 | 0.9 | 6.39E-03 |
| 127068 | PF00106 | short chain dehydrogenase | 1 | 1 | 1 | 1 | 2 | 1 | 22 | 1 | 2 | 2 | 28 | **14** | 2 | 1.3 | 2.12E-03 |

**Table S10:** Counts of Pfam domains associated with DNA photolyase in *C. augens* and 12 other arthropods.

| Domain | pfamID | *Smar** | *Dpul* | *Fcan** | *Ocin* | *Caqu** | *Caug** | *Cspl* | *Apis* | *Phum* | *Amel* | *Tcas* | *Dple* | *Dmel* |
|---|---|---|---|---|---|---|---|---|---|---|---|---|---|---|
| DNA photolyase | PF00875 | 0 | 8 | 0 | 6 | 0 | 0 | 5 | 6 | 1 | 2 | 1 | 4 | 3 |
| FAD binding domain of DNA photolyase | PF03441 | 0 | 4 | 0 | 4 | 0 | 0 | 3 | 5 | 1 | 1 | 1 | 3 | 2 |

* asterisks mark species which lack eyes.

**Table S11:** Counts of Pfam domains associated with the caspase domain in *C. augens* and 12 other arthropods.

| Domain | pfamID | *Smar* | *Dpul* | *Fcan* | *Ocin* | *Caqu* | *Caug* | *Cspl* | *Apis* | *Phum* | *Amel* | *Tcas* | *Dple* | *Dmel* |
|---|---|---|---|---|---|---|---|---|---|---|---|---|---|---|
| Caspase domain | PF00656 | 8 | 18 | 13 | 14 | 12 | 35 | 16 | 6 | 5 | 5 | 8 | 4 | 6 |

**Table S12:** Blast hit and Pfam domains of *C. augens* EVEs.

| Scaffold | Coordinates | nr best hit | e-value | Identity | Pfam hit |
|---|---|---|---|---|---|
| Scaffold_1050 | 14011-15100 | PB1 [Wuhan Mosquito Virus 3] | 3E-34 | 39% | PF00602 (Influenza RNA-dependent RNA polymerase subunit PB1) |
| Scaffold_6539 | 2499-3089 | PB1 [Wuhan Louse Fly Virus 3] | 3E-49 | 44% | PF00602 (Influenza RNA-dependent RNA polymerase subunit PB1) |
| Scaffold_12940 | 1460-1897 | PB1 [Wuhan Mosquito Virus 5] | 5E-48 | 51% | PF00602 (Influenza RNA-dependent RNA polymerase subunit PB1) |
| Scaffold_5450 | 15766-16765 | PB1 [Wuhan Louse Fly Virus 3] | 1E-25 | 43% | PF00602 (Influenza RNA-dependent RNA polymerase subunit PB1) |
| Scaffold_240 | 94436-94965 | PB1 [Wuhan Mosquito Virus 3] | 8E-32 | 42% | - |
| Scaffold_2171 | 26421-27126 | PB1 [Shuangao Insect Virus 4] | 2E-06 | 39% | - |
| Scaffold_2171 | 22958-23373 | polymerase basic 1 protein [Wellfleet Bay virus] | 3E-07 | 33% | - |

**Table S13:** Accession number of PB1 proteins from orthomyxoviruses used in the phylogenetic analysis of *C. augens* EVEs.

| Virus | PB1 Genbank Accesion |
|---|---|
| Influenzavirus A | BAU68348.1 |
| Influenzavirus B | NP_056657.1 |
| Influenzavirus C | YP_089653.1 |
| Influenzavirus D | AIE52114.1 |
| Salmon_isavirus | YP_145804.1 |
| Quaranfil quaranjavirus | ACY56282.1 |
| Johnston Atoll virus | ACY56284.1 |
| Wellfleet Bay virus | YP_009110686.1 |
| Aransas bay virus | AHB34061.1 |
| Dhori thogotovirus | ADF56030.1 |
| Thogoto thogotovirus | YP_145794.1 |
| Jingshan Fly Virus 1 | AJG39084.1 |
| Sanxia Water Strider Virus 3 | AJG39086.1 |
| Shuangao Insect Virus 4 | AJG39088.1 |
| Wuhan Louse Fly Virus 3 | AJG39089.1 |
| Wuhan Louse Fly Virus 4 | AJG39090.1 |
| Wuhan Mosquito Virus 3 | AJG39091.1 |
| Wuhan Mosquito Virus 4 | AJG39092.1 |
| Wuhan Mosquito Virus 5 | AJG39093.1 |
| Wuhan Mosquito Virus 6 | AJG39094.1 |
| Wuhan Mosquito Virus 7 | AJG39095.1 |
| Wuhan Mothfly Virus | AJG39096.1 |
| Sinu virus | APP91612.1 |

**Table S14:** Source of genomic data used in the analyses.

| Species | Fasta | gff | Protein |
|---|---|---|---|
| caqu | https://i5k.nal.usda.gov/sites/default/files/data/Arthropoda/cataqu-%28Catajapyx_aquilonaris%29/Current%20Genome%20Assembly/1.Genome%20Assembly/BCM-After-Atlas/Scaffolds/forcepstail.consistent.scaffolds.fa.gz | https://i5k.nal.usda.gov/sites/default/files/data/Arthropoda/cataqu-%28Catajapyx_aquilonaris%29/Current%20Genome%20Assembly/2.Official%20or%20Primary%20Gene%20Set/BCM_version_0.5.3/consensus_gene_set/CAQU.Models.gff3.gz | https://i5k.nal.usda.gov/sites/default/files/data/Arthropoda/cataqu-%28Catajapyx_aquilonaris%29/Current%20Genome%20Assembly/2.Official%20or%20Primary%20Gene%20Set/BCM_version_0.5.3/consensus_gene_set/CAQU.faa.gz |
| fcan | ftp://ftp.ncbi.nlm.nih.gov/genomes/refseq/invertebrate/Folsomia_candida/representative/GCF_002217175.1_ASM221717v1/GCF_002217175.1_ASM221717v1_genomic.fna.gz | ftp://ftp.ncbi.nlm.nih.gov/genomes/refseq/invertebrate/Folsomia_candida/representative/GCF_002217175.1_ASM221717v1/GCF_002217175.1_ASM221717v1_genomic.gff.gz | ftp://ftp.ncbi.nlm.nih.gov/genomes/refseq/invertebrate/Folsomia_candida/representative/GCF_002217175.1_ASM221717v1/GCF_002217175.1_ASM221717v1_protein.faa.gz |
| ocin | ftp://ftp.ncbi.nlm.nih.gov/genomes/genbank/invertebrate/Orchesella_cincta/representative/GCA_001718145.1_ASM171814v1/GCA_001718145.1_ASM171814v1_genomic.fna.gz | ftp://ftp.ncbi.nlm.nih.gov/genomes/genbank/invertebrate/Orchesella_cincta/representative/GCA_001718145.1_ASM171814v1/GCA_001718145.1_ASM171814v1_genomic.gff.gz | ftp://ftp.ncbi.nlm.nih.gov/genomes/genbank/invertebrate/Orchesella_cincta/representative/GCA_001718145.1_ASM171814v1/GCA_001718145.1_ASM171814v1_protein.faa.gz |
| dpul | ftp://ftp.ncbi.nlm.nih.gov/genomes/genbank/invertebrate/Daphnia_pulex/representative/GCA_000187875.1_V1.0/GCA_000187875.1_V1.0_genomic.fna.gz | ftp://ftp.ncbi.nlm.nih.gov/genomes/genbank/invertebrate/Daphnia_pulex/representative/GCA_000187875.1_V1.0/GCA_000187875.1_V1.0_genomic.gff.gz | ftp://ftp.ncbi.nlm.nih.gov/genomes/genbank/invertebrate/Daphnia_pulex/representative/GCA_000187875.1_V1.0/GCA_000187875.1_V1.0_protein.faa.gz |
| smar | ftp://ftp.ensemblgenomes.org/pub/metazoa/release-38/fasta/strigamia_maritima/dna/Strigamia_maritima.Smar1.dna.toplevel.fa.gz | ftp://ftp.ensemblgenomes.org/pub/metazoa/release-38/gff3/strigamia_maritima/Strigamia_maritima.Smar1.38.gff3.gz | ftp://ftp.ensemblgenomes.org/pub/metazoa/release-38/fasta/strigamia_maritima/pep/Strigamia_maritima.Smar1.pep.all.fa.gz |
| apis | ftp://ftp.ensemblgenomes.org/pub/metazoa/release-38/fasta/acyrthosiphon_pisum/dna/Acyrthosiphon_pisum.Acyr_2.0.dna.toplevel.fa.gz | ftp://ftp.ensemblgenomes.org/pub/metazoa/release-38/gff3/acyrthosiphon_pisum/Acyrthosiphon_pisum.Acyr_2.0.38.gff3.gz | ftp://ftp.ensemblgenomes.org/pub/metazoa/release-38/fasta/acyrthosiphon_pisum/pep/Acyrthosiphon_pisum.Acyr_2.0.pep.all.fa.gz |
| phum | https://www.vectorbase.org/download/pediculus-humanus-usdascaffoldsphumu2fagz | https://www.vectorbase.org/download/pediculus-humanus-usdabasefeaturesphumu24gff3gz | https://www.vectorbase.org/download/pediculus-humanus-usdapeptidesphumu24fagz |
| amel | ftp://ftp.ncbi.nlm.nih.gov/genomes/refseq/invertebrate/Apis_mellifera/representative/GCF_000002195.4_Amel_4.5/ | ftp://ftp.ncbi.nlm.nih.gov/genomes/refseq/invertebrate/Apis_mellifera/representative/GCF_000002195.4_Amel_4.5/ | ftp://ftp.ncbi.nlm.nih.gov/genomes/refseq/invertebrate/Apis_mellifera/representative/GCF_000002195.4_Amel_4.5/ |

| | | | |
|---|---|---|---|
| | GCF_000002195.4_Amel_4.5_genomic.fna.gz | GCF_000002195.4_Amel_4.5_genomic.gff.gz | GCF_000002195.4_Amel_4.5_protein.faa.gz |
| tcas | ftp://ftp.ncbi.nlm.nih.gov/genomes/refseq/invertebrate/Tribolium_castaneum/representative/GCF_000002335.3_Tcas5.2/GCF_000002335.3_Tcas5.2_genomic.fna.gz | ftp://ftp.ncbi.nlm.nih.gov/genomes/refseq/invertebrate/Tribolium_castaneum/representative/GCF_000002335.3_Tcas5.2/GCF_000002335.3_Tcas5.2_genomic.gff.gz | ftp://ftp.ncbi.nlm.nih.gov/genomes/refseq/invertebrate/Tribolium_castaneum/representative/GCF_000002335.3_Tcas5.2/GCF_000002335.3_Tcas5.2_protein.faa.gz |
| dple | ftp://ftp.ensemblgenomes.org/pub/metazoa/release-38/fasta/danaus_plexippus/dna/Danaus_plexippus.Dpv3.dna.toplevel.fa.gz | ftp://ftp.ensemblgenomes.org/pub/metazoa/release-38/gff3/danaus_plexippus/Danaus_plexippus.Dpv3.38.gff3.gz | ftp://ftp.ensemblgenomes.org/pub/metazoa/release-38/fasta/danaus_plexippus/pep/Danaus_plexippus.Dpv3.pep.all.fa.gz |
| dmel | ftp://ftp.flybase.net/genomes/Drosophila_melanogaster/current/fasta/dmel-all-chromosome-r6.22.fasta.gz<br><br>ftp://ftp.ncbi.nlm.nih.gov/genomes/refseq/invertebrate/Drosophila_melanogaster/reference/GCF_000001215.4_Release_6_plus_ISO1_MT/GCF_000001215.4_Release_6_plus_ISO1_MT_genomic.fna.gz | ftp://ftp.flybase.net/genomes/Drosophila_melanogaster/current/gff/dmel-all-filtered-r6.22.gff.gz<br><br>ftp://ftp.ncbi.nlm.nih.gov/genomes/refseq/invertebrate/Drosophila_melanogaster/reference/GCF_000001215.4_Release_6_plus_ISO1_MT/GCF_000001215.4_Release_6_plus_ISO1_MT_genomic.gff.gz | ftp://ftp.flybase.net/genomes/Drosophila_melanogaster/current/fasta/dmel-all-translation-r6.22.fasta.gz<br><br>ftp://ftp.ncbi.nlm.nih.gov/genomes/refseq/invertebrate/Drosophila_melanogaster/reference/GCF_000001215.4_Release_6_plus_ISO1_MT/GCF_000001215.4_Release_6_plus_ISO1_MT_protein.faa.gz |
| lmig | ftp://ftp.ncbi.nlm.nih.gov/genomes/genbank/invertebrate/Locusta_migratoria/representative/GCA_000516895.1_LocustGenomeV1/GCA_000516895.1_LocustGenomeV1_genomic.fna.gz | | |
| bger | ftp://ftp.ncbi.nlm.nih.gov/genomes/genbank/invertebrate/Blattella_germanica/representative/GCA_003018175.1_Bger_1.1/GCA_003018175.1_Bger_1.1_genomic.fna.gz | | |
| choo | ftp://ftp.ncbi.nlm.nih.gov/genomes/genbank/invertebrate/Clitarchus_hookeri/representative/GCA_002778355.1_ASM277835v1/GCA_002778355.1_ASM277835v1_genomic.fna.gz | | |
| mext | ftp://ftp.ncbi.nlm.nih.gov/genomes/genbank/invertebrate/Medauroidea_extradentata/representative/GCA_003012365.1_ASM301236v1/GCA_003012365.1_ASM301236v1_genomic.fna.gz | | |

### SUPPLEMENTARY FIGURES

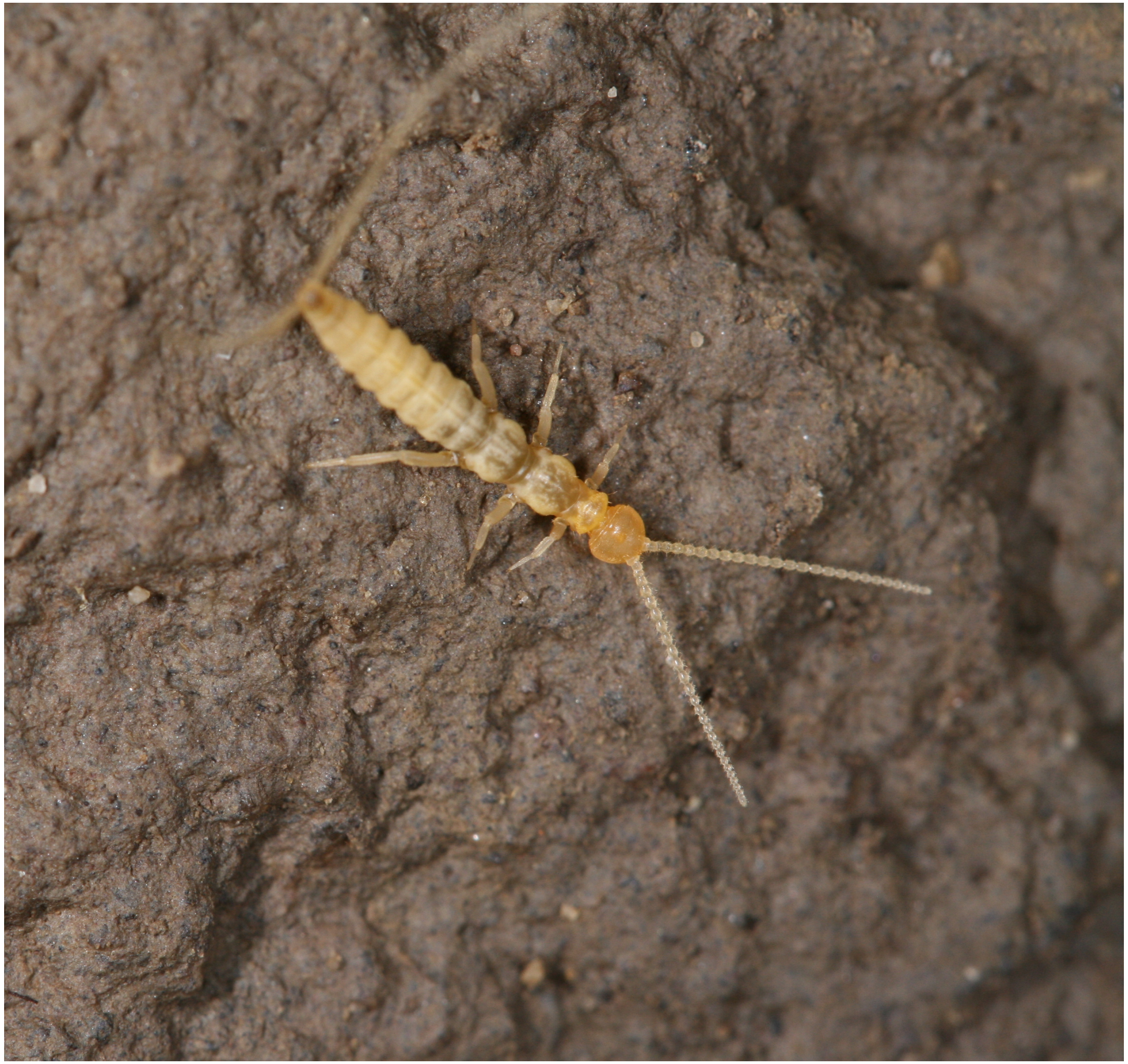

**Figure S1a:** High-resolution picture of *C. augens*.

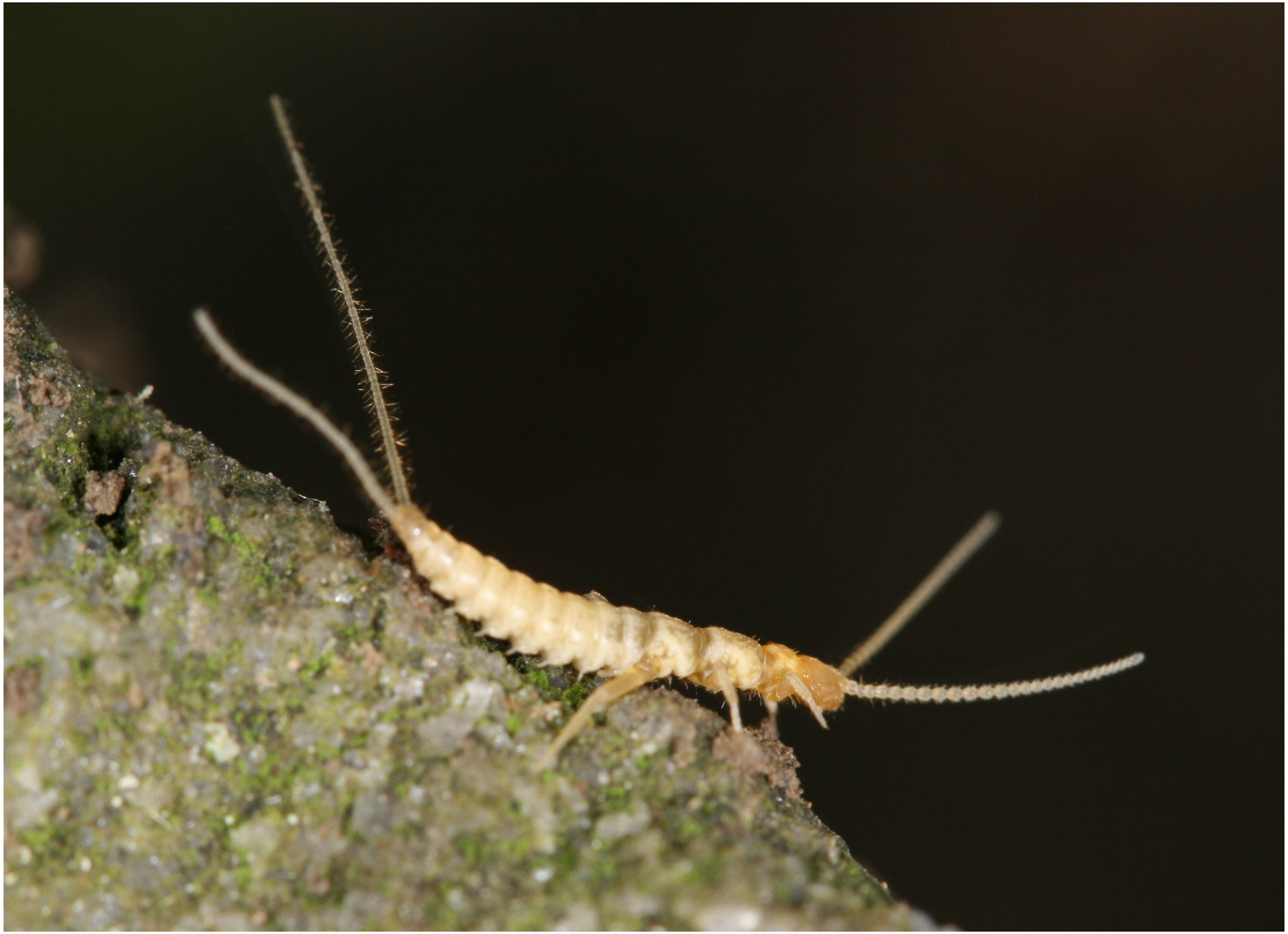

**Figure S1b:** High-resolution picture of *C. augens*.

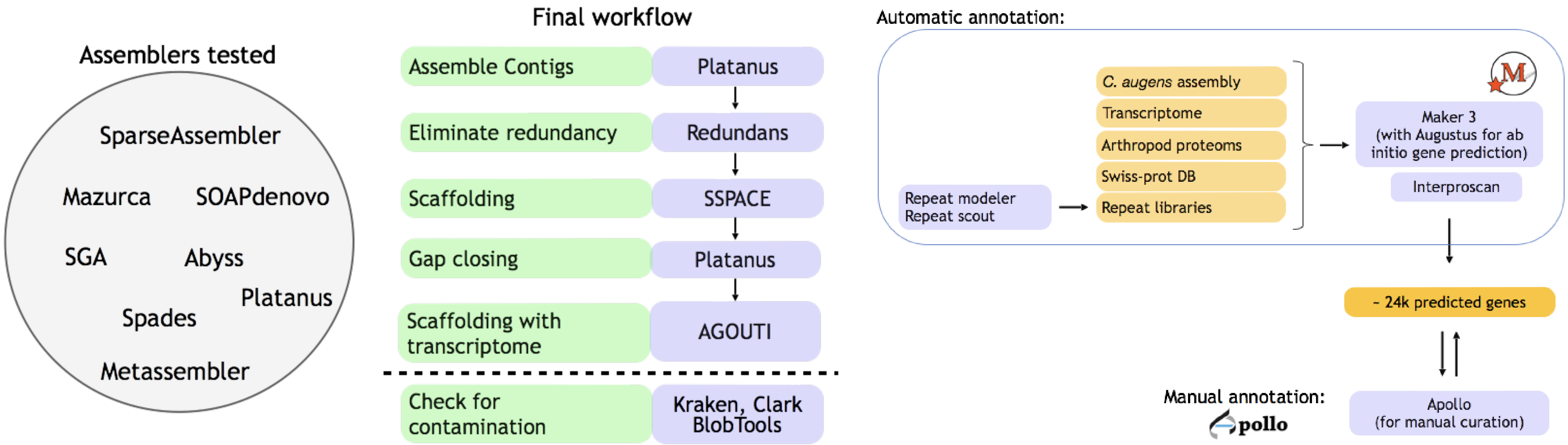

**Figure S2:** Assembly and annotation workflow used for *C. augens*.

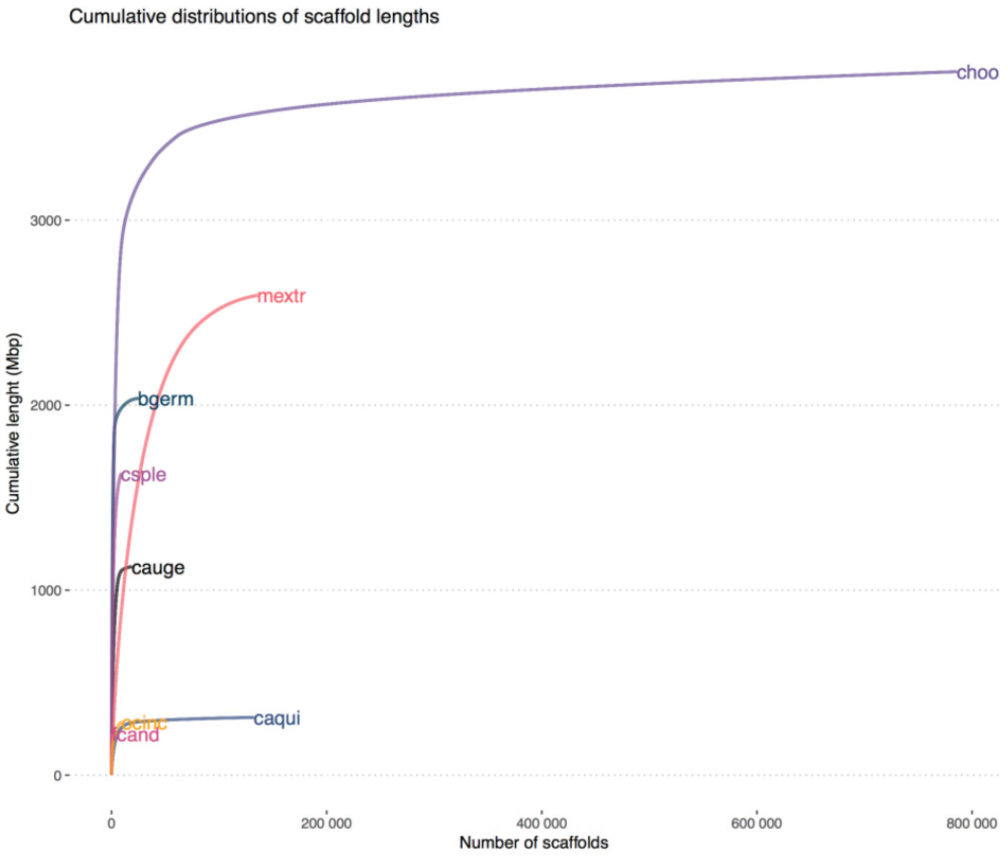

**Figure S3:** Cumulative distributions of scaffold lengths in *C. augens* and other arthropods. Csple, *Calopteryx splendens* (banded demoiselle); Caqui, *Catajapyx aquilonaris* (northern forcepstail); Fcand, *Folsomia candida* (springtail); Ocinc, *Orchesella cincta* (springtail); Bgerm, *Blattella germanica*; Chook, *Clitarchus hookeri*; Mextr, *Medauroidea extradentata*.

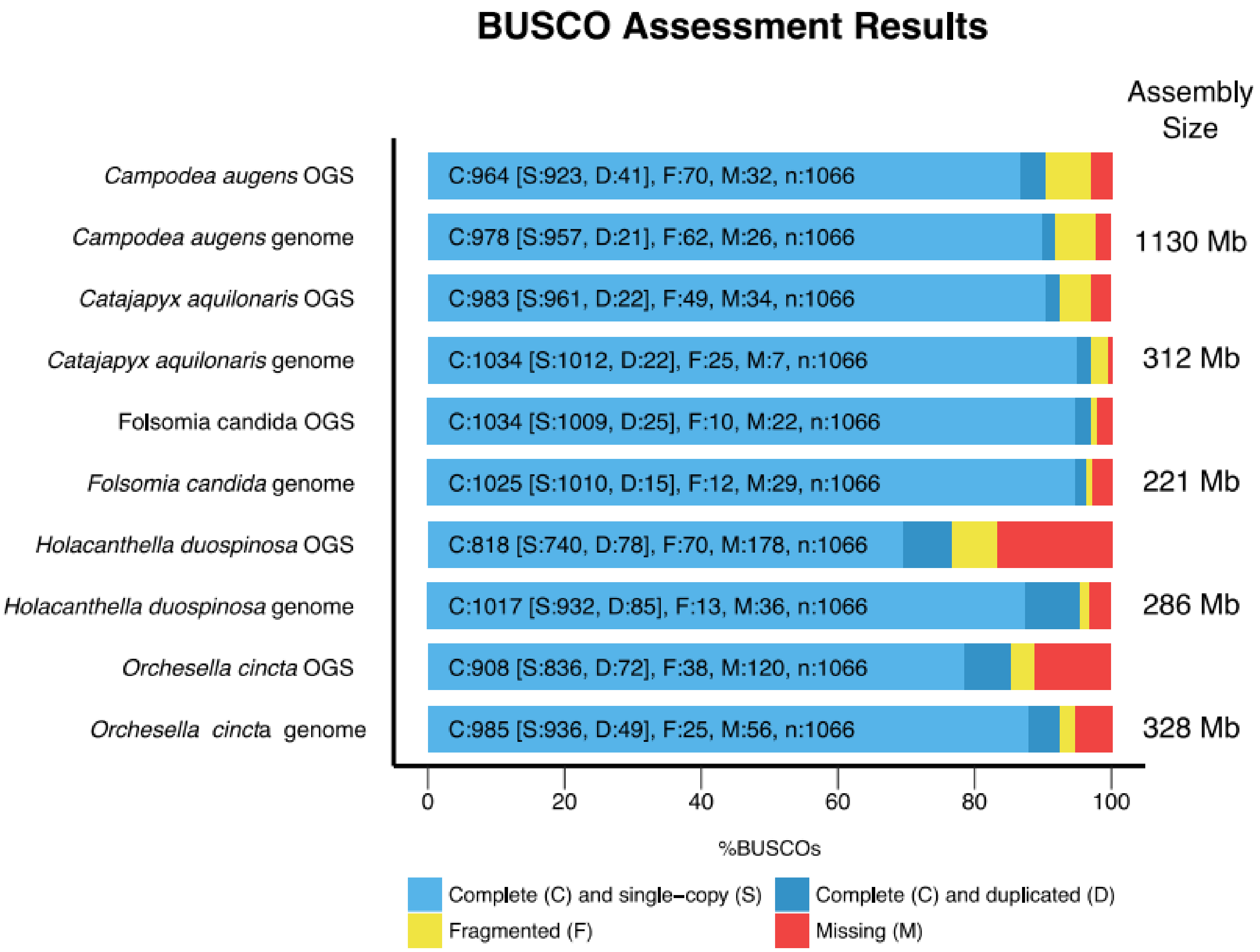

**Figure S4:** Plots of BUSCO scores for *C. augens* and the other ancestrally wingless hexapods using the arthropoda_v9 dataset.

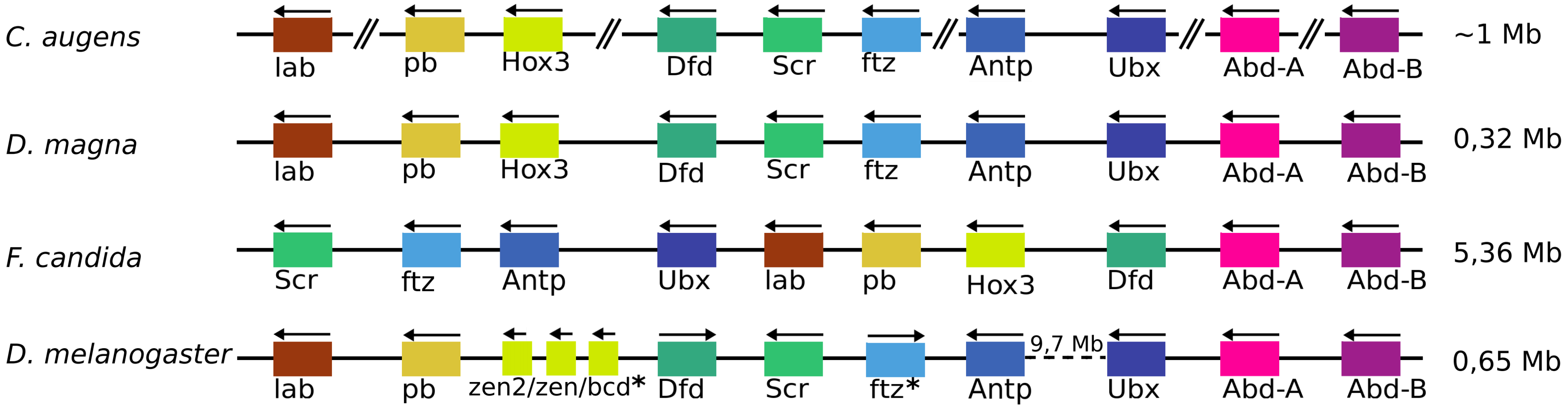

**Figure S5:** Hox genes cluster arrangement for *Campodea augens, Daphnia magna, Folsomia candida* and *Drosophila melanogaster*.
Double slash delimits different scaffolds. Dotted line indicates a large genomic region of ca. 10 Mb which characterizes the hox genes cluster of *D. melanogaster*.
(*) in *D. melanogaster* genes indicates genes that do not have homeotic function (Hughes & Kaufman 2002).

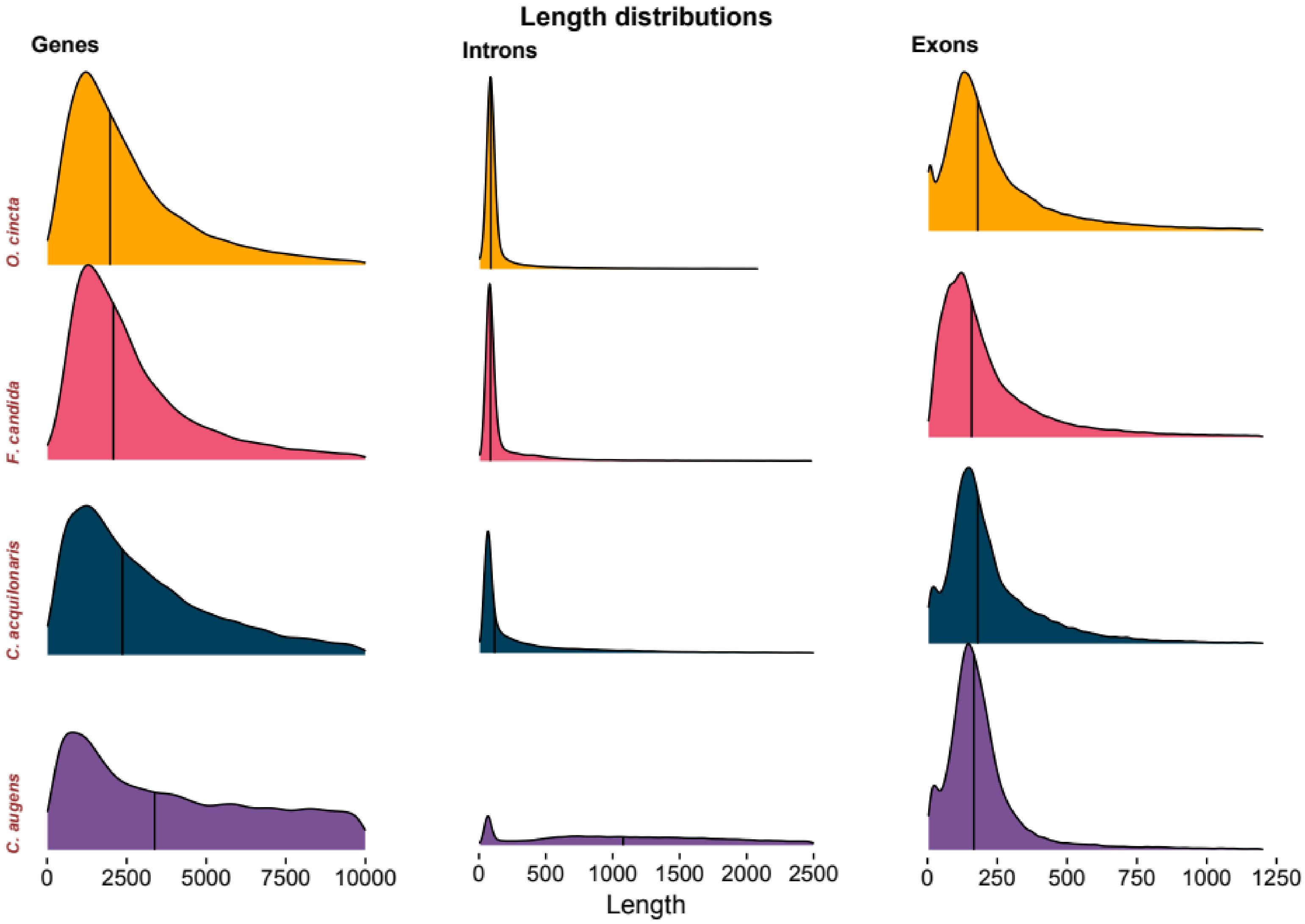

**Figure S6:** Gene, intron and exon length distributions in *C. augens* and other ancestrally wingless hexapods.

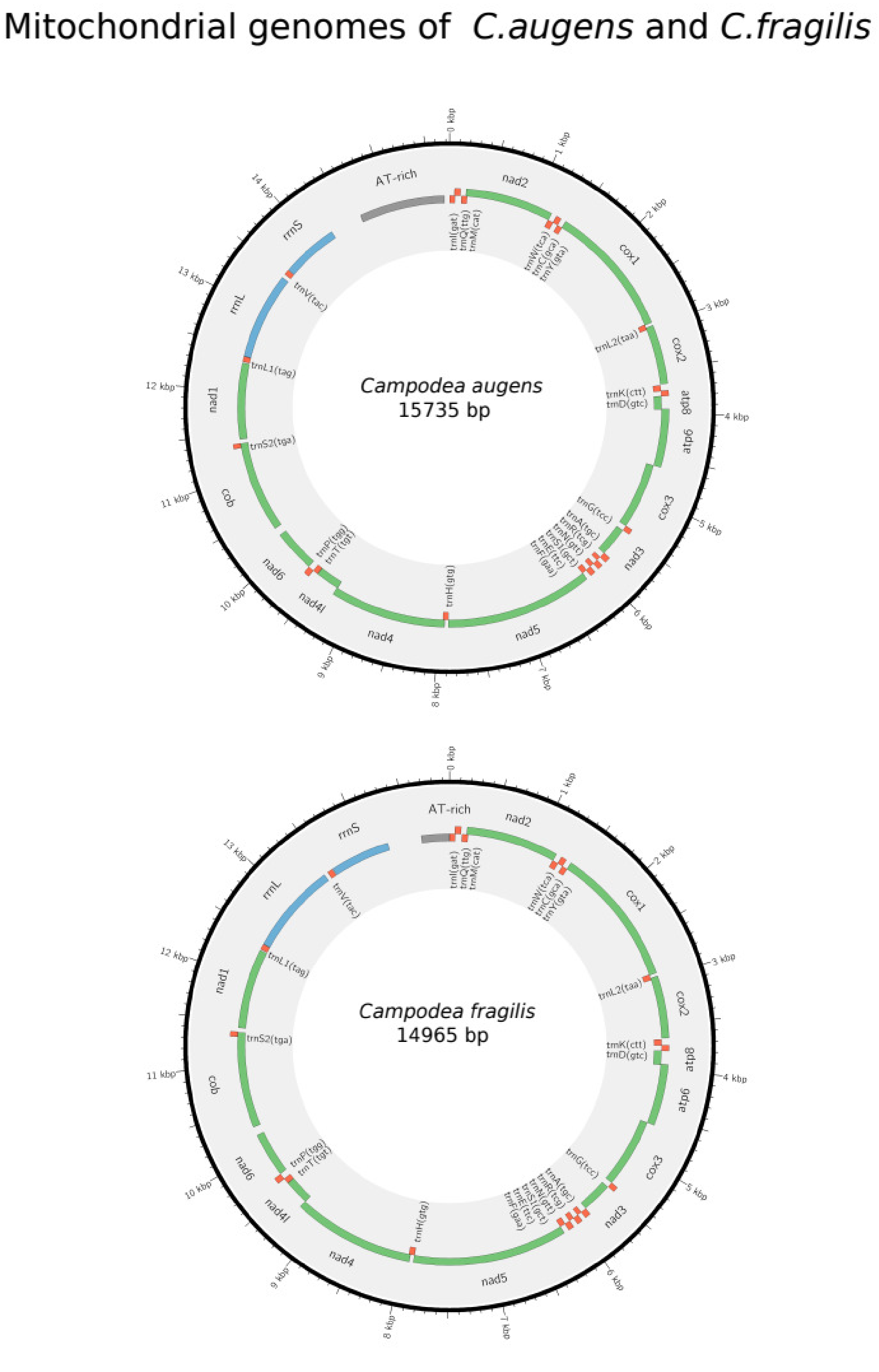

**Figure S7:** Graphical representation of the mitochondrial genomes of *C. augens* and *C. fragilis.*

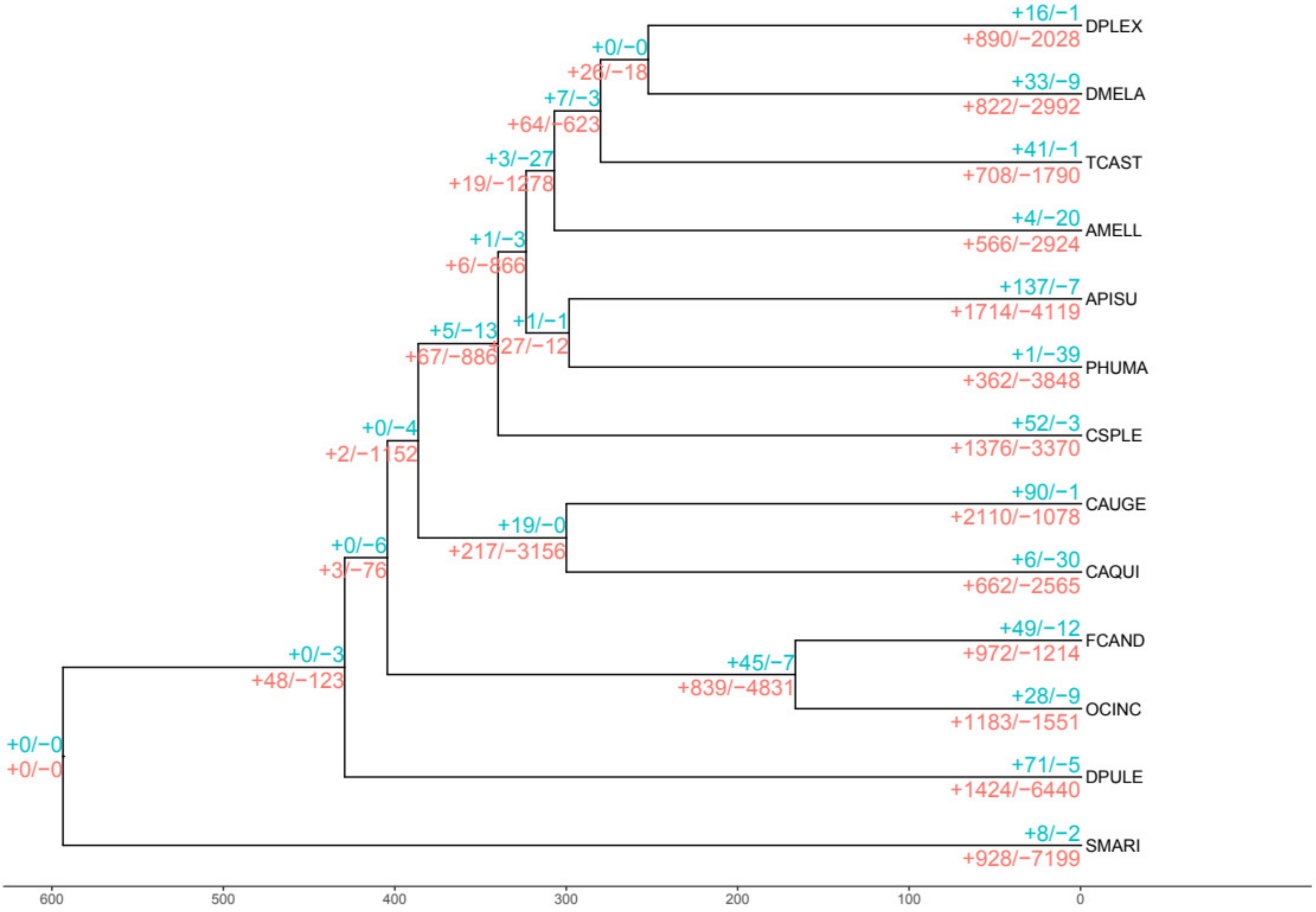

**Figure S8:** Ultrametric species phylogeny displaying expansions/contractions of gene families as estimated using CAFE. Values in green correspond to the number of significantly (Viterbi p-value < 0.01) expanded (+) and contracted (-) gene families. Values in red correspond to the total number of expansions/contractions including the non-significant ones, and correspond to the values represented in the pie charts of Figure 2.

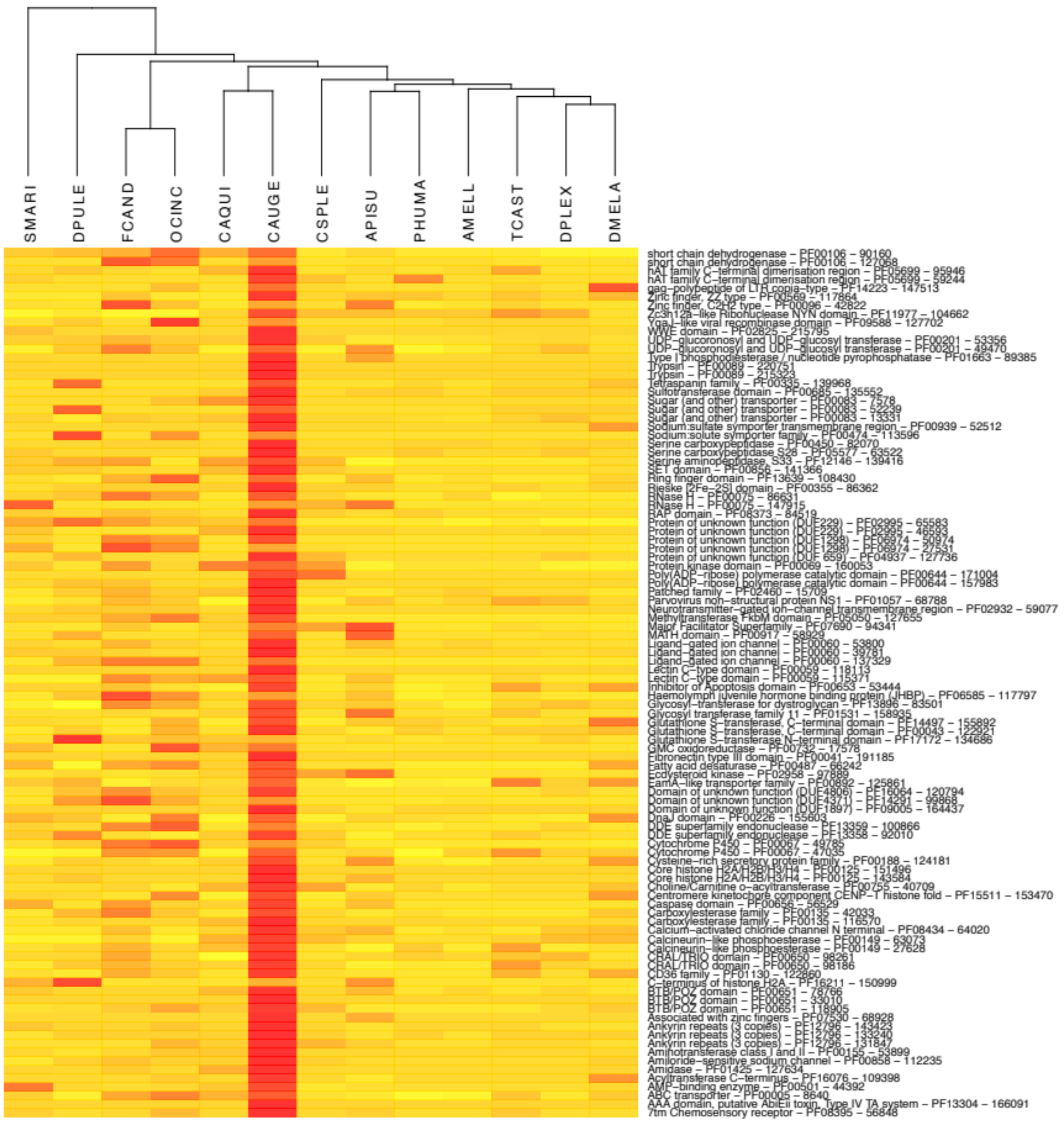

**Figure S9:** Heat map of OrthoDB clusters that are significantly larger in *C. augens* compared to the mean counts from insect species. The most representative pfam domain for each cluster is reported on the right.

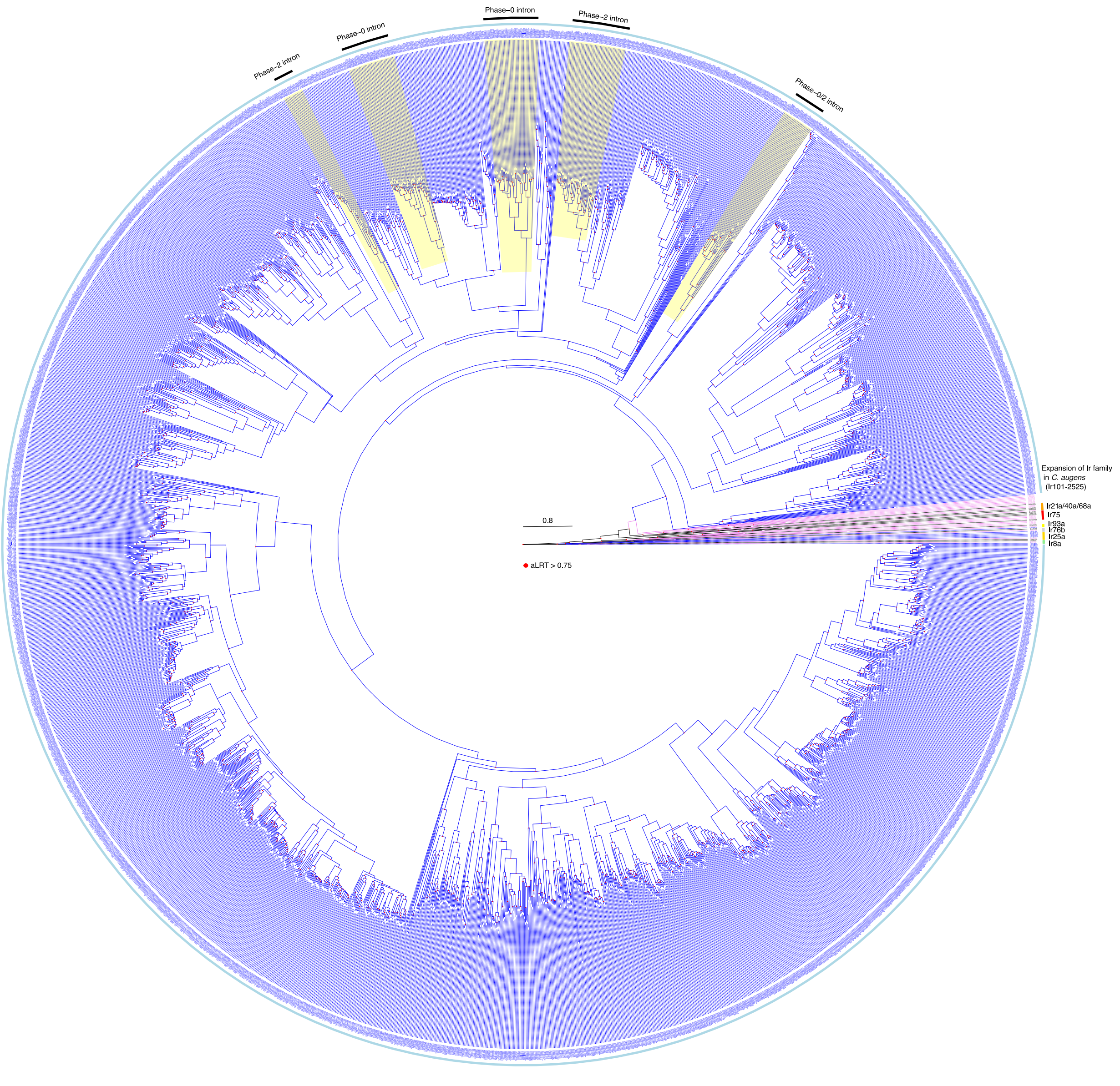

**Figure S10:** Phylogenetic tree of the IR family. This tree was rooted by declaring the Ir8a and 25a lineages as the outgroup, based on their basal positions within larger trees including the ionotropic glutamate receptors from which the IRs evolved. The *Campodea augens* (Caug) proteins are in blue, the *Drosophila melanogaster* (Dmel) proteins for the seven conserved IRs with orthologs in *C. augens*, as well as the Ir75 clade, are colored black, while the *Calopteryx splendens* (Cspl) proteins are colored purple. The seven conserved lineages are highlight in colors and indicated outside the circle. The five CaugIr lineages that have idiosyncratically gained introns are highlighted in yellow, with the intron phase(s) indicated outside the circle, showing the independence of these five lineages whose introns are all in different locations. The scale bar indicates substitutions per site. Red filled circles indicate nodes with an approximate Likelihood-Ratio Test (aLRT) > 0.75. Zoom in to view the details and names of proteins.

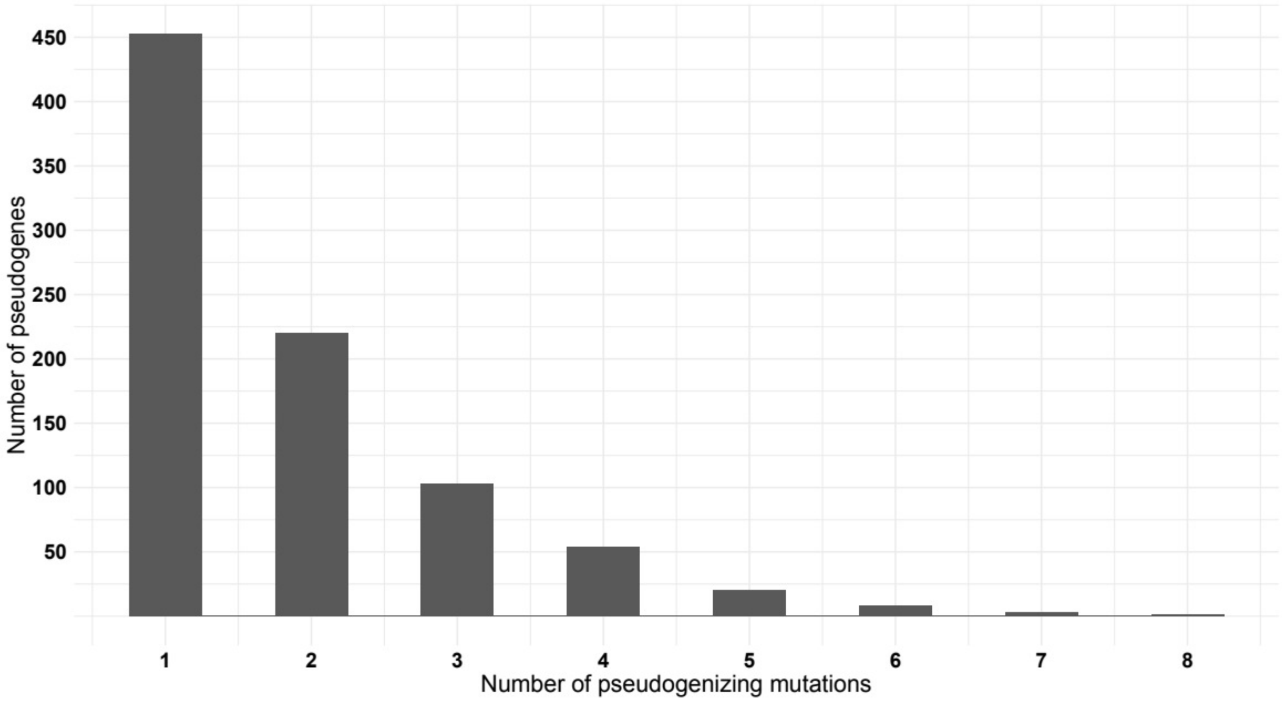

**Figure S11:** Histogram of the numbers of pseudogenes with 1-8 pseudogenizing mutations.

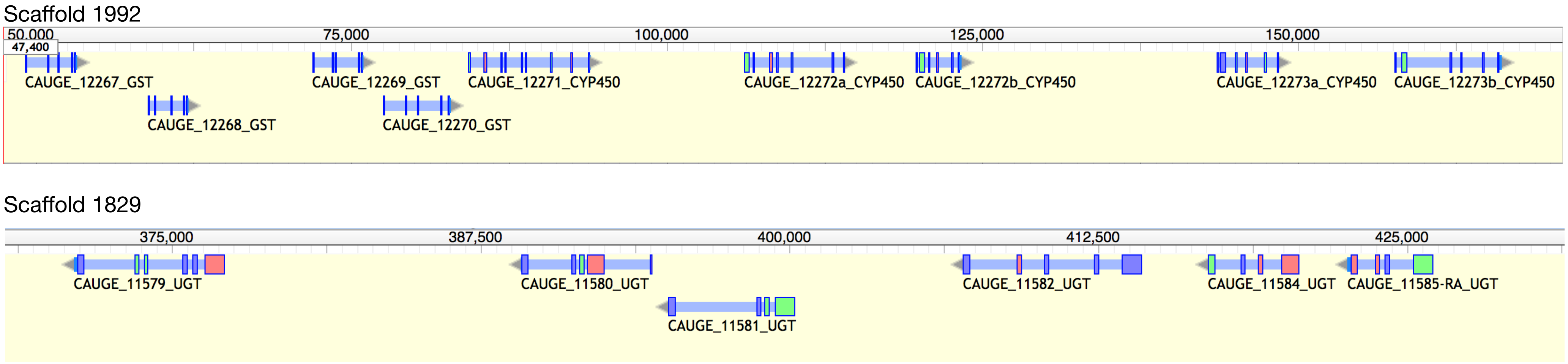

**Figure S12:** Genomic regions encoding a cluster of 5 CYP450 with 4 GST genes (Scaffold_1992), and a cluster of 6 UGTs (Scaffold_1829). Large boxes correspond to exons.

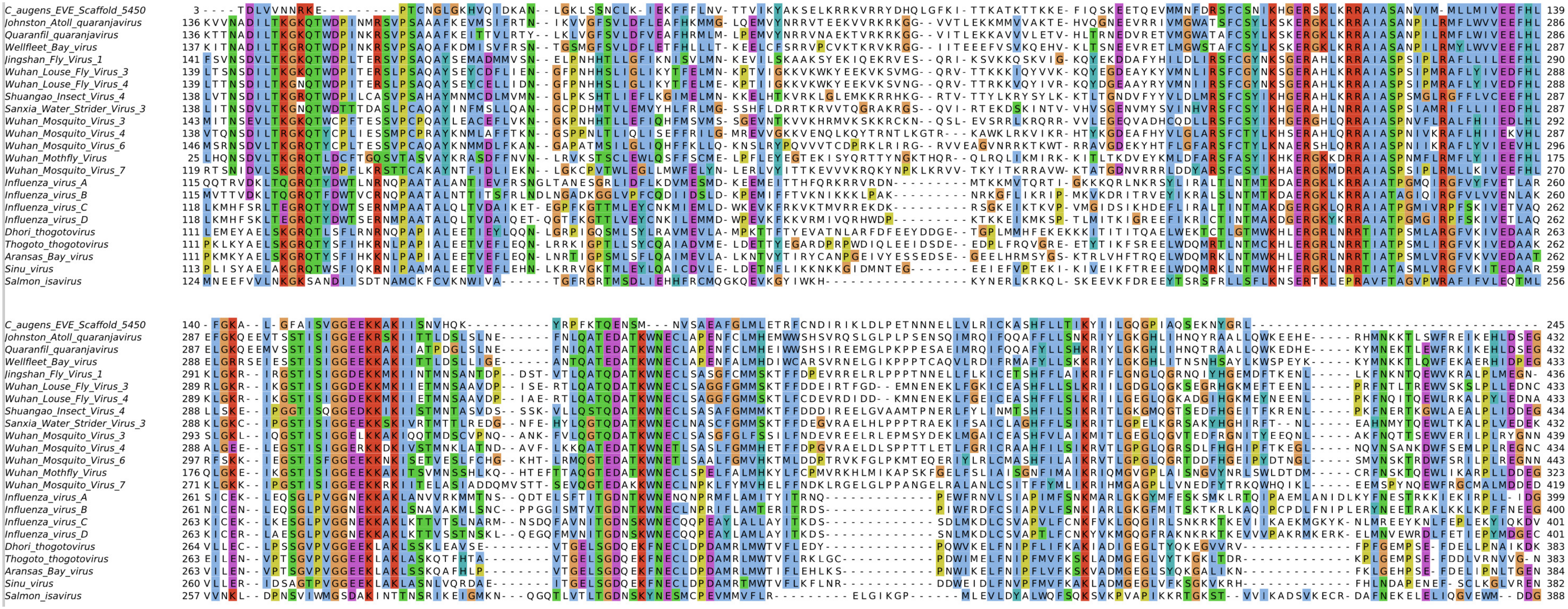

**Figure S13:** MSA of a putative endogenous viral elements in *C. augens* genome (C_augens_EVE_5450) and PB1 sequences of Orthomyxoviridae.

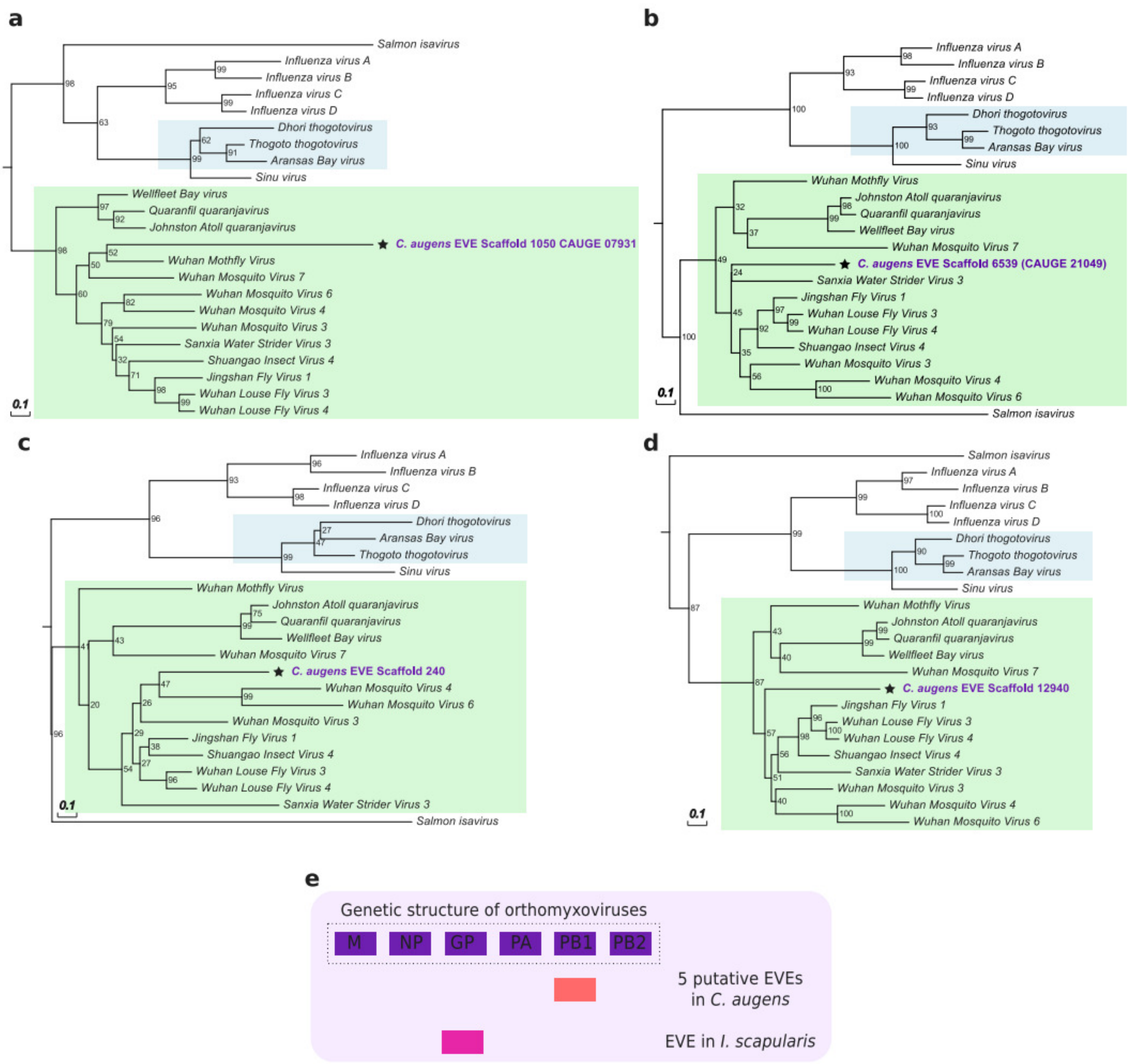

**Figure S14:** Phylogenetic relationships of the EVEs found in *C. augens* related to -ssRNA viruses of the Orthomyxoviridae family. **a-d**, All EVEs in *C. augens* correspond to orthomyxoviral Polymerase Basic protein 1 (PB1) (Pfam Id PF00602; "Flu_PB1"). Neighbor-joining trees were constructed using alignments of EVE amino acid sequences with BP1 proteins of representatives of the Orthomyxoviridae family (see Supplementary Table 14 for GenBank accession nos.). Support for trees was evaluated using 1,000 pseudo replicates. Values correspond to the bootstrap support values. Scale bars indicate amino acid substitutions per site. Green boxes highlight viruses of the Quaranjavirus genus. **e**, Comparison of EVEs related to quaranjaviruses found in *C. augens* and the tick *Ixodes scapularis* (Katzourakis and Gifford, 2010). While all EVEs in *C. augens* correspond to PB1, EVE in *I. scapularis* corresponds to the glycoprotein (GP) of quaranjaviruses.
